# Reef Cover: a coral reef classification to guide global habitat mapping from remote sensing

**DOI:** 10.1101/2020.09.10.292243

**Authors:** Emma Kennedy, Chris Roelfsema, Mitchell Lyons, Eva Kovacs, Rodney Borrego-Acevedo, Meredith Roe, Stuart Phinn, Kirk Larsen, Nicholas Murray, Doddy Yuwono, Jeremy Wolff, Paul Tudman

## Abstract

Coral reef management and conservation stand to benefit from improved high-resolution global mapping. Yet classifications employed in large-scale reef mapping to date are typically poorly defined, not shared or region-specific. Here we present *Reef Cover*, a new coral reef geomorphic zone classification, developed to support global-scale coral reef habitat mapping in a transparent and version-based framework. We developed scalable classes by focusing on attributes that can be observed remotely, but whose membership rules also reflect knowledge of reef formation, growth and functioning. Bridging the divide between earth observation data and geo-ecological knowledge of reefs, *Reef Cover* maximises the trade-off between applicability at global scales, and relevance and accuracy at local scales. We use the Caroline and Mariana Island chains in the Pacific as a case study to demonstrate use of the classification scheme and its scientific and conservation applications. The primary application of *Reef Cov*er is the *Allen Coral Atlas* global coral reef mapping project, but the system will support bespoke reef mapping conducted at a variety of spatial scales.

## 1. Background and Summary

Enhanced earth observation and analytical capabilities have revolutionised the way we view our planet, allowing a global perspective that has the potential to dramatically influence the way humanity manages finite planetary resources^1^. Yet for coral reefs, a valuable and rapidly degrading ecosystem^2,3^, these accelerating capabilities to collect and analyse data are not often translated into improved environmental outcomes^4^. One barrier is the transfer of insights gained from remote sensing data from producers (remote sensing scientists) to end-users. Translation of remotely collected observations of coral reefs into user-friendly spatial information for investigation, monitoring, planning and management requires a classification system to discretise continuous data about natural phenomena into spatial units of manageable information.

Classification is an important process for making data available to people that need it^5^. Maps are an efficient way to share large amounts of spatial data, but to be effective any classification needs to a) follow a clear and transparent rationale for developing thematic map classes, b) develop classes that are meaningful, clearly described and communicated and c) be accessible to the people that utilise them. The latter is particularly important if information is to be successfully distributed and meaningfully integrated into practical solutions founded on map data^5^. Here we present the *Reef Cover* classification, consisting of 17 shallow tropical coral reef internal geomorphic class descriptors, developed specifically for global reef mapping to support reef science and conservation.

Classification of coral reef geomorphology has typically occurred using two broad approaches, which are largely congruent^6^, but differ considerably by the disciplinary approach, methodology employed, datasets used, and the scale of investigation. For centuries, reef classification has involved an *a priori* grouping of natural features into classes using a combination of detailed field observations and expert knowledge, often drawing from multiple disciplinary fields, and making use of diverse ecological and geological datasets (e.g. drill cores, bathymetry readings, ecological benthic data) and natural history theory (*Expert-led* approach, Fig. 1). These natural history classifications draw from a breadth of understanding of reef genesis and history, but lack a standardised approach (e.g. compared to highly regulated knowledge structures such as taxonomic classification of organisms, linguistics or computer science^7^), due to difficulties of integrating diverse information sources and complex knowledge. Geographic, linguistic and disciplinary differences have sometimes caused lack of agreement on a) nomenclature (terminology of reef features), b) structure (how some of these features are grouped) and c) meaning (how they are interpreted and relevant and how useful they may be for practical research). For example, the term *Back Reef* can be interpreted in many ways^8^, and may be understood differently by an ecologist vs a geologist, or a Caribbean vs a Pacific- based scientist.

**Figure 1.**
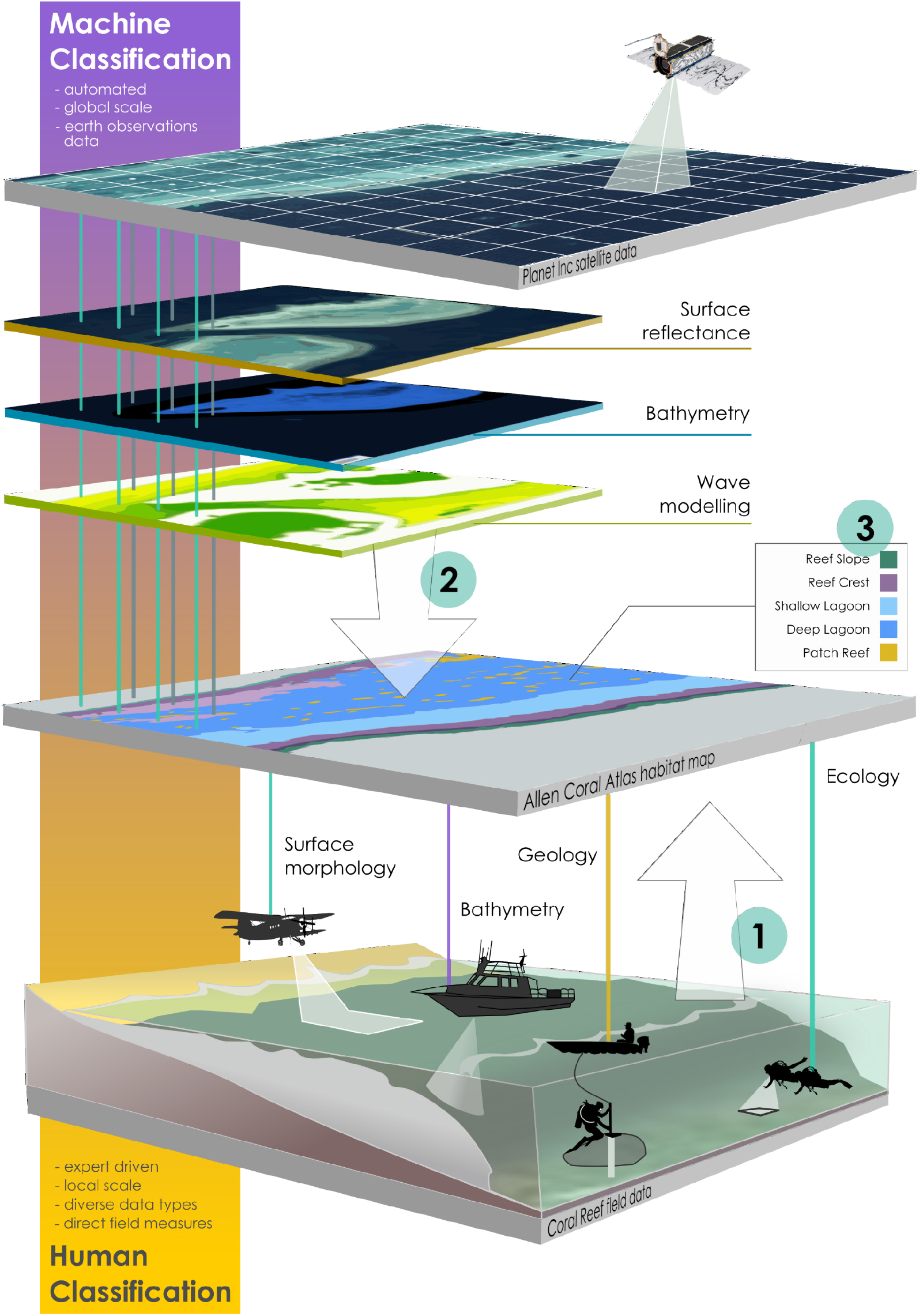
Different scales (global vs local) and scientific approaches (direct field measurements vs remote satellite observations) for capturing natural variability of morphological reef features can shape our understanding of how reefs are structured. Coral reef habitat maps, like the *Allen Coral Atlas*, NOAA’s *Biogeography of Coral Reef* maps, *Living Oceans Foundation* and *Millennium Coral Reef Mapping Project*, aim to distil the vibrant natural diversity of coral reef ecosystems into clustered information that is relevant, interpretable and useful to humans, so regional-to-global scale patterns of variability can be widely disseminated. A classification system for coral reef zones should try to integrate (1) decades of local scientific knowledge on coral reef systems, with the (2) global-scale information that is becoming more accessible from satellite sensors such as those operated by Planet Inc. and information derived from these products, to (3) generate map classes that can inform understanding or management of coral reefs.

In the last 40 years, however, technological advancements in the field of remote sensing have led to a different approach to reef classification. Specifically for scaling habitat mapping from remote sensing, this largely computational approach (*Machine-led* approach, Fig. 1), involves the *a posteriori* seascape-scale grouping of pixels into classes, often using probability-based sampling of a one or two broad-coverage remote sensing datasets (e.g. spectral reflectance, bathymetry). Classification methods developed by remote sensing scientists has driven rapid expansion of reef mapping efforts from reef-scale to ocean-scale extents^9^. However, these approaches are known to be limited by the spatial, spectral, radiometric and temporal resolutions of source data used in the classifications^10^, and sometimes do not adequately consider the wealth of natural history, existing “traditional” coral reef classifications and associated terminology, or the requirements of users.

Three challenges related to remote sensing classifications can hamper effective use of new generation coral reef maps. The first challenge relates to divergence in how a natural scientist or manager understands a complex system, and how that system can be characterized by biophysical data layers (e.g. satellite imagery, physical and environmental data layers).

Secondly, a lack of transparency about classification processes may confound interpretation of classes by users. A recent study showed that just 32% of publications provided sufficient information to replicate a series of mapping studies (99/304 published studies reviewed^11^).

The third challenge is communication: map classes are frequently not accessible to users, or not sufficiently defined or described. A recent review found that only 52% of satellite-derived coral reef and seagrass habitat maps were accompanied by a detailed class descriptor that would allow users to interpret mapped classes (49/72 published coral reef and seagrass habitat maps^12^).

To address these challenges, we have developed *Reef Cover*, a classification system that aims to bridge disciplinary gaps by focussing on attributes of reef features that can be mapped from most remotely-sensed data (Methods: Step 2) (e.g., nanosatellites, Fig. 1), while simultaneously aligning with foundational geo-ecological understanding of reef formation and growth (Methods: Step 1)^13^. Class definitions were developed with users in mind (Methods: Step 3), and accessibility issues were addressed by making *Reef Cover*, and the maps that have been derived with *Reef Cover* classes, open access and freely available.

## 2. Methodology

The *Reef Cover* classification dataset presented defines a set of 17 classes that are transferable between two domains: the traditional ecological-biophysical perspective and the earth observations systems view of reefs (Fig. 1). The typology acts as a key to bridge historic and contemporary knowledge, plot-scale and aerial viewpoints, and pixel data with natural history to convert pixel data into information in a form suitable for reef management decisions.

*Reef Cover* was specifically developed to support the process used to produce and deliver a global set of coral reef mapping and field survey products from remotely sensed data^14^. Accompanying methods^15^ and datasets^16,90^ to aid use are also publically available. These products were developed specifically to support science and conservation of coral reef ecosystems.

We sought to develop a parsimonious system which balances the geomorphic complexity of reefs with the need to develop high accuracy maps of each class in the system. The result is a 17 class system that can be (i) applied to remote sensing datasets for future mapping, (ii) used to interpret maps (iii) effectively disseminated to users – mainly in coral reef ecology and conservation space – in a way that promotes use in research and conservation.

Three steps were used in development of the classification.

Step 1. **Review**. Existing coral reef geomorphic classification schemes related to different sources and user cases were reviewed, specifically to examine terminology and class groupings and how they can be described in terms of biophysical data related to remote sensing. This allowed us to select terminology and groupings for the classification that build constructively on previous foundational knowledge on coral reef geomorphology and can be related to existing mapping efforts.

Step 2. **Development**. Physical attributes datasets available to remote sensing scientists were examined in order to create a set of 17 meaningful internal reef classes that relate to broader interpretation from a natural history point of view, gathered in Step 1. Methodology behind generating *Reef Cover* classes from attributes data builds on the reef mapping theory presented in Roelfsema *et al* ^9^.

Step 3. **Dissemination**. *Reef Cover* classes were drafted in a way to promote re-use and cross-walking, with a strong focus on needs of the users (*Reef Cover* dataset). This included consideration of 1) *relevance* e.g. rationale behind why it was important to map this class, but also broader global applicability of the class, *2) simplicity* e.g., promoting user-uptake by employing plain language, not over-complicating descriptors and limiting the number of classes to manageable amount, *3) transparency* supplying methodological basis behind each class, and exploring caveats and ambiguities in interpretation, 4) *accessibility* including discoverability, open access and language translations to support users, and 5) *flexibility* allowing for flexible use of the scheme depending on user needs, allowing for flexible interpretation of classes by providing cross-walk to other schemes and existing maps, and making the classification adaptable, and open to user feedback.

Finally, the *Reef Cover* classification was tested by applying it to a mapping exercise in Micronesia^90^ (Technical Validation section). During this process the *Reef Cover* dataset was reviewed to explore how useful it was for a) producers using *Reef Cover* to map large coral reef areas from satellite data, and b) consumers using *Reef Cover* to interpret map products for application to real world problems.

### 2.1 Step 1. Review. Building global classes and terminology on foundational reef mapping and classification work

#### 2.1.1. Global reef mapping: the need for a classification to map coral reefs at scale

Coral reefs represent pockets of biodiversity that are widely dispersed, often remote/inaccessible and globally threatened^2,3^. Communities and economies are highly dependent on the ecosystem services they provide^17-19^. This combination of vulnerability, value and a broad and dispersed global distribution mean global strategies are needed for reef conservation, for which maps (and the classifications that underpin them) play a supporting role. Global coral reef maps have been fundamental to geo-political resource mapping and understanding inequalities^20,21^, the valuation of reef ecosystem services^18^, understanding the past^22^, present^23^, and future threats to reefs^24^, supporting more effective conservation^25,26^ and reef restoration strategies^27,28^, and facilitating scientific collaborations and research outcomes^29^. Reef conservation science and practice may particularly benefit from technological advancements that allow delivery of more appropriate map-based information, particularly across broader, more detailed spatial scales and in a consistent manner^14,26,28^.

Shallow water tropical coral reefs are particularly amenable to global mapping from above^30^. They develop in clear, oligotrophic tropical waters, so many features are detectable from space^31^. Remote sensing scientists have been developing automated methods to make sense of the increasing availability of earth observation data over coral reefs, yielding information on ecosystem zones derived from data sources such as spectral reflectance and bathymetry at increasingly larger scales^30,32^. As more data reveal the diversity and complexity of reefs, selecting an appropriate level at which to map reefs on the global scale requires balancing the need for a limited number of classes that can be mapped consistently based on available earth observation data, with user need for information.

#### 2.1.2. Scaling and consistency: why use geomorphic zones as classes?

##### Reef type classification

Morphological diversity can make global geomorphic classification – particularly between reefs (at the “reef type” level, e.g. fringing, atoll reefs) - challenging. Divergent regional morphologies (e.g. Pacific atolls *vs* Caribbean fringing reefs) and endemic local features (e.g. Bahamian shallow carbonate banks, Maldivian farus) are driven by underlying tectonics, antecedent topography, eustatics, climate and reef accretion rates which can all vary geographically^33^. This diversity is reflected in the large number of map classes in the impressive *Millennium Coral Reef Mapping Project* (68 classes at the between-reef geomorphic level L3), the most comprehensive globally applicable coral reef classification system to date^34^.

##### Geomorphic zone classification

Internally, reef morphology becomes a lot more consistent. Physical boundaries in the depth, slope angle and exposure of the reef surface help create stark partitioning into “geomorphic zones” (e.g. reef flat, reef crest), developed in parallel to the reef edge and coastlines and generally with a distinct ecology^13,35,36^. These internal patterns of three-dimensional geomorphic structure and associated partitioning of organisms across that structure can be remarkably predictable, even between oceans. This makes geomorphic zonation a good basis for consistent and comparable mapping at regional to global scales^20,37^. Moreover, overlap between geomorphic zones and ecological partitioning means that ecological understanding can be implied from geomorphic habitat classes, making geomorphic mapping useful to conservation practitioners^38^.

##### Benthic classification

Many classifications developed for reef mapping (e.g. *Living Oceans Foundation*^39^ *NOAA Biogeography Reef Mapping Program*^*40*^), monitoring (*Atlantic and Gulf Rapid Reef Assessment*^*41*^, *Reef Cover Classification System*^*42*^) and management (*Marine Ecosystem Benthic Classification*^*43*^) have included an ecological component. Classifying reef benthos is important as associated metrics, such as abundance of living coral and algae, are widely-used indicators of ecosystem change. However, most classifications that consider benthic cover are operational at reef^44^ to regional scales, due to the need for very high resolution remote sensing data^45^ (e.g. from UAVs and CASI^46^, or high resolution satellites like QuickBird and WorldView 1m) to be able to reliably determine classes such as coral cover and type, soft coral, turf, coralline algae, rubble and sand. Remotely-sensed data sources available with full global coverage of coral reef benthos on daily timescales still have 3.7 m pixel resolution (Fig. 1, *Machine-led* classification), while ecological classifications generally begin at the metre (quadrat) scale (Fig. 1, *Human*-classification). This spatial mismatch meant that at the present time it was not possible to develop a comprehensive benthic coral reef classification^47^ that met the *Reef Cover* objective of being globally scalable (both in terms of remote sensing biophysical data availability and processing capabilities) but that also fully recognises and includes the rich benthic detail required to address ecological questions at sub-metre scales. Benthic mapping by the Allen Coral Atlas is determined by whats possible from remote sensing, and not how scientists view the reef (Fig 1). However in the coming years it is likely that further advancements in technology – both downscaling of remote-sensing and up-scaling of field observations^48^ -will enable us to address this spatial mismatch. At that point *Reef Cover* can be updated with a full benthic classification that meets the objectives of better matching remote-sensing data producers with user needs to support real-world outcomes.

#### 2.1.3. Previous field and ecologically driven reef classifications

Traditionally, coral reef features have been grouped and mapped based on observations of morphological structure, distributions of biota and theories on genesis, gleaned from aerial surveys, bathymetric surveys, geological cores and biological field censuses by natural scientists^36^ (Fig. 1). Natural scientists were struck by both the uniformity and predictability of much of the large-scale three-dimensional geomorphic structure of reefs and biological partitioning across that structure, and how consistent these characteristic geologic and ecological zones were across large biogeographic regions^49,50^). Technological developments of the 20^th^ century, such as SCUBA demand regulators and compressed air tanks (commercially available in the 1940s^51^), acoustic imaging for determining seafloor bathymetry (e.g. side-scan sonar developed in the 1950s), light aircraft for aerial photography (first applied in the 1950s^52^) and lightweight submersible drilling rigs for coring (applied in the 1970s^53^), allowed reef structure to be viewed from fresh perspectives. New aerial, underwater and internal assessments of reef structure expanded the diversity of external and internal classes, with thousands of new terms for features defined^8^. However, the localised nature of most of these applications (Fig. 1) meant that many of the classes developed using these tools were region-specific, leading to Stoddart et al (1978) warning against too heavy a reliance on the “the imperfect and perhaps biased existing field knowledge on reefs” for developing global classifications.

One of the first steps in creating the *Reef Cover* classification was reconciling existing classification schemes, across the nomenclature driven by disciplinary, linguistic and regional biases. To do this we conducted a review of reef geomorphic classifications, looking for consistencies and usage of terms that transcended divides in discipline^91^ (see Table 5).

#### 2.1.4. Previous reef classifications derived from satellite image data

Satellite technology has spawned a wealth of data on reefs, enabling large area coverage, with resolution of within reef variations. Initial approaches to reef mapping in the 1980s expanded our traditional viewpoint from single reef mapping and extent mapping to habitat mapping of whole reef systems^54^. Through the 1990s and early 2000s field survey techniques described above enabled more effective linkage of ecological surveys to remote sensing data^55,56^. Accessibility to higher spatial resolution images over larger areas in combination with detailed field data, physical attributes and object-based analysis resulted in large reef area mapping^34,39,40,57^. In the last five years, the increase in daily to weekly global coverage of this type of imagery, in combination with cloud-based processing capability has expanded to a global capability for reef mapping^14^. This is a new type of global information that requires a different approach to classification to make sense of complex natural systems at ocean scales.

The first challenge of creating the *Reef Cover* classification was to create a set of classes that related to natural science observations, despite using data pulled from remote sensing. Intra-reef zones defined by natural scientists often represent different biophysical /ecological communities that in turn reflect environmental gradients (e.g. in light, water flow) and geo-ecological processes (sediment deposition, reef vertical accretion) below the water that led to the arrangement^13^. However, these classes frequently also can be related to biophysical information on slope, depth and aspect that can be determined remotely. A thoughtfully prepared classification – that adheres to Stoddart’s (1978) classification principles, schemes - can support production of maps and other science (monitoring, management) that are still relevant to historic work but that can go forward with consistent definitions^34^.

### 2.2 Step 2. Development. Creator requirements - relating Reef Cover classes to remote sensing data

Development of mapping classes required a sensitive trade-off between the requirements of users (in terms of the level of detail needed, appropriate for scaling, consistent across regions, simple enough to be manageable but detailed enough to be understandable), the input resources available and the quality of the globally repeatable mapping methods to the producers.

which state that classes should be explicit, unique, comprehensible, and should follow the language of prior

While vast in terms of scalability, data producers are more constrained in terms of sensor capabilities such as spatial resolution (limited to pixels) and depth detection limits, and processing power (high numbers of map classes becoming more computationally expensive). Physical conditions and colour derived from remote sensing, along with their textural and spatial relationships, can be informative about reef zonation^57^, with depth and wave exposure being the important information to understanding geomorphology^58,59^.

To select a set of *Reef Cover* classes that could be defined by attributes available from most commonly available public access or commercial satellite data, but that also corresponded to common classes found in the classification literature, and also made logical sense from a user perspective, we looked for intersectionality between physical attribute data that can be derived from satellites but also help shape and define reef morphology. Below we relate data to ecological meaning.

#### 2.2.1. Physical attributes

The physical environment – light, waves and depth – plays a large role in controlling much of reef structural development and the ecological patterning across zones^36^. Underlying geomorphic structural features can almost always be characterised in terms of three core characteristics: i) depth, ii) slope angle and iii) exposure to waves (Fig. 2).

**Figure 2.**
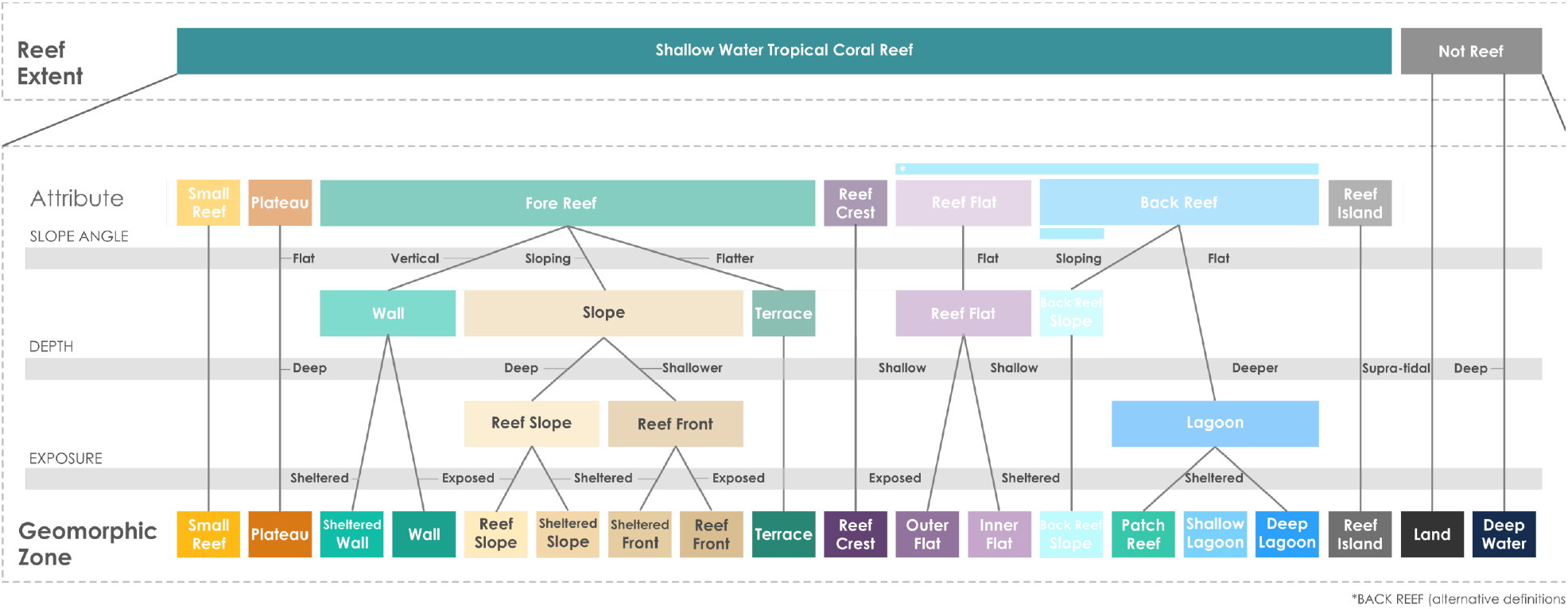
Physical attributes derived from remote sensing data such as depth, slope angle and exposure are enough to delineate some of the key reef zones in the classic literature. The coral reef classifier for global scale analyses of shallow water tropical coral reefs shows how relative measures can characterise reef zones.

##### Depth

Depth is a useful attribute for bridging human and machine classifications. Bathymetry can be derived from spectral information from satellites, since the absorption of light at specific wavelengths also has known relationships with water column depth^60^ but also relates to reef geomorphology^92^. Bathymetric data also provides the basis for other critical depth-derived products, slope and aspect, which are used to distinguish geomorphic classes and reef environmental parameters, e.g. exposure to breaking wave energy (Fig. 2).

From a natural science point of view, primary production drives much of the reef development, making light – and depth, due to rapid attenuation of light with depth - a critical driver of zonation and key attribute for classification of reef zones by natural scientists. For example, *Reef Crest* is often described as the shallowest part of the reef^46^, while *Lagoon* represents a deep depression in the reef structure^61,62^. Depth thresholds are sometimes included in definitions: a threshold of 10 m was suggested to differentiate true lagoons from shallow water areas^63^, and in another classification an 18 m threshold has been used to distinguish *Reef Front* from *Reef Slope*^13^. In the *Reef Cover* classification, depth was particularly important for distinguishing Fore Reef classes (e.g. Reef Slope, Terrace) from *Reef Crest* and *Reef Flat* classes (Table 1. Fig 2). Generally, tides and variability in water clarity and regional eustatic discrepancies in reef top depth (e.g., *Reef Flat* in Atlantic systems generally lie much lower with respect to tides than in the Indo-Pacific^62^) mean relative depths are more appropriate, which is why absolute numbers weren’t used in *Reef Cove*r definitions.

##### Slope

Slope angle, either absolute angle or discontinuities in angle acting as a break between zones, is an important differentiator of reef zones. *Reef Flat*s are defined as being horizontal ‘flattened”^64^ “flat-topped”^65^; *Fore Reef* slope zones often include references to slope angle (e.g. in one classification *Fore Reef* has been defined as “any area of the reef with an incline of between 0 and 45 degrees”^46^), and *Walls*– common on atolls - are often defined as “near vertical” features. Variability in slope continuity can also be an important way to demarcate zones. *Reef Crest* is sometimes defined as a demarcation point separating the *Fore Reef* from the *Reef Flat*^35,46,66^, while Montaggioni illustrated a range of representative profiles across atolls and barrier reefs, with convoluted profiles often allowing subdivisions of reef slope, particularly on fringing reefs which are less likely to show a uniform reef slope than an atoll^35^. Where water depth can be derived from remotely sensed spectral data, bathymetry can be used to directly calculate slope (i.e. by calculating the slope angle between a pixel and its neighbours) or by considering the local variance in depth (e.g. the standard deviation in depth values within some radius of each pixel).

In the *Reef Cover* classification, slope angle was important in distinguishing *Fore Reef* classes such as *Reef Slope* and *Reef Front*, from horizontal classes such as *Outer and Inner Reef Flat* and *Lagoons* (Table 1, Fig 2).

##### Exposure

Physical exposure of reefs is a key driver of zonation. *Reef Cre*sts – linked to wave breaking - are often described as “an area of maximum wave shoaling”, i.e. a zone that absorbs the greatest wave energy^46,64^. *Fore Reefs* are frequently sub-divided based on relative exposure (e.g. exposed vs sheltered slope, or windward vs leeward ^61^). Exposure influences profile shape and importantly the communities growing in the zone, so that slopes with identical profiles could have very different communities^61,67^. Sometimes these zones are related to the communities found there. Meanwhile, exposure across the reef means back-reef zones contain sheltered water bodies. Together with data on water depth and bathymetry, wave energy data was key for distinguishing key *Reef Cover* classes^68,69^.

##### Colour and texture

Sub-surface spectral reflectance data from satellites can provide measurements of reef colour and texture over large areas. Concentrations of photosynthetic pigments in coral, algae and seagrass as well as light scattering by inorganic materials means spectral reflectance measurements can also be used to measure biophysical properties of the reef^60^. Colour and texture information derived from satellites can be used to manually draw polygons around similar geomorphologic units or habitats, but provide the basis to drive image-based thematic mapping (such as digital number, radiance, reflectance) and texture, through spectral processing^58^. Texture measures are also used to improve classification by allowing spectrally similar substrates like corals and macroalgae to be distinguished. *Reef Flats*, for example, having a single driver of zonation, in contrast to several drivers on most other zones, makes benthic zonation particularly distinct^36^, and easily detectable as coloured bands in aerial images of reef flats. This allows colour and texture to be used to distinguish *Outer Reef Flats*, which have a greater component of photosynthetically active corals and algae, from *Inner Reef Flats* which appear brighter due to a higher proportion of sand build up in this depositional area (Table 2).

#### 2.2.2. Spatial Relationships

##### Size and shape

The size and shape of reef features can help determine *Reef Cover* class. Most large scale reef structural features appear elongate as the shelf constrains shape – and reef shapes can even help predict as they constrain accommodation space and influence deposition^70^. *Reef Flats*, for example, by definition boast the broadest horizontal extent of any geomorphic zone, typically 500 to 1000 m across, but reaching several kilometres in width across in large Pacific atoll structures^71^. *Lagoons* also tend to be broad in width although width and shape can be variable depending on reef type. Understanding some of these characteristics can help determine classes, although these are usually defined relationally rather than by application of size thresholds.

##### Neighbourhood and enclosure

Natural scientists agree that reefs feature three major geomorphic elements: a *Fore Ree*f, a *Reef Crest* and a *Back Reef* (although subdivisions and complexities exist around these). Because of the influence of large scale processes on reef development, these zones occur in order^13,35,36^. *Reef Crest* is arguably the most defining characteristic of any reef – the break point at which a sharply defined edge divides the shallower platform from a more steeply shelving reef front^71^, around which other geomorphic zones arranged in parallel^72^. As a result, spatial arrangement of zones can be informative for mapping (Table 3). For example, *Back Reef* is often defined as being contiguous to the *Reef Crest* (*Back Reef* is often defined as any reef feature found landward of the crest).

Enclosure to semi-enclosure within a bordering reef construction (Fairbridge defines lagoons as “bounded by reef”^63^) is another feature used classically to define reef zones, but that could also be derived from satellite imagery.

The *Reef Cover* typology presented is derived from earth observation data, but attempts to link classes to genetic process, social, ecological and geological importance. By focussing on the attributes of depth, light, exposure, colour and texture and spatial relationships that are common to both domains, our traditional biophysical knowledge of reefs can be integrated with remote-sensing capabilities. Attributes can be combined to make decision trees (provided in *Reef Cover* document^47^) to help use satellite data to map reefs at the global scale. The *Reef Cover* list of classes can all be distinguished from these physical attributes alone, supporting production of maps that are still relevant to existing work but that can allow computationally inexpensive determination of mapping classes to beyond what was previously possible^14^.

### 2.3. Step 3. User-requirements. Providing Reef Cover class descriptors that facilitates uptake and use

Computers have revolutionised our ability to classify multidimensional data sources, which allows mapping and modelling at far larger scales for the same effort compared to a human taxonomist. However, without proper consideration of the needs of the end user, classified data may not be effectively applied to conservation challenges. The *Reef Cover* classification was developed with five user-needs in mind: relevance, simplicity, transparency, accessibility and flexibility.

#### 2.3.1. Relevance

The typology needed to be informative to users with science or conservation challenges: map classes are not useful unless relevant. Different habitats within reefs contribute differently to biological and physical processes. For example, *Reef Crests* play a disproportionate role in coastal protection, dissipating on average 85% of the incoming wave energy and 70% of the swell energy^73,74^; *Reef Slopes* supply an order of magnitude more material to maintain island stability^44,75^; shallow *Reef Front* areas often host more coral biodiversity^36^; *Reef Flats* support herbivorous fish biomass^93^ and accessibility of *Lagoons* often affords them great cultural importance as places important for artisanal harvesting^76^. A classification that effectively captures the appropriate diversity of these habitats can therefore better inform social, biological and physical studies^28^, for example global conservation planning to safeguard reefs, for example, in order to meet the *Convention on Biodiversity* Aichi targets^26^. Map classes need to reflect differences of interest to a wide range of reef scientists, from oceanographers to paleoecologists and fisheries scientists – so careful consideration of natural history is important. Global mapping is usually to enable spatial comparisons, so a classification that is globally applicable was also important.

To explore relevancy, a cross-walk was performed between *Reef Cover* and a selection of major regional to global coral reef classification^42,64,77,78^, mapping^20,34,39^ and monitoring efforts^41^, to make sure important classes from established classifications had not been missed^47^ (Table 5).

#### 2.3.2. Simplicity

Simplicity was achieved by 1) choosing an appropriate mapping scale (internal geomorphic classes), 2) limiting the number of geomorphic classes (17 classes), and 3) providing clear (1 line) descriptors with additional information to address issues of semantic interoperability.

1. *Reef Cover* was developed to provide consistent mapping of reefs across very large areas: classification of geologic and ecological zones is much more amenable to mapping using remote sensing, given greater consistency in geomorphology across large biogeographic regions^30^. Satellite data has supported the development of several detailed regional “reef type” classifications, such as nine reef classes for the Great Barrier Reef from Landsat imagery^79^, six reef classes from the Torres Straight^80^ and 16 classes for the Red Sea from Quickbird^6^. However, local reef type classifications are not always applicable globally due to large regional discrepancies in Reef Type. As a result, detailed reef type typologies are more suitable for local to regional classifications^6,33^. For global mapping, an internal geomorphic approach is better. Finer spatial scale classifications from satellite data are also challenging, due to differences in the spatial scale at which spectral data can be generated (metres) and which benthic assemblages display heterogeneity (sub-metres)^30^. Medium spatial resolution multispectral data (5 to 30 m) is the most commonly used satellite information used for coral reef habitat mapping^57^, and classification of internal geomorphic structures may be best suited to this kind of data.
2. Reviews of habitat mapping from remote sensing found the number of map classes averages 18 at continental and global scales^11^. More than this can become confusing for users. Many coral reef classifications contain four or five hierarchical levels and high numbers of classes: the *Millennium Coral Reef Mapping Project (MCRMP)* was ambitious in developing a standardised typology that captured much of the reef type diversity, but despite defining over 800 reef classes defined at the finest (level 5, essential for local reef mapping) scale^34^, level 3 (68 classes) continue to be more popularly adopted in publications using this dataset. To keep the classification simple, *Reef Cover* was limited to 17 geomorphic classes, with simple one line definition provided. This limited number of classes was needed 1) to make it manageable for users, 2) to make it computationally manageable for very large (regional and global) data processing and 3) reduction in classes compared to MCRMP allowed for consistent automated mapping at the global scale – so that whole regions could be directly compared for monitoring and management.
3. Short definitions were provided in plain language for simplicity. Because there are many terminology in use, to address additional uses issues of semantic interoperability – each *Reef Cover* class definition also outlines other commonly used terms for concepts (synonymy) and explains different interpretations of the same meanings and understanding of the relations between concepts.

#### 2.3.3. Transparency

One barrier to the use of analysing and interpreting big data is user-friendliness. Of 79 coral reef mapping attempts reviewed (62 benthic coral reef maps, 6 geomorphic coral reef maps and 11 mixed), only 13% were accompanied by a clear classification that defined the meaning of map classes^12^. Describing how the classification relates to data (Step 2) and producing a detailed descriptor (Step 3) along with a diagram allows classification to be understood and also adopted for different projects. We also attempted to address transparency by relating *Reef Cover* classes to other major global mapping and monitoring efforts (Table 5) and providing a decision support tree for users^47^.

#### 2.3.4. Accessibility

Another barrier to the use of analysing and interpreting big data is access^81^. Much information remains locked behind paywalls, and additional barriers exist including discoverability. To promote accessibility and encourage use, all data were made publically available (see Data Records section for access). Terms were translated into different languages, as science published in just one language has been shown to hinder knowledge transfer and new findings getting through to practitioners in the field^94^.

#### 2.3.5. Flexibility

One criticism of thematic habitat maps derived from remote sensing is a lack of flexibility: categorical descriptions of habitats can be over simplified and discrete instead of dynamic and continuous meaning classifications limit the interpretation and questions that can be asked^82^. Flexibility issues were addressed by 1) not prescribing absolute thresholds to each class, instead providing information on how classes relate to each other (Tables 1-3) allowing a) map producers to adapt application of *Reef Cover* to their own needs, perhaps where different datasets are available and b) users to interpret with flexibility, 2) providing additional information (Standard Descriptors) including main features, exceptions to rule and broadness so to provide users with a broader understanding of hidden complexities when interpreting class meaning, 3) remaining open to feedback, we hope this *Reef Cover* version 1 can be improved upon with feedback from the community.

## 3. Data Records

*Reef Cover* classification (Version 1.0) presented in this paper has been made freely available as a list of map classes and descriptors “*Reef Cover Classification (v1). Internal coral reef class descriptors for global coral reef habitat mapping*” (pdf format) through Dryad^47^. The dataset includes 17 Reef Classes (Fig. 2) and five variables: *Standard Name* - short class name, *Standard Label* - longer class name, *Standard Description* – detailed class descriptor, including context and main attributes (highlighted), *Translations* - Standard Name in different languages, *Synonyms* - list of commonly used synonyms (Table 4). Diagrams of how each class relates to major reef types, and a Glossary of terms is also provided.

The dataset also contains an Attribute Table, explaining how classes relate to each other based upon on *Depth, Slope, Exposure, Substrate, Colour, Rugosity* and *Benthic Cover* data, information that might be available to mappers either from spectral reflectance, bathymetric, oceanographic or ecological datasets, and a Crosswalk Table comparing how *Reef Cover* classes align with other major reef mapping and monitoring efforts.

Two other resources accompany this data record in order to improve the uptake and use of *Reef Cover* classification. The first provides resources to support producers (remote-sensing data scientists) to create their own coral reef maps from *Reef Cover* (Methods). The second supplies users (coral reef practitioners and scientists) with a downloadable regional coral reef map of Micronesia developed using the *Reef Cover* classification (Maps).

### Methods^15^

Methods and codes for use in global coral reef habitat mapping are freely available <u>here</u>.

### Maps^90^

Static coral reef habitat maps of the Caroline and Mariana Islands produced using an amended version of the *Reef Cover* classification outlined are available to download <u>here</u>. Updates and further maps will be available in future as part of a dynamic datasource provided here^16^.

## 4. Technical Validation

The goal of the *Reef Cover classification* was to facilitate conversion of large amounts of remote-sensing data into regional-scale coral reef mapping products that can prove useful to supporting the work of coral reef practitioners and scientists. To test *Reef Cover*, the classification was applied to a large scale mapping exercise to convert 20TB of daily remote sensing data captured by Planet Dove satellites from across a three-million km^2^ area, into a sharable habitat map of some of the planets remotest coral reefs^90^. Challenges for producers in adopting the classification are described. Successes were assessed by getting feedback on maps to see if products could potentially be translated into conservation outcomes.

### 4.1 Application: using Reef Cover to produce coral reef maps

Two remote island chains in Micronesia were targeted for mapping. The Mariana Islands are a crescent-shaped archipelago of 15 volcanic islands making up the Commonwealth of the Northern Marianas (CNMI) and the Territory of Guam. The Caroline Islands, an archipelago of over 500 small islands, span a distance of 3500 km from Palau’s Hatohobei Reef in the westernmost point to Kosrae (Federated States of Micronesia) in the east. Spread across a 3 million km^2^ area, the wide dispersal of these isolated reef systems make them challenging to map using traditional in-water surveys, and regional mapping to date has relied on earth observation data. Large parts of Micronesia have been mapped by collation of existing maps (*ReefBase*’s Pacific Maps) and through detailed bathymetry surveys of US jurisdictions of the Mariana Islands by NOAA’s *Pacific Islands Benthic Habitat Mapping Center* (PIBHMC). The *Millennium Coral Reef Mapping Project* (MCRMP) used Landsat data to map 6000 km^2^ of reef across Palau, Federated States of Micronesia, the Marshall Islands and Gilbert Islands^83^ and the Japanese Ministry of the Environment mapped the region using AVNIR2 and Landsat data in 2010^84^. Many of these maps were used to produce the United Nations Environment Program (UNEP) World Conservation Monitoring Centre (UNEP-WCMC) *Global Distribution of Coral Reefs* data product, that is used as a global coral extent layer today^85^.

The *Reef Cover* classification was used to generate coral reef habitat maps of the Marianas and Caroline Islands from satellite data, following the methodology developed by Lyons, et al. ^14^ (Fig. 3). Sentinel-2, Landsat 8 and Planet Dove satellite-derived bathymetry and slope angle were used to identify major structural classes such as lagoons or reef slopes. In combination information on the substrate, calculated from colour, brightness levels and texture produced by spectral reflectance data from the satellites, geomorphic classes can be mapped (Fig. 3). A second map was created using benthic classes that could be reliability determined from the available satellite data (Supplementary Figure 1, see Reef Cover classification for class description). All code for developing the maps are publically available^15^.

**Figure 3.**
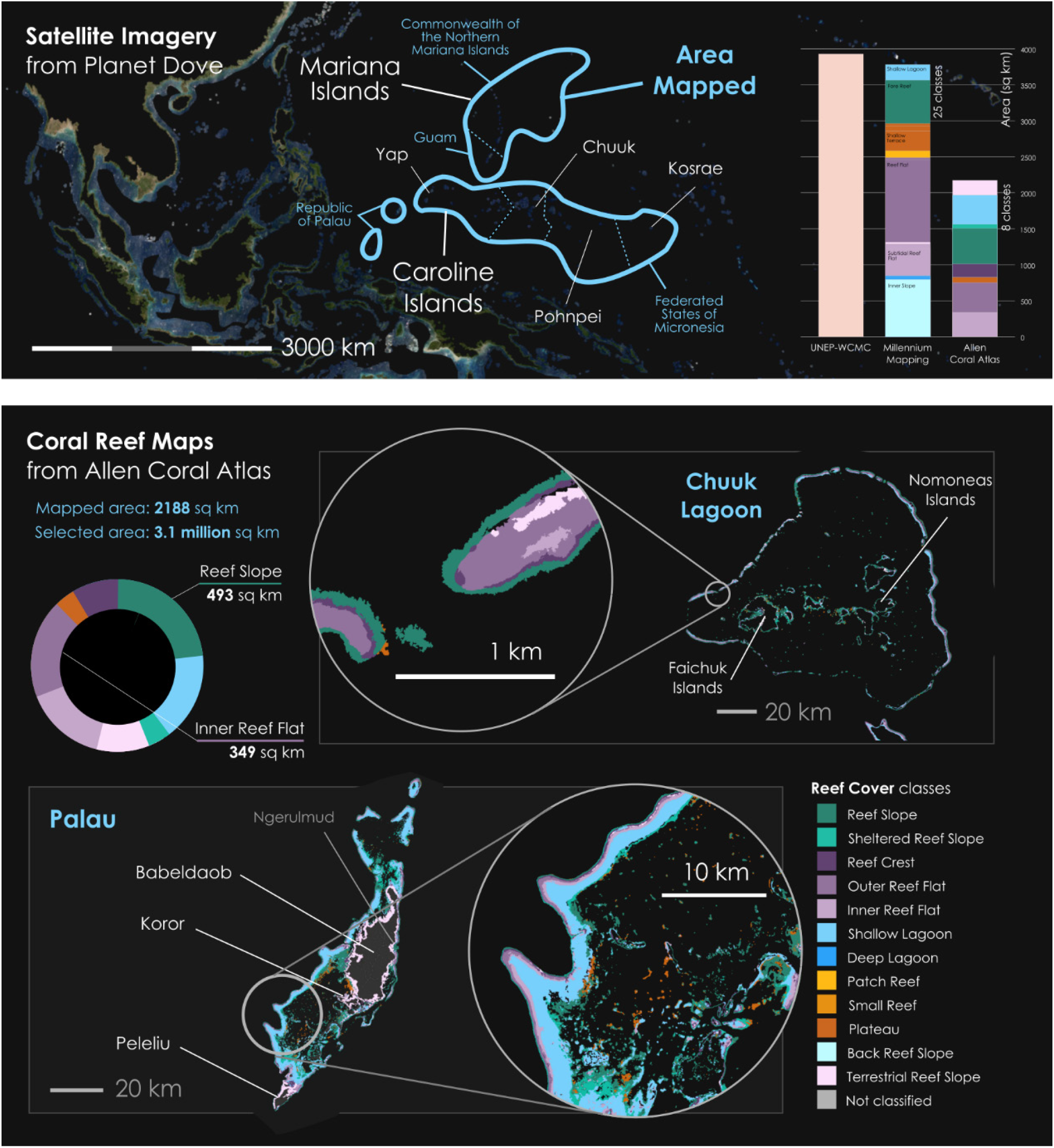
*Reef Cover* geomorphic classification scheme was applied to a coral reef mapping exercise: taking 3,072,192 km^2^ of spectral reflectance data (top panel) to generate coral reef habitat maps for reefs across Micronesia (e.g., Palau and Chuuk Lagoon, bottom panel). Over 2000 km^2^ of shallow coral reef of the Caroline Island chain (Republic of Palau, Federated State of Micronesia) and Mariana Island archipelago (Commonwealth of the Northern Marianas and US Territory of Guam) was mapped. *Reef Cover* classes (8 classes used) corresponded well with the 25 classes mapped by *Millennium Coral Reef Mapping Project* (barchart).

### 4.2 Assessment: challenges and benefits of Reef Cover

*Reef Cover* was adapted to suit the specifics of the mapping exercise (Supplementary Figure 1). Not all *Reef Cover* classes were used^47^: bathymetry was only available to a depth of 10 m, so the deeper *Reef Slope* and *Sheltered Reef Slope* classes were too deep to be applicable. *Reef Island* was not included due to the way the landmask was applied. Turbid water in inshore areas led to the requirement for a *Terrestrial Reef Flat* class (similar to fringing reef flat, but directly attached to land and subject to freshwater runoff, nutrients and sediment), that should be considered for next versions of *Reef Cover*.

A total of 2,188 km^2^ of reef were mapped, with eight classes used (Fig. 3). Dominating in terms of areal extent were *Reef Fron*t (called Reef Slope) comprising 23% of the shallow mapped reef area, the *Inner* and *Outer Reef Flat* classes (together 35%) and *Shallow Lagoon* (18%), reflecting that the region is typified by atolls or high islands with or without expansive lagoons. The spread of classes was largely in agreement with MCRMP outputs, who mapped more than twice as many classes (24 classes at the L4 level which aligns best with *Reef Cover*), but found five of these (Bay Exposed Fringing, Inner Slope, Fore Reef, Shallow Terrace and Subtidal Reef Flat) made up 85% of the map extent suggesting at very large scales some of the additional detail is redundant. A crosswalk (see *Reef Cover* Classification document) found the distribution of classes was comparable with the *Allen Coral Atlas* classes, with *Reef Front* (MCRMP *Forereef*) accounting for 16%), *Inner and Outer Reef Flat* seven classes combined) 43% and *Shallow Lagoon* (three classes combined) 30%. The total reef area mapped by MCRMP across the same area was 15,808 km^2^ and 38 classes at the L4 level (see crosswalk table), but once the dataset was filtered to remove deep classes (*Deep Reef, Deep Non-Reef*) and *Land*, this became 3,809 km2 and 24 classes, the higher extent value reflecting MCRMP’s ability to map deeper (e.g. 37 km^2^ of *Deep Lagoon* was mapped, while no *Deep Lagoon* was detected by the Allen Coral Atlas) and the inclusion of “Variable Depth” classes in this number.

Shallow reef extent estimates for the whole region were closer in value: *Allen Coral Atlas* estimated 1,569 km^2^ of *Reef Top*, MCRMP 1,293 km^2^ of *Shallow Reef* and UNEP-WCMC, who provide an outline of shallow reef, estimated an areal coverage of 1,590 km^2^ of shallow reef area.

Application of the *Reef Cover* classification allowed us to test the usability of the typology to map producers, and it was generally found to be flexible and adaptable. It could be aligned with existing schemes, but with just 17 options remained simple enough to allow for automated processing at very large scales with minimal human supervision.

### 4.3 Outcomes: use of maps to address real-world challenges

The main purpose of developing the *Reef Cover* classification was to improve the conversion of large amounts of remote sensing data into a mapped format that not only could visually communicate reef information, but would practically support coral reef science and conservation work. The mapped area is home to 353,000 islanders whose culture and many of whose livelihoods are related to coral reefs though fishing, recreation and tourism, yet reefs continue to be impacted by pressures from climate change (warming and high intensity storms) and local pollution (associated with crown-of-thorns sea star outbreaks) and overfishing^86^. The six nations remain committed to the *Micronesia Challenge*, an international conservation strategy across the six nations of Micronesia to conserve 30% of marine resources by 2020 in line with UN Sustainable Development goals. Mapping can help support marine spatial planning exercises: habitat maps such as the MCRMP of the area have been previously used to calculate estimates of fish biomass^87^.

In the production of the Micronesia maps, 125 GB of remote sensing spatial data (extracted from over 20TB of raw Planet Imagery) was converted into eight *Reef Cover* classes and displayed in an interactive web platform at AllenCoralAtlas.org. These products are now being used to a) communicate coral reef information and b) support conservation planning by coral reef practitioners in the region, from NGOs and research organisations to the communities that live on the islands. For example, maps of reefs in the Republic of Palau have been integrated into a spatial analysis model by *The Nature Conservancy*, to help the organisation identify where to site aquaculture as part of a national-scale project in partnership with the *Palau Environmental Quality Protection Board, Bureau of Marine Resources* and *Palau Community College*. Meanwhile, maps of the reefs across the Federated States of Micronesia are being used by the *Waitt Institute* to help plan research expeditions to remote submerged reefs, as well as support marine spatial planning exercises in the area. Researchers commented that, “*for a place like the FSM that has a pretty crude current estimate for distribution of coral reef structures, the Atlas is a huge value for them and a very necessary tool in the marine spatial planning process*” and “*These are some of the most critical pieces of information to create policy*.” In Micronesia’s Outer Islands, *One People One Reef*, a collaboration of communities and scientists, believed the open-access and easy to use format of the products could prove useful in communicating ideas and connecting with remote communities involved in traditional management. Finally, researchers leading the *Micronesia Challenge* at the University of Guam found the maps to be a dynamic and interactive communication tool during stakeholder meetings, allowing meeting participants to design and review current marine protected areas, and negating the need for an on-site GIS expert during meetings.

There is still room to improve both the mapping process and user experience: the number of classes used in the typology was refined due to data limitations (Supplementary Figure 1). A survey of 26 users found 42% of those frequently worked in places with little or no internet connectivity, and 10% needed to work on a mobile device^88^.

## 5. Usage Notes

Read through the *Reef Cover* classification before applying the classification scheme to your own mapping process following the methodology^14^, or interpreting Micronesia maps^90^ or other Allen Coral Atlas mapping products^15^. You might want to consider setting your own thresholds for attributes such as slope angle, depth, depending on the scope of your datasets, and what data sources are available. *Reef Cover* contains suggested classes, and you may want to add or remove classes (see example from Technical Validation). For example, very few other mapping exercises consider “Sheltered Reef Slope” vs “Reef Slope”.

### 5.1 Using western Micronesia maps (and other Allen Coral Atlas resources)

Static geomorphic digital maps of shallow coral reef habitat across the Caroline and Mariana Islands in western Micronesia (Version 1.0) are downloadable in a number of formats as a tar file (western-micronesia-aca.tar.gz) from <u>here</u>. MacOS can open tar and tar.gz files by default with the Archive UtilityUsers, Windows users will need an external program (7-Zip or WinRar) to extract files. The compressed folder *Western-Micronesia* contains three sub-folders: *geomorphic, benthic* and *boundary*. The *geomorphic* folder contains digital geomorphic coral reef habitat maps described in this study, mapped to 12 Allen Coral Atlas *Reef Cover* classes (Fig 3, class descriptor available in *Reef Cover* document^47^) and stored in a variety of standard geospatial formats. Shapefiles are available for use with most GIS software (geomorphic.shp, geomorphic.shx, geomorphic.dbf and geomorphic.prj), a KML file for viewing in a browser such as Google Earth (geomorphic.kml), and GeoJSON (geomorphic.geojson). The *benthic* folder contains the same file formats, but same reefs have been mapped to six benthic classes: *Seagrass, Coral Habitat, Rubble, Sand, Microalgal Mats* and *Rock* (see *Reef Cover* for class descriptors^47^), and the *boundary* folder contains the outline of the area mapped, again in the same three formats. Users will be able to access updates and download further regional maps through via this <u>Zenodo link</u> which links through dynamic maps currently hosted to AllenCoralAtlas.org^16^, where up-to-date usage notes are available through the “FAQ” section. Further support is available by emailing. The Allen Coral Atlas currently has two mapped themes: one that displays *global geomorphic zones* (12 classes) and another for *global benthic zones* (6 classes) commonly associated with shallow water tropical coral reefs (Supplementary Figure 1). The adapted *Reef Cover* Allen Coral Atlas map geomorphic classes and benthic classes are available in the *Reef Cover* document.

Map users should bear in mind the following limitations.

- *Bathymetric constraint*. Satellite data limitations mean maps display reef features 10 m deep (for benthic) and 20 m (for geomorphic) or shallower. Features beyond 20 m - for example mesophotic reefs found from 30 – 150 m depth, drowned reefs and submerged reef platforms - will not be visible on the global scale maps. Depth on the map is relative – i.e. not linked to a vertical datum, which is why class definitions provide approximate depths. Very steep reef features – like reef walls – may not appear on the map, or their true extent (width) will be under-represented in two-dimensional space. This is why some *Reef Cover* classes (e.g. Wall and Sheltered Wall, Reef Slope and Sheltered Reef Slope) do not appear on Atlas maps.
- *Latitudinal constraint*. Only shallow water tropical reefs occurring 30° either side of the equator are displayed. Limestone reefs tend not to develop at higher latitudes, due to cooler sea temperatures. Other biogenic reef types – e.g. serpulid reefs, *Halimeda* reefs – are also not captured.
- *Biological constraint*. Despite its name, the *Allen Coral Atlas* is actually a Coral Reef (and not a Coral) Atlas: remote sensing can detect reefs well but is less good at detecting individual corals. Moreover, photosynthetic pigments mean living corals and algae (including turfs) are difficult to distinguish by earth-observation systems. Therefore, corals growing in or on non-reef habitats are not represented here. For an interactive map of coral species distribution, visit <u>Coral Geographic</u>.
- *Temporal constraint*. Maps represent the current detectable distribution of coral reefs, as seen from space. Large (kilometre) scale coral reef limestone structures (displayed in the global geomorphic map) develop over millennia, but across the surface of these structures benthic features – such as the distributions of corals and algae – will fluctuate on seasonal to decadal timeframes, meaning benthic map classes may become outdated. Maps are being continually updated as more data become available: please check you are using the latest map version.
- *Turbidity constraint*. Coral reefs located in turbid areas are not well mapped by the *Allen Coral Atlas*. An additional map class “Terrestrial Reef Flat” describes “a broad, flat, shallow semi-exposed area of fringing reef found directly attached to land at one side, and subject to freshwater runoff, nutrients and sediment” to account for difficulty of mapping turbid near-shore areas^89^.
- *Spatial constraint*. The *Allen Coral Atlas* is designed as a large extent mapping exercise, hence local scale applications could potentially be limited. Maps are designed to capture coral reef distributions and features at the regional to ocean-basin scales. It is less appropriate for local exploration, e.g. below the “at the reef” level. Small reef features, such as individual coral colonies, small bommies, boulders and channels may be beyond the map’s detection limits.

Where “no class” may be assigned. While the Atlas takes global level mapping to the next level in terms of detail, when exploring the map, keep in mind that the main value in a global map is being able to compare reef features across scales of kilometres to ocean basins. For a more detailed, metre-scale understanding of an individual reef it will always be better to source a local map (e.g. Level 5 local maps).

### 5.2 Using *Reef Cover* to create your own maps

The *Allen Coral Atlas* classification process and links to the code used to produce the maps, along with the data themselves can be accessed through *Zenodo* link <u>here</u>^14,15^. This link provides resource and citation for code, standards and publications arising from the Allen Coral Atlas, including Google Earth Engine source code for mapping algorithms: https://github.com/CoralMapping/gee-mapping-source. This repository contains all the Google Earth Engine source code that generates the mapping outputs on the *Allen Coral Atlas*, including maps and validation statistics. Adapted *Reef Cover* class definitions (Supplementary Figure 1) can also be found in Reef Cover document^47^.

## Supporting information

Reef Cover v1

## 6. Code Availability

The Google Earth Engine code used to produce the Micronesia maps presented in this dataset (and all regions) is fully open access (https://github.com/CoralMapping/gee-mapping-source) and in full detail at https://zenodo.org/record/3833246#.XtBAN_gzaUl^15^.

## Acknowledgements

This work was initiated and funded primarily through Paul Allen Philanthropies and Vulcan Inc. as part of the Allen Coral Atlas. We acknowledge the late Paul Allen and Ruth Gates for their fundamental vision and drive to enable us to work together on this critical reef mapping problem. Project partners providing financial, service and personnel include: Planet Inc., National Geographic, University of Queensland, Arizona State University, and University of Hawai’i. Significant support has also been provided by Google Inc., Great Barrier Reef Foundation, and Trimble (Ecognition). Contributors to establishing and running the project include: Vulcan Inc. [James Deutsch, Lauren Kickam, Paulina Gerstner, Charlie Whiton, Kirk Larsen, Sarah Frias Torres, Kyle Rice, Janet Greenlee]; Planet Inc. [Andrew Zolli, Trevor McDonald, Joe Mascaro, Joe Kington]; University of Queensland [Chris Roelfsema, Stuart Phinn, Emma Kennedy, Mitch Lyons, Nicholas Murray, Doddy Yuwono, Dan Harris, Eva Kovacs, Rodney Borrego, Meredith Roe, Jeremy Wolff, Katherine Markey, Alexandra Ordonez, Chantal Say, Paul Tudman]; Arizona State University [Greg Asner, Dave Knapp, Jiwei Li, Yaping Xu, Nick Fabina, Heather D’Angelo]; and National Geographic [Helen Fox, Brianna Bambic, Brian Free, Zoe Lieb] and Great Barrier Reef Foundation [Petra Lundgren, Kirsty Bevan].

This work was also supported by the Great Barrier Reef Marine Park Authority, and we are grateful to Mike Ronan and Maria Zann from the Queensland Government for guidance with classification.

Nick Murray is the recipient of an Australian Research Council Australian Discovery Early Career Award (DE190100101) funded by the Australian Government.

We would also like to thank Coral Reef Classification workshop participants Emma Kennedy, Chris Roelfsema, Eva Kovacs, Daniel Harris, Mitchell Lyons, Greg Webb, Doddy Yowono and Atefeh Sansoleimani (University of Queensland), Sarah Hamylton (University of Wollongong), Javier Leon (Southern Cross University) and Stephanie Duce (James Cook University).

We are grateful to Serge Andréfouet, Hiroya Yamano, Sam Purkis, Mark Spalding and Sarah Hamylton for providing expert guidance and helpful feedback on an early manuscript draft.

## References

[1] Runting, R. K., Phinn, S., Xie, Z., Venter, O. & Watson, J. E. M. Opportunities for big data in conservation and sustainability. Nature Communications 11, 2003, doi:10.1038/s41467-020-15870-0 (2020).

[2] Hughes, T. P. et al. Spatial and temporal patterns of mass bleaching of corals in the Anthropocene. Science 359, 80–83, doi:10.1126/science.aan8048 (2018).

[3] Hoegh-Guldberg, O., Jacob, D. & Taylor, M. in Special Report on Global Warming of 1.5°C (Intergovermental Panel on Climate Change, 2018).

[4] Madin, E. M. P., Darling, E. S. & Hardt, M. J. Emerging technologies and coral reef conservation: Opportunities, challenges, and moving forward. Frontiers in Marine Science 6, doi:10.3389/fmars.2019.00727 (2019).

[5] Sokal, R. R. Classification: Purposes, Principles, Progress, Prospects. Science 185, 1115–1123, doi:10.1126/science.185.4157.1115 (1974).

[6] Rowlands, G., Purkis, S. & Bruckner, A. Diversity in the geomorphology of shallow-water carbonate depositional systems in the Saudi Arabian Red Sea. Geomorphology 222, 3–13, doi:https://doi.org/10.1016/j.geomorph.2014.03.014 (2014).

[7] Garshol, L. Metadata? Thesauri? Taxonomies? Topic maps! Making sense of it all. Journal of Information Science 30, 378–391 (2004).

[8] Kuchler, D. Geomorphological nomenclature: reef cover and zonation on the Great Barrier Reef. (1986).

[9] Roelfsema, C., Phinn, S., Jupiter, S., Comley, J. & Albert, S. Mapping coral reefs at reef to reef-system scales, 10s–1000s km2, using object-based image analysis. International Journal of Remote Sensing 34, 6367–6388, doi:10.1080/01431161.2013.800660 (2013).

[10] Green, E. P., Mumby, P. J., Edwards, A. J. & Clark, C. D. A review of remote sensing for the assessment and management of tropical coastal resources. Coastal Management 24, 1–40, doi:10.1080/08920759609362279 (1996).

[11] Morales-Barquero, L., Lyons, M. B., Phinn, S. R. & Roelfsema, C. M. Trends in remote sensing accuracy assessment approaches in the context of natural resources. Remote Sensing 11, 2305 (2019).

[12] Roelfsema, C. M. & Phinn, S. R. in Coral Reef Remote Sensing (ed J.A. Goodman) 375–401 (Springer, 2013).

[13] Stoddart, D. R. Ecology and morphology of recent coral reefs. Biological Reviews 44, 433–498, doi:10.1111/j.1469-185X.1969.tb00609.x (1969).

[14] Lyons, M. B. et al. Mapping the world’s coral reefs using a global multiscale earth observation framework. Remote Sensing in Ecology and Conservation n/a, doi:10.1002/rse2.157 (2020).

[15] Allen Coral Atlas Partnership. CoralMapping/AllenCoralAtlas: DOI release (Version 1.0). [Data set] Zenodo. http://doi.org/10.5281/zenodo.3833246 (2020)

[16] Allen Coral Atlas. Allen Coral Atlas Partnership and Vulcan, Inc. and licensed CC BY 4.0 (https://creativecommons.org/licenses/by/4.0/). (2019)

[17] Pendleton, L. et al. Coral reefs and people in a high-CO2 world: where can science make a difference to people? PLOS ONE 11, e0164699, doi:10.1371/journal.pone.0164699 (2016).

[18] Spalding, M. et al. Mapping the global value and distribution of coral reef tourism. Marine Policy 82, 104–113, doi:https://doi.org/10.1016/j.marpol.2017.05.014 (2017).

[19] Moberg, F. & Folke, C. Ecological goods and services of coral reef ecosystems. Ecological Economics 29, 215–233, doi:https://doi.org/10.1016/S0921-8009(99)00009-9 (1999).

[20] Spalding, M., Ravilious, C. & Green, E. World Atlas of Coral Reefs. (University of California Press, 2001).

[21] Wolff, N. H. et al. Global inequities between polluters and the polluted: climate change impacts on coral reefs. Global Change Biology 21, 3982–3994, doi:10.1111/gcb.13015 (2015).

[22] Heron, S. F., Maynard, J. A., van Hooidonk, R. & Eakin, C. M. Warming trends and bleaching stress of the world’s coral reefs 1985–2012. Scientific Reports 6, 38402, doi:10.1038/srep38402

[23] Burke, L., Reytar, K., Spalding, M. & Perry, A. Reefs at Risk Revisited. (ISBN 978-1-56973-762-0, 2011).

[24] van Hooidonk, R. et al. Local-scale projections of coral reef futures and implications of the Paris Agreement. Scientific Reports 6, 39666, doi:10.1038/srep39666

[25] Beyer, H. L. et al. Risk-sensitive planning for conserving coral reefs under rapid climate change. Conservation Letters 0, e12587, doi:10.1111/conl.12587 (2018).

[26] Gairin, E. & Andréfouët, S. Role of habitat definition on Aichi Target 11: Examples from New Caledonian coral reefs. Marine Policy 116, 103951, doi:https://doi.org/10.1016/j.marpol.2020.103951 (2020).

[27] Foo, S. A. & Asner, G. P. Scaling up coral reef restoration using remote sensing technology. Frontiers in Marine Science 6, doi:10.3389/fmars.2019.00079 (2019).

[28] Roelfsema, C. M. et al. Habitat maps to enhance monitoring and management of the Great Barrier Reef. Coral Reefs, doi:10.1007/s00338-020-01929-3 (2020).

[29] McManus, J. & Ablan, M. ReefBase: a global database on coral reefs and their resources. (1996).

[30] Purkis, S. J. Remote sensing tropical coral reefs: The view from above. Annual Review of Marine Science 10, 149–168, doi:10.1146/annurev-marine-121916-063249 (2018).

[31] Andréfouët, S. et al. Multi-site evaluation of IKONOS data for classification of tropical coral reef environments. Remote Sensing of Environment 88, 128–143, doi:https://doi.org/10.1016/j.rse.2003.04.005 (2003).

[32] Hedley, J. D. et al. Remote sensing of coral reefs for monitoring and management: A review. Remote Sensing 8, 118–157 (2016).

[33] Hopley, D. in Perspectives on Coral Reefs (ed D.J. Barnes) (Australian Institute of Marine Science, 1983).

[34] Andréfouët, S. et al. in Proceedings of 10th International Coral Reef Symposium.

[35] Montaggioni, L. F. & Braithwaite, C. J. Quaternary Coral Reef Systems: history, development processes and controlling factors. (2009).

[36] Done, T. J. in Perspectives on Coral Reefs (ed D.J. Barnes) 107–147 (Australian Institute of Marine Science, 1983).

[37] Andréfouët, S. in Encyclopedia of Modern Coral Reefs: Structure, Form and Process (ed David Hopley) 906–910 (Springer Netherlands, 2011).

[38] Hamylton, S., Andréfouët, S. & Spencer, T. Comparing the information content of coral reef geomorphological and biological habitat maps, Amirantes Archipelago (Seychelles), Western Indian Ocean. Estuarine, Coastal and Shelf Science 111, 151–156, doi:https://doi.org/10.1016/j.ecss.2012.06.001 (2012).

[39] Purkis, S. J. et al. High-resolution habitat and bathymetry maps for 65,000 sq. km of Earth’s remotest coral reefs. Coral Reefs (2019).

[40] Monaco, M. E., Battista, T., Kendall, M. S., Wedding, L. S. & Clarke, A. M. National Summary of NOAA’s Shallow-water Benthic Habitat Mapping of U.S. Coral Reef Ecosystems. (Prepared by the NCCOS Center for Coastal Monitoring and Assessment Biogeography Branch, Silver Spring, MD., 2012).

[41] Lang, J. & Marks, K. AGRRA (Atlantic and Gulf Rapid Reef Assessment): Reef Terms, 2018).

[42] Kuchler, D. Reef cover and zonation classification system for use with remotely sensed Great Barrier Reef data. (Townsville, 1986).

[43] Holthus, P. F. in Small Islands: marine science and sustainable development (ed George A Maul) (American Geophysical Union, 1996).

[44] Hamylton, S. M. & Mallela, J. Reef development on a remote coral atoll before and after coral bleaching: A geospatial assessment. Marine Geology 418, 106041, doi:https://doi.org/10.1016/j.margeo.2019.106041 (2019).

[45] Hamylton, S. M. Mapping coral reef environments:A review of historical methods, recent advances and future opportunities. Progress in Physical Geography: Earth and Environment 41, 803–833, doi:10.1177/0309133317744998 (2017).

[46] Mumby, P. J. & Harborne, A. R. Development of a systematic classification scheme of marine habitats to facilitate regional management and mapping of Caribbean coral reefs. Biological Conservation 88, 155–163, doi:https://doi.org/10.1016/S0006-3207(98)00108-6 (1999).

[47] Reef Cover Classification (v1). Reef Cover (Version 1.0): Internal coral reef class descriptors for global coral reef habitat mapping., [Dataset] Kennedy, E. & Roelfsema CR Editors. in prep: Dryad doi:10.5061/dryad.7h44j0zrr (2020)

[48] González-Rivero, M. et al. Scaling up ecological measurements of coral reefs using semi-automated field image collection and analysis. Remote Sensing 8, 30 (2016).

[49] Wells, J. W. in Treatise on marine ecology and paleoecology Vol. 1 (Geological Society of America, 1957).

[50] Darwin, C. The Structure and Distribution of Coral Reefs. Being the first part of the geology of the voyage of the Beagle, under the command of Capt. Fitzroy, R.N. during the years 1832 to 1836. 214 pp (Smith Elder & Co, 1843).

[51] Martin, L. SCUBA diving explained: Questions and answers on physiology and medical aspects of SCUBA diving. (1997).

[52] Fairbridge, R. W. Recent and Pleistocene coral reefs of Australia. The Journal of Geology 58, 330–401, doi:10.1086/625751 (1950).

[53] MacIntyre, I. G. A diver-operated hydraulic drill for coring submerged substrates. Atoll Research Bulletin 185, 21–25 (1975).

[54] Jupp, D. L. B. et al. Remote sensing for planning and managing the Great Barrier Reef of Australia. Photogrammetria 40, 21–42, doi:https://doi.org/10.1016/0031-8663(85)90043-2 (1985).

[55] Joyce, K. E., Phinn, S. R., Roelfsema, C. M., Neil, D. T. & Dennison, W. C. Combining Landsat ETM+ and Reef Check classifications for mapping coral reefs: a critical assessment from the southern Great Barrier Reef, Australia. Coral Reefs 23, 21–25, doi:10.1007/s00338-003-0357-7 (2004).

[56] Goodman, J. A., Purkis, S. & Phinn, S. Coral Reef Remote Sensing: a Guide for Mapping, Monitoring and Management. 436 (Springer Netherlands, 2013).

[57] Roelfsema, C., Phinn, S., Jupiter, S., Comley, J. & Albert, S. Mapping coral reefs at reef to reef-system scales, 10s–1000s km2, using object-based image analysis AU - Roelfsema, Chris. International Journal of Remote Sensing 34, 6367–6388, doi:10.1080/01431161.2013.800660 (2013).

[58] Yamano, H. in Coral Reef Remote Sensing: A Guide for Mapping, Monitoring and Management (eds James A. Goodman, Samuel J. Purkis, & Stuart R. Phinn) 51–78 (Springer Netherlands, 2013).

[59] Roelfsema, C. et al. Coral reef habitat mapping: A combination of object-based image analysis and ecological modelling. Remote Sensing of Environment 208, 27–41, doi:https://doi.org/10.1016/j.rse.2018.02.005 (2018).

[60] Phinn, S. R., Hochberg, E. M. & Roelfsema, C. M. in Coral Reef Remote Sensing: A Guide for Mapping, Monitoring and Management (eds James A. Goodman, Samuel J. Purkis, & Stuart R. Phinn) 3–28 (Springer Netherlands, 2013).

[61] Kuchler, D. Reef cover and zonation classification system for use with remotely sensed Great Barrier Reef data: uder guide and handbook. (1987).

[62] Woodroffe, C. D. & Biribo, N. in Encylopedia of Modern Coral Reefs (Springer, The Netherlands, 2011).

[63] Fairbridge, R. W. in Geomorphology. Encyclopedia of Earth Science.594–598 (Springer Berlin Heidelberg, 1968).

[64] Zitello, A. G. et al. Shallow-Water Benthic Habitats of St. John, U.S. Virgin Islands. (Silver Spring, MD. 53 pp., 2009).

[65] Collins, L. B. in Encyclopedia of Modern Coral Reefs: Structure, Form and Process (ed David Hopley) 896–902 (Springer Netherlands, 2011).

[66] Kennedy, D. M. & Woodroffe, C. D. Fringing reef growth and morphology: a review. Earth-Science Reviews 57, 255–277, doi:https://doi.org/10.1016/S0012-8252(01)00077-0 (2002).

[67] Geister, J. in Proceedings of Third International Coral Reef Symposium.(Rosenstiel School of Marine and Atmospheric Science).

[68] Callaghan, D. P., Leon, J. X. & Saunders, M. I. Wave modelling as a proxy for seagrass ecological modelling: Comparing fetch and process-based predictions for a bay and reef lagoon. Estuarine, Coastal and Shelf Science 153, 108–120, doi:https://doi.org/10.1016/j.ecss.2014.12.016 (2015).

[69] Purkis, S. J., Rowlands, G. P. & Kerr, J. M. Unravelling the influence of water depth and wave energy on the facies diversity of shelf carbonates. Sedimentology 62, 541–565, doi:10.1111/sed.12110 (2015).

[70] Purkis, S. J., Kohler, K. E., Riegl, B. & Rohman, S. O. The statistics of natural shapes in modern coral reef landscapes. The Journal of Geology 115, 493–508 (2007).

[71] Blanchon, P. in Encyclopedia of Modern Coral Reefs: Structure, Form and Process (ed David Hopley) 469–486 (Springer Netherlands, 2011).

[72] Montaggioni, L. F. History of Indo-Pacific coral reef systems since the last glaciation: Development patterns and controlling factors. Earth-Science Reviews 71, 1–75, doi:https://doi.org/10.1016/j.earscirev.2005.01.002 (2005).

[73] Ferrario, F. et al. The effectiveness of coral reefs for coastal hazard risk reduction and adaptation. Nature Communications 5, 3794, doi:10.1038/ncomms4794 (2014).

[74] Lowe, R. J. et al. Spectral wave dissipation over a barrier reef. Journal of Geophysical Research: Oceans 110, doi:10.1029/2004jc002711 (2005).

[75] Hamylton, S. Will coral islands maintain their growth over the next century? A deterministic model of sediment availability at lady elliot island, great barrier reef. PLOS ONE 9, e94067, doi:10.1371/journal.pone.0094067 (2014).

[76] Aswani, S. & Vaccaro, I. Lagoon Ecology and Social Strategies: Habitat Diversity and Ethnobiology. Human Ecology 36, 325–341, doi:10.1007/s10745-007-9159-9 (2008).

[77] Holthus, P. F. & Maragos, J. E. in Marine and Coastal Biodiversity in the Tropical Island Pacific Region 239–278 (1995).

[78] Nichol, S. L. et al. Geomorphological classification of reefs. 27 (Geoscience Australia, 2016).

[79] Hopley, D., Parnell, K. E. & Isdale, P. J. The Great Barrier Reef Marine Park: Dimensions and Regional Patterns. Australian Geographical Studies 27, 47–66, doi:10.1111/j.1467-8470.1989.tb00591.x (1989).

[80] Leon, J. X. & Woodroffe, C. D. Morphological characterisation of reef types in Torres Strait and an assessment of their carbonate production. Marine Geology 338, 64–75, doi:https://doi.org/10.1016/j.margeo.2012.12.009 (2013).

[81] Pendleton, L. H. et al. Disrupting data sharing for a healthier ocean. ICES Journal of Marine Science 76, 1415–1423, doi:10.1093/icesjms/fsz068 (2019).

[82] Coops, N. C. & Wulder, M. A. Breaking the Habit(at). Trends in Ecology & Evolution 34, 585–587, doi:https://doi.org/10.1016/j.tree.2019.04.013 (2019).

[83] Imars-USF, I. I. d. R. p. l. D. (Cambridge (UK): UNEP World Conservation Monitoring Centre, 2005).

[84] Kakuta, S. et al. Satellite-based mapping of coral reefs in east asia, micronesia and melanesia regions International Archives of the Photogrammetry, Remote Sensing and Spatial Information Science 38, 534–537 (2010).

[85] UNEP-WCMC, Centre, W., WRI & TNC. (Cambridge (UK): UNEP World Conservation Monitoring Centre. URL: data.unep-wcmc.org/datasets/13, 2010).

[86] Houk, P. et al. The Micronesia Challenge: Assessing the relative contribution of stressors on coral reefs to facilitate science-to-management feedback. PLOS ONE 10, e0130823, doi:10.1371/journal.pone.0130823 (2015).

[87] Harborne, A. Modelling and mapping fishing pressure and the current and potential standing stock of coral reef fishes in five jurisdictions of Micronesia. (2017).

[88] Vulcan Inc. Insights research: Coral Atlas Feature Survey Results. 12 (2020).

[89] Kennedy, E. V. & Roelfsema, C. Global Geomorphic and Global Benthic Map Class Descriptors: A Guide for how to use the Allen Coral Atlas global geomorphic and benthic habitat maps.. 24 (Remote Sensing and Research Centre, School of Earth And Environmental Sciences, University of Queensland, Brisbane, Australia, 2020).

[90] Allen Coral Atlas Partnership. Allen Coral Atlas maps of the benthic cover and geomorphic zones of coral reefs in western Micronesia (Version 1.0) [Data set]. Zenodo. http://doi.org/10.5281/zenodo.3953053 (2020).

[91] Kennedy et al (in prep) Classification of coral reefs: a review of geomorphological terminology and definitions (in prep).

[92] Hochberg, E.J., S.A. Peltier, and S. Maritorena, Trends and variability in spectral diffuse attenuation of coral reef waters. Coral Reefs, (2020).

[93] Bellwood, D.R., et al., The role of the reef flat in coral reef trophodynamics: Past, present, and future. Ecology and Evolution, 2018. 8(8): p. 4108–4119.

[94] Amano, T., J.P. González-Varo, and W.J. Sutherland, Languages are still a major barrier to global science. PLOS Biology, 2016. 14(12): p. e2000933.

