## Supplementary material for "Reef Cover: a coral reef classification to guide global habitat mapping from remote sensing": Reef Cover v1

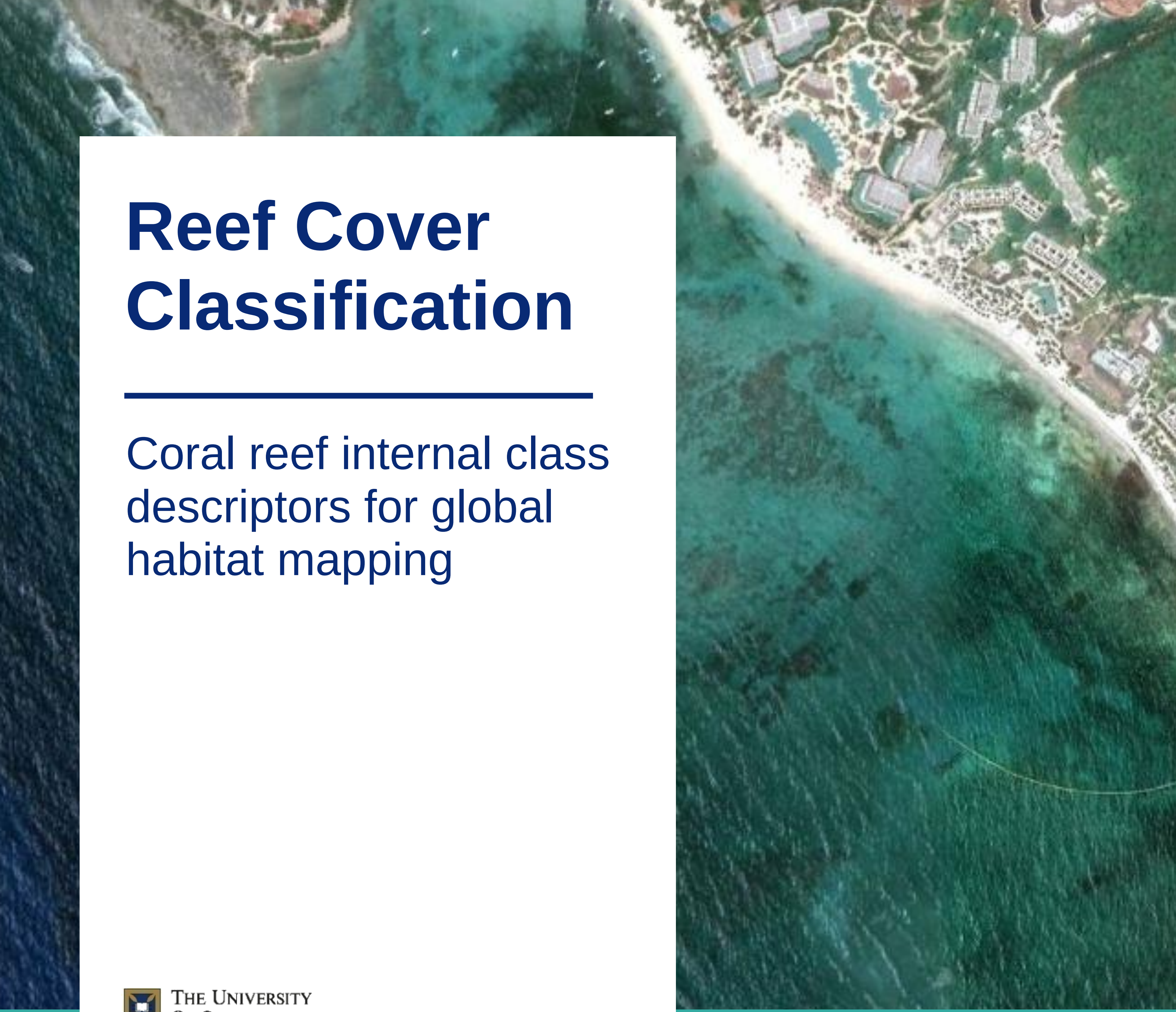

### Reef Cover Classification

#### Coral reef internal class descriptors for global habitat mapping

THE UNIVERSITY OF QUEENSLAND
AUSTRALIA

**VERSION 1, JULY 2020**

**AUTHORED BY: Kennedy EV & Roelfsema CR**

Kovacs E, Lyons M, Borrego-Acevedo R, Roe M, Yuwono D, Wolff J, Tudman P, Murray N, Phinn S

### Reef Cover Classification

#### Description of seventeen shallow coral reef geomorphic mapping categories for satellite-based global mapping

This document contains intra-reef geomorphic **Reef Cover** descriptors, developed for shallow-water tropical coral reef habitat mapping. **Reef Cover** describes spatially explicit features that can be used in different academic disciplines (e.g. geography, ecology, oceanography, marine sciences, marine policy and planning) as a geographical reference (e.g. to support political decision making or scientific research). The categorization is designed to allow broad-scale comparisons between reef ecosystems to support reef research at regional to global scales, and to both guide development and aid interpretation of global coral reef geomorphic maps.

#### Standard Name.

##### Standard Label.

Standard Description. This longer descriptor details diagnostic attributes of the class, with key **environmental attributes** highlighted in green and specific **technical attributes** or **thresholds** related to mapping in blue. Relationships to **other classes** (dark blue) are also clarified, with numbered citations to relevant literature [X].

Context. Putting the class into context, including linking to and other **terms** (purple) defined in the Glossary. The final sentence relates to the potential relevance of this class to users.

English • Bahasa • Española • Française • عربي

*Other common names • Alternative terms • Synonyms*

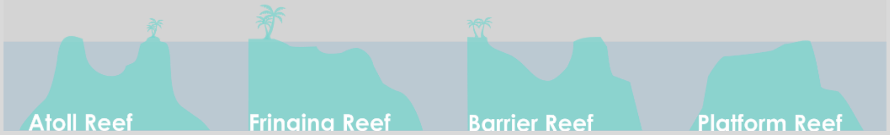


The seventeen Reef Cover categories presented are derived from a combination of *a priori* classifications that draw on foundational knowledge of coral reef structural and morphological characteristics, but tailored for *a posteri* classification of reefs from remotely sensed satellite data for the specific purpose of global coral reef habitat mapping. This version 1 classification attempts to provide some standardization and harmonization of reef knowledge from across different disciplinary perspectives (ecology, geomorphology, remote sensing), and the number and type of classes and their descriptions reflect this merging of traditional local and global technological approaches. Classes were developed with end-users in mind, specifically coral reef science and conservation practitioners.

The Reef Cover classification is based on standard classification principals [1-4], with a special focus on addressing common issues with semantic interoperability that can prevent maps being effectively used by coral reef scientists and conservation practitioners [5]. Steps taken to improve interoperability include:

- Building on existing reef classifications and choosing commonly used terms ("Standard Name" describes the most parsimonious class name) and definitions ("Standard Label" provides a simplified text description to accompany maps)
- Provision of more detailed class description ("Standard Description") to supplement each definition. These longer descriptions summarize the 1-5 major diagnostic **environmental attributes** of the class, natural variability in the class and how they might be interpreted, and provides other contextual information regarding geomorphology, geography, ecology and **remote sensing** as well as comments about the possible relevance of the class to users.
- Listed commonly used synonyms and cross-walking between established reef mapping and monitoring projects (with citable references and a Cross-Walk table provided at the end)
- Translation into languages that cover the political jurisdictions of 60% of reef areas (e.g. Bahasa because Indonesia hosts 17% of all reefs, French because 14% of reefs fall into French-speaking territories [6]) to encourage broader use and feedback
- Simple diagram to illustrate where these classes might typically be found across four morphologically different reef types: fringing, barrier, atoll and platform reefs (see Glossary for

Reef Cover classes are listed in logical order from external seaward-facing through to internal reef structural features. While the shape and size of coral reefs varies widely between reefs, this level of detail appropriate as it reflects broadly some of the main internal structural classes commonly found in reefs across most biogeographic regions. Moreover, geomorphic classes like these have been shown to be good predictors of biological habitat richness and diversity [7].

As with any classification, the Reef Cover classes listed here is an approximation of reality and can never fully represent the full diversity of natural features presented by coral reefs. This classification represents a first step in supporting development and use of a new breed of dynamic habitat map [8], and will hopefully be further refined with input from the community and as technological advances allow for expansion of finer-scale mapping methodologies. We hope Reef Cover will be tool to assist in knowledge sharing at regional to global scales, acknowledging that there are many other ways of viewing or understanding coral reefs and their complex features.

### Reef Cover Classes


| Standard Name | Standard Label | Page |
|---|---|---|
| Reef Slope | Reef Slope is a submerged sloping lower **Fore Reef** area, beginning below the natural break in reef profile or, if no break exists, beyond 18 m, and extending seaward. | 5 |
| Sheltered Slope | Sheltered Slope is any **Reef Slope** (submerged sloping area extending into **Deep Water**) more protected from strong directional prevailing wind or current, either by land or by opposing reef structures, than the rest of the reef. | 7 |
| Reef Front | Reef Front is a submerged, sloping area extending seaward from the **Reef Crest** (or **Reef Flat**) towards the shelf break. | 8 |
| Sheltered Front | Sheltered Reef Front is any **Reef Front** with an aspect protected from strong directional prevailing wind or current, either by land or by opposing reef structures. | 9 |
| Terrace | Terrace is any seaward-facing **Fore Reef** feature with a shallow to near horizontal slope angle. Terraces often accumulate sand. | 10 |
| Wall | Wall is any seaward-facing **Fore Reef** feature with a near vertical slope. | 11 |
| Sheltered Wall | Sheltered Wall is **Wall** with an aspect protected from strong directional prevailing wind or current, either by land or by opposing reef structures. | 12 |
| Reef Crest | Reef Crest is a shallow zone characterised by highest wave energy absorbance, marking the boundary between the **Reef Flat** and the **Reef Front**. | 13 |
| Outer Reef Flat | Adjacent to the seaward edge of the reef, Outer Reef Flat is a levelled (near horizontal) broad and shallow carbonate platform, displaying distinct wave-driven zonation. | 14 |
| Inner Reef Flat | Inner Reef Flat is a low energy, sediment-dominated horizontal and generally shallow to gently sloping platform behind the **Outer Reef Flat** | 15 |
| Back Reef Slope | Back Reef Slope is a complex interior **Back Reef** zone occurring behind the **Reef Flat**. Of variable depth, it is sheltered, gently sloping, sediment-dominated and often punctuated by coral outcrops. | 16 |
| Lagoon | Lagoon is any sheltered broad body of water semi-enclosed by reef, with a flat, deep bottom (but depth shallower than surrounding ocean) dominated by soft sediment. | 17 |
| Shallow Lagoon | Shallow Lagoon is any sheltered, shallow (shallower than 5 m approximately), flat-bottomed sediment-dominated lagoon-like water body. | 18 |
| Plateau | Plateau is any submerged (deeper than **Reef Flat**), hard-bottomed, horizontal to gently sloping seaward facing reef platform. | 19 |
| Patch Reef | Patch Reef is any small, detached to semi-detached lagoonal coral outcrop arising from a sheltered, sandy-bottomed area. | 20 |
| Small Reef | Small Reef is any detached (stand-alone) reef, surrounded by deep water and too small to exhibit a central depression or other clear geomorphic zonation besides a **Fore Reef**. | 21 |
| Reef Island | Reef Island is a supratidal structural coral reef feature consisting of low-lying accumulations of reef-derived material. | 22 |

### Reef Slope

#### Reef Slope is a submerged sloping lower **Fore Reef** area, beginning below the natural break in reef profile or, if no break exists, beyond 18 m, and extending seaward. Windward facing, or any direction if no dominant prevailing wind or current exists.

Reef Slope (a term sometimes homonymously used to describe the entire **Fore Reef** area) refers to the deeper section of the exposed lower **Fore Reef** [9, 10]. Its main features are that it is 1) **deeply submerged**, 2) **sloping** and 3) **seaward-facing**. **Deeply submerged** or **subtidal**, this zone begins below the natural break in reef profile or beyond 18 m if none exists, and may descend beyond the shelf break into oceanic water, encompassing a large vertical area including the transition from the neritic shelf waters to the deeper ocean [11]. This depth means Reef Slope will eventually exceed the limit of features visible in optical imagery so will always **neighbour Deep Water**. **Seaward-facing** means this is an exterior reef zone, opening up into open ocean and facing away from land or lagoons [12, 13]. Reef Slopes are always **sloping**, but to varying degrees: atolls have straighter, steeper profiles; barriers tend to be subdivided into bumps or "terraces", and fringing reef profiles are gentler [9, 14-16]. This **steeply sloping** (**slope angle may be between 20°** and **45°**) quality means Reef Slope zones are subject to gradients of light and water movement, resulting in strong depth-driven patterns of coral community zonation [15]. Below 30 m Reef Slopes host mesophotic communities [17]. Reef Slope can be distinguished from **Reef Front** from space, as a visible **colour change** reflecting a change in slope angle [18].

Reef Slope zones will be of interest to scientists as their **depth** and **exposure** make them less accessible and less well studied than **Reef Fronts** [19]. They are often missing from maps as their planar extent is limited (often only extending 50 m in width from shallowest to deepest parts). However they may drop over several kilometers over which environmental gradients and coral communities change dramatically [19], expanding the amount of known reef habitat [20]. Below the strong influence of wave action these deeper zones may host greater coral diversity [10], and may be also provide potential refugia for corals from disturbances such as ocean warming and cyclones [21] (but see [17, 22]).

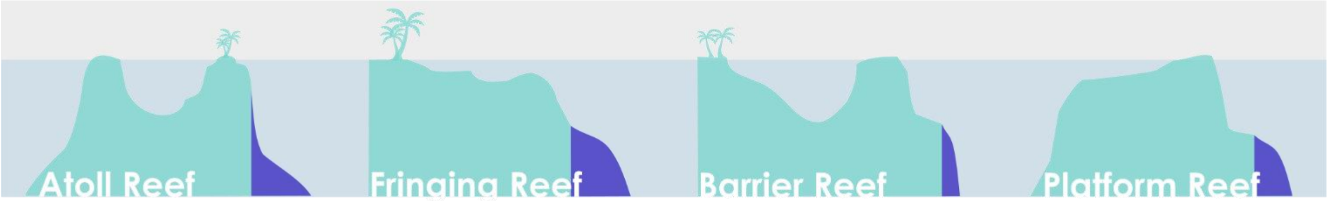


Reef Slope • Terumbu depan • Pendiente arrecifal frontal • Pente externe • الشعب منحدر المرجاني

*Drop Off • Escarpment • Seaward Slope • Outer Reef • Fore Reef subzone • Outer Reef Margin • Deep Reef Slope • Outer Fore Reef • Windward Slope • Exposed Slope*

### Sheltered Reef Slope

Sheltered Reef Slope is a **Reef Slope** (submerged sloping area extending into **Deep Water**) protected from strong directional prevailing wind or current, either by land or by opposing reef structures.

Sheltered Reef Slope classes are context dependent (i.e. no established thresholds exist for what constitutes “exposed” in terms of wind speeds, current velocities or wave energy), and so these classes will only usually occur on the leeward sides of **Atoll** and **Platform Reefs** where a pronounced exposure gradient exists around the reef. Where exposure is not pronounced, or prevailing wind direction fluctuates throughout the year, all slopes should default to being classed as **Reef Slope** rather than Sheltered Reef Slope.

Sheltered Reef Slope is therefore defined here as any **Reef Slope** (**submerged**/**subtidal**, **sloping** area extending into **Deep Water** – i.e. **seaward facing** exterior rather than interior reef zone) that is **protected** either by opposing reef structures or by an island from strong directional prevailing wind or currents most of the time. See Table 1 (Attribute Table) to see how Sheltered Reef Slope differs from **Reef Slope**, **Reef Front** and **Sheltered Reef Front**.

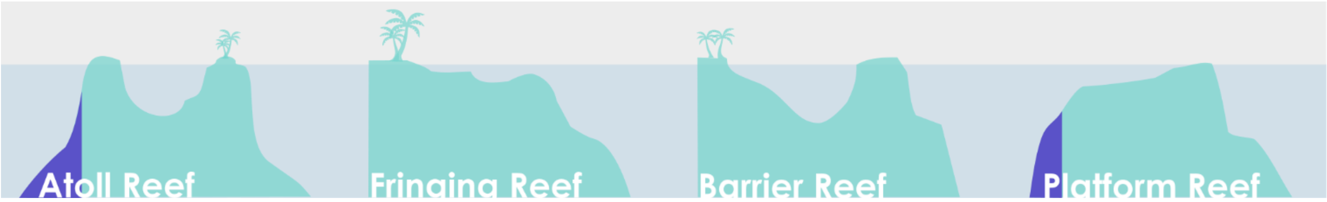


Sheltered Reef Slope • Terumbu depan terlindung • Pendiente arrecifal frontal protegidos • Pente externe abrité • المرجاني الشعب منحدر المحميه

*Leeward Slope* • *Protected Slope* • *Sheltered Slope*

### Reef Front

Reef Front is a submerged, sloping area extending seaward from the **Reef Crest** (or **Reef Flat**) towards the shelf break. Windward facing, or any direction if no dominant prevailing wind or current exists.

**Seaward facing**, **steeply sloping** (slope angle 20° - 45°) **shallow** upper **Fore Reef** zone (sitting above the natural break in **Fore Reef** profile, or above 18 m if no break exists), either windward facing or any direction if no dominant prevailing wind or current exists. Reef Front is the principal zone around which many other geomorphic zones are arranged [14]. Reef Front is always **subtidal** but shallower than **Reef Slope** [10] and this, along with being seaward-facing (opening up into open sea or ocean / facing away from land or lagoons) [2, 7] means they are characterised by **high wave exposure** [7] [13]. Reef Fronts often have steep inclines [13] but vary on a continuum of being fairly flat to more undulating spur and groove formations [8-11]). Reef Front zones are subject to strong gradients of both light and water movement, resulting in depth-driven patterns of coral community zonation [9]. Zonation may be more strong than on **Reef Slopes**, but more subtle than on **Reef Flats** that are subject mainly to just one environmental gradient (i.e. water exposure because uniform light) [15]. Other characteristics of Reef Front include being **proximal to the Reef Crest** [23].

High abundance of corals and fish, combined with accessibility to divers and snorkelers (due to their shallow nature) mean Reef Fronts are often targets for ecological monitoring programs, and are generally popular tourism spots.

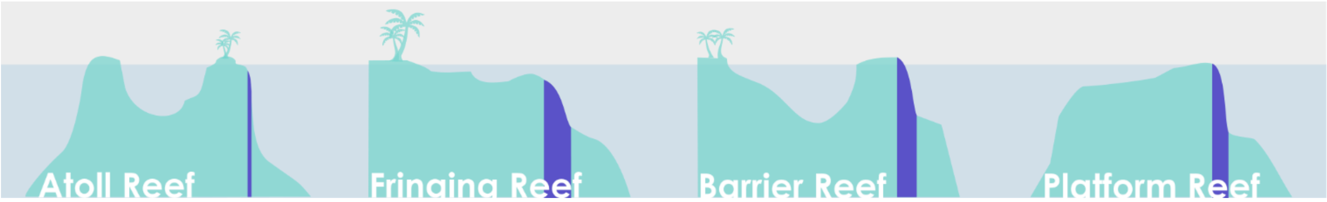


Reef Front • Terembu depan • Arrecifes delanteros • Front récifal • المرجاني الشعب واجهة

*Fore Reef • Reef Face • Foreslope • Shallow Zone • Spur and Groove Zone • Reef Slope • Windward Front • Exposed Front*

### Sheltered Reef Front

Sheltered Reef Front is any **Reef Front** with an aspect protected from strong directional prevailing wind or current, either by land or by opposing reef structures.

Sheltered Reef Front is any leeward **Reef Front** (**shallow**, **submerged**, **sloping** area extending **seaward** away from the **Reef Crest** or **Reef Flat**) that is comparatively more **protected (**either by land, and island or by opposing reef structures from strong directional prevailing winds or currents) than the rest of the **Reef Front**.

Similar to **Sheltered Reef Slope**, the Sheltered Reef Front class is highly context dependent (i.e. no established thresholds exist for what constitutes “exposed” in terms of wind speeds, current velocities or wave energy). This class will likely only usually occur on the leeward sides of **Atoll** and **Platform Reefs** where a pronounced exposure gradient exists around the reef. Where exposure is not pronounced, or prevailing wind direction fluctuates throughout the year, all slopes should default to being classed as **Reef Front** rather than Sheltered Reef Front.

Wave energy is one of the most important physical variables structuring coral reef communities. Geomorphologists frequently sub-divide **Fore Reef** zones based on relative exposure: e.g. from *Leeward* vs *Windward* to up to six types of breaker zones defined by Geister (1977) [24]. For coral reef scientists, Sheltered Reef Front zones are useful to distinguish from **Reef Fronts**, as long term exposure can greatly influence benthic community development, and also affect **Reef Front** profile shape (e.g. geomorphology) [9, 19, 25].

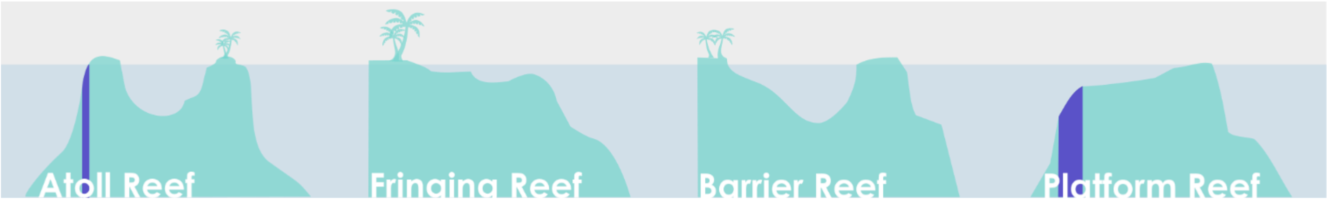


Sheltered Front • Terumbu depan terlindung • Arrecifes delanteros protegidos • Pente externe abrité • المرجاني الشعب واجهة المحميه

*Leeward Front • Sheltered Fore Reef • Reef Flank • Leeward Flank • Protected Fore Reef*

### Terrace

Terrace is any seaward-facing **Fore Reef** feature with a shallow to near horizontal slope angle. Terraces often accumulate sand and rubble.

Terrace is the most **horizontal** section of the **seaward-facing** **Fore Reef** zone, characterized by a **shallow slope angle**, and frequently acting as a division between the **Reef Front** and deeper **Reef Slope** [26]. Terraces represent any **Fore Reef** area with a gentle gradient (generally < 20° to near horizontal). They often **occur around 14 – 18 m depth**, and **neighbour** **Reef Front** and **Reef Slope**. Due to their gently sloping nature, Terraces often accumulate sand and coral detritus [18, 26]. Terrace is not a fundamental geomorphic zone: not all **Fore Reefs** exhibit discontinuities in the reef profile [27]; and terraces tend to be less common on steep-sided **Atolls** and more commonly found in **Barrier Reefs**.

Shallow reef Terraces can be distinguished from the **Reef Front** from satellite imagery, as a visible **colour change** reflecting the change in slope angle [18].

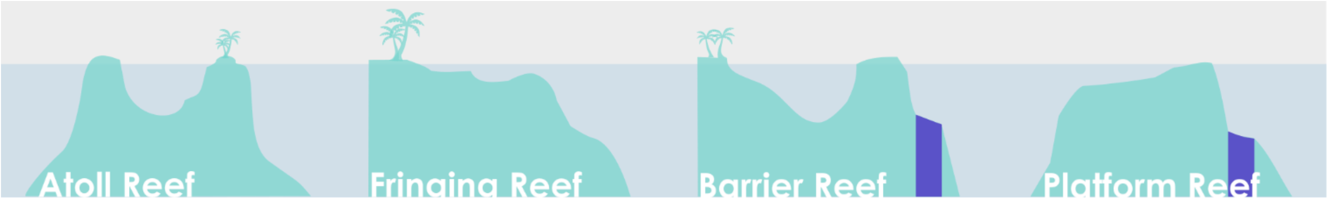


Terrace • Teras terumbu • Terraza de arrecife • Terrasse de récif • مدرّج

*Terrace • Sand Flat • Fore Reef Flat • Gentle Slope • Ten fathom terrace • Low gradient sand terrace • Windward Platform*

### Wall

#### Wall includes any seaward-facing **Fore Reef** feature with a near vertical slope.

Walls describe **near vertical** sections of **Fore Reef** [28], with the main feature distinguishing them from other **Fore Reef** zones being the slope angle, in contrast to "**Reef Slopes**" and "**Terraces**". Other defining features as **Fore Reef** features include being **seaward-facing**, **subtidal** and **steep**. Walls are not frequently ascribed an angle, instead being described as "**near-vertical**". Despite there not being much specific published guidance on what angle a gentle slope becomes a steep slope and when a steep slope becomes a wall, it has been suggested that **Reef Slope** is "any area of the reef with an incline of between 0 and 45 degrees" [23], and so a Wall may have an **angle > 45°**. Walls steep slope angle affects incident light and (along with exposure) may make it difficult for some benthic species to establish, meaning often supports soft coral dominated communities. Moreover, Walls exhibit strong depth/light gradients which will influence community composition, but may not exhibit a strong demarcation point unless accompanied by a change in slope angle (then move to another class).

The vertical nature of Walls mean these habitats are not well captured by planar mapping exercises, which cannot effectively represent the detail in these 3D vertical reef features. Walls are important draws for tourists as the near vertical faces and high water clarity may bring divers into close contact with large pelagic species, resulting in greater tourism and conservation value [29]. As with steep **Reef Fronts**, Marine Parks that incorporate Walls into their design will preserve more habitat per unit area, due to their vertical nature.

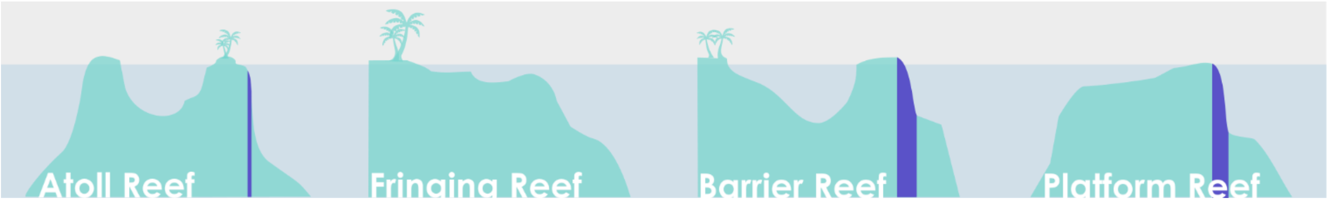


Wall • Tubir • Pared arrecifal • Tombant récifal • عمودية مرجانية شعب

*Reef Wall • Drop Off • Face • Cliff • Escarpment • Scarp Face • Steep Reef Slope • Vertical Rampart • Outer Rampart*

### Sheltered Wall

Sheltered Wall includes any seaward-facing **Fore Reef** feature with a near vertical slope, and aspect protected from strong directional prevailing wind or current, either by land or by opposing reef structures.

Sheltered Wall is any **Wall** (**near vertical, subtidal** **Fore Reef** feature) that is comparatively **protected** either by land, an island or by opposing reef structures from strong directional prevailing winds or currents than other **Fore Reef** areas.

Most Walls are associated with seaward facing highly exposed slopes: this Sheltered Wall class will likely only usually occur on the leeward sides of **Atoll** reefs where Walls are common and a pronounced exposure gradient exists around the reef. Sheltered Wall is highly context dependent (i.e. no established thresholds exist for what constitutes "sheltered" in terms of wind speeds, current velocities or wave energy). Where exposure is not pronounced, or prevailing wind direction fluctuates throughout the year, all Walls should default to being classed as **Wall** rather than Sheltered Wall.

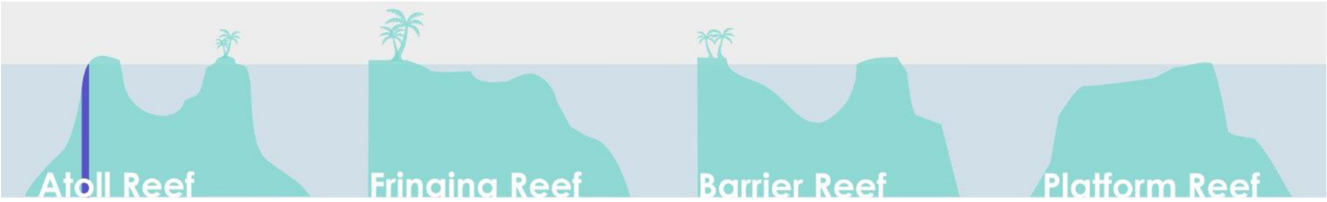


Sheltered Wall • Tubir terlindung • Pared arrecifal protegidos • Tombant récifal abrité • مرجانية شعب المحميه عمودية

*Sheltered Reef Wall • Drop Off • Leeward Face • Cliff • Sheltered Escarpment • Protected Outer Rampart*

### Reef Crest

#### Reef Crest is a shallow zone characterised by highest wave energy absorbance, marking the boundary between the Reef Flat and the **Reef Front**.

Reef Crest represents the break point at which a sharply defined edge divides the shallower reef platform from a more steeply shelving **Reef Front** [18]. It is arguably the most defining feature of any reef, as other geomorphic zones tend to be arranged in parallel to the crest [27]. Two characteristics of Reef Crest zones are that they represent a **demarcation point** separating the **Reef Front** from the **Reef Flat** [14, 23, 30], and are "an area of maximum wave shoaling", i.e. a zone that **absorbs the greatest wave energy**, playing a key role in coastal defense [23]. Reef Crest zones dissipate >85% of the incoming ocean wave energy and 70% of the swell energy [31, 32]. Darwin also recognised them as an area of maximal carbonate accretion; Reef Crests are the "growing point" of most reefs. Reef Crest zones are also sometimes described as the **shallowest** and often **emergent** part of the reef [23]. These types of Reef Crests are common on **Atolls**, where they appear "**flattened**" [11] or "**flat-topped**" [12], and are usually dominated by coralline algae in the breaker zone (Reef Rim), with live corals developing under the water surface in areas less likely to experience long periods of emersion. However the Reef Crest area of maximal energy absorbance can be found much **deeper** and is often "**gently sloping**" [18], curving at the boundary between horizontal reef flat and vertical slope. This is more common on **Fringing Reefs**, where Reef Crest development is linked to sea level: here the demarcation point between flat and slope is likely to be submerged, deeper than the **Reef Flat**, and gently curving, representing a less sharply defined transition. Coralline algae may be less of a dominant feature here. Some younger fringing reef types lack a clear Reef Crest altogether, despite the presence of a crest being one of the oft-cited defining elements of a **Fringing Reef** [30].

Reef Crest is of interest to scientists due to its accretionary growth potential and the role it plays in coastal protection, both of important in the context of climate change and associated predicted storm activity and sea level rise [33].

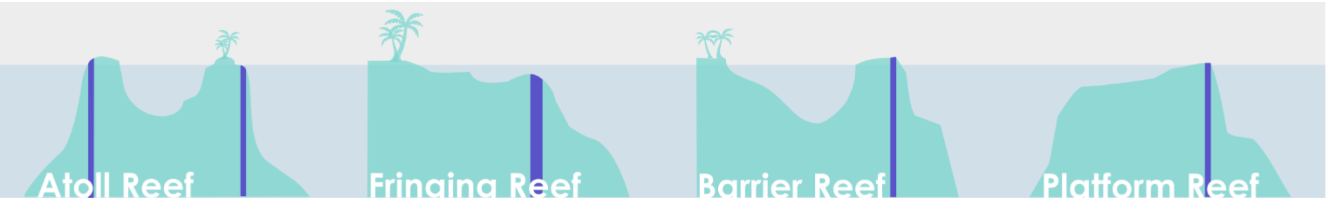


Reef Crest • Igir terumbu • Cresta arrecifal • Crête récifale • قمة الشعب المرجاني

*Surf Zone • Breaker Zone • Reef Edge • Reef Rim • Reef Margin • Rim Margin • Hardline Perimeter*

### Outer Reef Flat

#### Adjacent to the seaward edge of the reef, Outer Reef Flat is a levelled (near horizontal) broad and shallow carbonate platform, displaying distinct wave-driven zonation.

Extending inwards from the seaward edge of the reef, Outer Reef Flats are characterised by **broad** (hundreds of meters wide), **level surfaces** and a **shallow depth** (generally no deeper than a few centimeters to a few meters [10, 18]). In some regions – particularly the Pacific - they can be intertidal. The shallow and flat characteristics of this zone are a consequence of their geological development: upward growth of reefs is halted by the sea surface and so a growing reef platform can only really expand outwards. **Reef Flat** internal **zonation is particularly distinct** [15, 34], due to wave energy being the single driver (relatively uniform exposure to light). Belt-like sub-zones (e.g., algal rim, coralgal flat and coral windrows) are often detectable in aerial images of Outer Reef Flats as **coloured bands** [35]. Features such as deeper pockets (troughs or moats) where corals persist, scattered microatolls, “feo” (undercut mushroom shaped rocks remnant from where sea level has fallen) and stranded fossil ridges can punctuate Outer Reef Flat zones.

Outer Reef Flat is a sub-feature of **Reef Flat**. With one edge located **parallel** to and **neighbouring the seaward edge** of reef, Outer Reef Flat zones are also distinguished from interior **Inner Reef Flats** by depth (**shallower**); slope angle (**flatter**, but may slope gently down towards Inner Reef Flat); exposure (more **high energy waves** and more likely to experience emersion); benthos (**harder**, with more corals and coralline algae in the case of coral-dominated flats or larger pieces of scattered debris on rubble-dominated flats), and **zonation** (e.g., clearer zonation due to sharper exposure gradient, with coral cover and or rubble size diminishing away from the seaward edge). Although the Outer Reef Flat receives protection from the **neighbouring** **Reef Crest** which absorbs most (85%) of the wave energy, the outer 150 m of **Reef Flat** will dissipate 65% of the remaining wave energy and reduce wave height by a further 43% [31]. **Outer Reef Flat** zones may contribute disproportionately to global reef extent calculations, making up the bulk of reef area (Reef Top) from an aerial perspective.

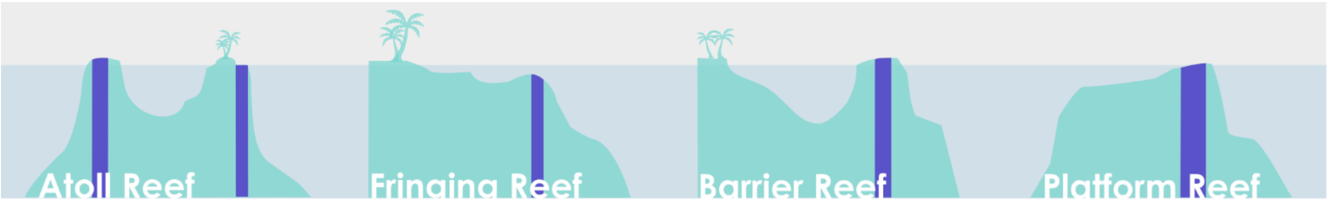


Outer Reef Flat • Rataan terumbu luar • Arrecife plano exterior • Exterieure du platier récifal • خارجي مرجاني مسطح

*Reef Top • Inter-Reef Tract • Coralgal Flat • Outer Living Coral Zone • Coral Windrows*

### Inner Reef Flat

#### Inner Reef Flat is a low energy, sediment-dominated horizontal to gently sloping platform behind the **Outer Reef Flat**

Inner Reef Flat lies inwards of the **Outer Reef Flat** and shares many characteristics in terms of being **shallow**, **horizontal** and **broad [35]**. The leeward "sand zone" of a **Reef Flat**, Inner Reef Flats are highly depositional systems **dominated by sediment**. They are distinguished from the **Outer Reef Flat** by depth (often slightly **deeper**), slope (still flat but with a greater downward gradient), exposure (**low energy** leading to calmer conditions and more sand) and benthos (**softer substrates**). Additionally, Inner Reef Flat has a relative paucity of living coral and algae compared to coral-dominated **Outer Reef Flats** (and smaller rubble pieces on rubble-dominated **Outer Reef Flats**). Internal zonation is also less pronounced with material stratified across wide Inner Reef Flats. These differing benthic features often mean Inner Reef Flat zones appear distinct in colour from Outer Reef Flat zones from aerial imagery, despite underlying geomorphology being broadly similar. Inner Reef Flat will **neighbour** **Back Reef Slope**, **Lagoon/Shallow Lagoon** or **Sheltered Reef Slope** depending on the reef type [36]. Where Inner Reef Flat meets **Shallow Lagoon**, it can be distinguished by its shallower depth, more horizontal angle, lower sand content and more complex features, or unfused coral windrows grading into coral patches and sand flats if neighbouring a coral-dominated **Outer Reef Flat** (or rubble pieces if neighbouring a rubble-dominated **Outer Reef Flat**) [34].

Inner Reef Flat areas may often constitute the broadest extent class, although often excluded from reef extent (Reef Top) calculations due to depth and sandy composition. Additional Inner Reef Flat features include a low cover and diversity of corals due to the harsh conditions and limited opportunity for upward growth [9], which means they are often described as "barren" [18]. Despite this, they are a significant habitat for many organisms, supporting high abundance and biomass of herbivorous fish, for example [37]. **Reef Flat** descriptions also frequently reference the presence of rubble, describing flats as "strewn with detritus", and "littered with coral sand and storm tossed shine or blocks of reef limestone brought in from outer margin" [38].

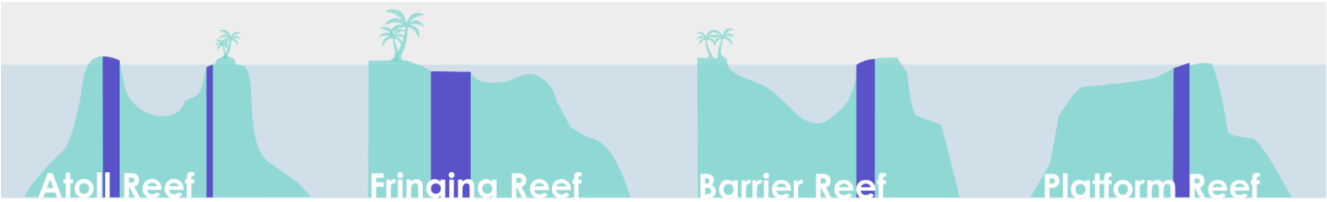


Inner Reef Flat • Rataan terumbu dalam • Arrecife plano interior • Extérieure du platier récifal • داخلي مرجاني مسطح

*Sand Flat • Sand Zone • Leeward Reef Flat • Coral Patches • Unfused Coral Windrows*

### Back Reef Slope

Back Reef Slope is a complex interior **Back Reef** zone occurring behind the **Reef Flat**. Of variable depth, it is sheltered, gently sloping, sediment-dominated and often punctuated by coral outcrops.

Back Reef Slope is a specific **Back Reef** feature. **Back Reef** is a widely used term (see Glossary), which can include **Reef Flats** and **Lagoons**. Back Reef Slope describes the transitional landward **gently sloping** area that links the **Reef Flat** to **Lagoon** (or to shelf areas in the case of **Platform Reefs)** [10]. It is an **interior-reef zone** (i.e. always found behind the **Reef Crest** or flat) and characteristically **sheltered**. Being a low energy area means the Back Reef Slope area (like all **Back Reef** areas - including **Lagoons** and **Reef Flats**) is a largely **depositional environment**, receiving debris swept landwards from the **Reef Crest** and **Fore Reef** – from rubble strewn across reef flats, to sand sheets and mud deposits into lagoons [12, 27]– meaning this sloping area will generally be **sediment-dominated**. Back Reef Slopes are variable in terms of slope angle and depth, but often are punctuated by coral outcrops, including **Patch Reefs**, particularly deeper ones.

Back Reef Slopes **shallow depth, sandy substrate** and distance from the **Fore Reef** often make them important anchorages, and their sheltered nature and abundant coral outcrops can make them popular dive sites too.

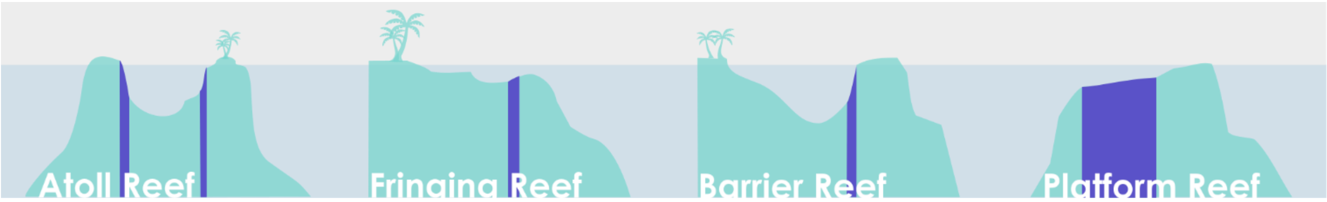


Back Reef Slope • Lereng terumbu belakang • Pendiente de arrecife posterior • Pente récifale interne • الخلفي الشعب منحدر

*Open Complex Lagoon • Subtidal Reef Flat • Lagoon Reef Slope • Back Reef • Escarpment • Back Barrier • Sediment Apron*

### Lagoon

#### Lagoon is any sheltered broad body of water semi-enclosed by reef, with a flat, deep bottom (but depth shallower than surrounding ocean) dominated by soft sediment.

Lagoons are **sheltered** internal water bodies protected from wave energy by a **Reef Crest** and **Reef Flat** by which it is semi-enclosed, resulting in a calm and stable environment [13]. Lagoons are deep, but depths are variable: while 10 m [30] and 5 m [39] are suggested depth thresholds for distinguishing a "true" Lagoon (a defining feature associated with **Barrier Reefs** and **Atolls** only) from shallower water bodies (e.g. associated with all reef types, including **Fringing Reefs** and **Platforms**) atoll lagoon floors are typically 20-36 m [40] and can reach >70 m [41]. Here, we suggest a **threshold of 5 m** for Lagoon depth, but as a more general rule, lagoon depth is always **shallower than the neighbouring water body** on the reef's seaward side [11]. Lagoons tend to be **broad in width** although width and shape can be variable depending on reef type (e.g., **Atoll** lagoons more rounded, **Barrier Reef** lagoons elongated).

Lagoons are always highly **depositional environments**, receiving near-continuous supply of calcium carbonate bioeroded from **Reef Flats**, **Crests** and **Fore Reefs**. This means lagoon floors are largely **soft bottomed**, **dominated by sediment** of biological origin, and constantly infilling [41]. **Enclosure to semi-enclosure** within a bordering reef construction **limits water exchange with the sea** creating a relatively **calm environment**, typified by low currents [42]), long residence times (up to 12 days), and seawater supersaturated with calcium carbonate [41] which can promote coral growth [18]. As a result, Lagoons often feature beds of coral (see **Patch Reef**). While floor level remains constant, many lagoons develop small knolls (non-living coral heads and pinnacles) as well as deep sinkholes caused by karst solution effects.

Their sheltered nature and accessibility means lagoons are often of significant cultural and economic importance, as a natural harbour, recreational and fishing area [43].

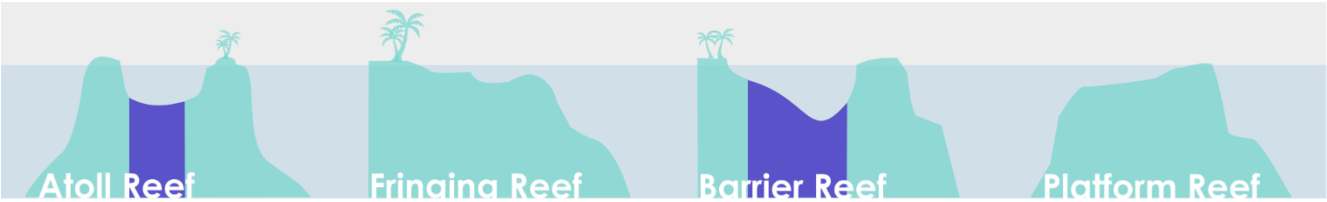


Lagoon • Laguna (dalam) • Laguna (profunda) • Lagon (profound) • بحيرة

*Blue Lagoon* • *Deep Lagoon*

### Shallow Lagoon

#### Shallow Lagoon is any sheltered, shallow (shallower than 5 m approximately), flat-bottomed sediment-dominated lagoon-like water body.

Shallow Lagoon zones are characterised mainly by their **shallow depth** (no deeper than 5 m), but they also tend to be **soft bottomed**, feature fewer coral patches than true lagoons, and are **sand-dominated**. Unlike the **Lagoon** class, which describes true lagoons, Shallow Lagoons are too shallow to technically be lagoons, although they share many features. The type of sediment in a Shallow Lagoon is typically foraminiferal dominated sediment, which differs from the *Halimeda*-derived sediment in deeper atoll lagoons [41].

Shallow Lagoons can be found associated with any reef type, while technically, true lagoons (deep lagoon, or **Lagoon**) by definition are associated with **Barrier Reefs** and **Atolls** only [44], and not **Platform Reefs**, **Fringing Reefs** or other reef types. A threshold used to distinguish **Barrier Reefs** from **Fringing Reefs** in the literature is that the lagoon depth needs to exceed 10 m [30, 45]. However, this implies that **Fringing Reefs** can be associated with lagoon-like water bodies, with a depth <10m. There are exceptions to this rule where true **Lagoons** become very shallow; Lighthouse Reef in Belize has a 120 sq m sandy lagoon, just 2-6 m deep. Unlike true **Lagoons**, Shallow Lagoons do not need to be enclosed by reef.

Shallow Lagoons are often important as access channels for coastal communities [45], as they can cut through reef crests and flats and provide access to shorelines.

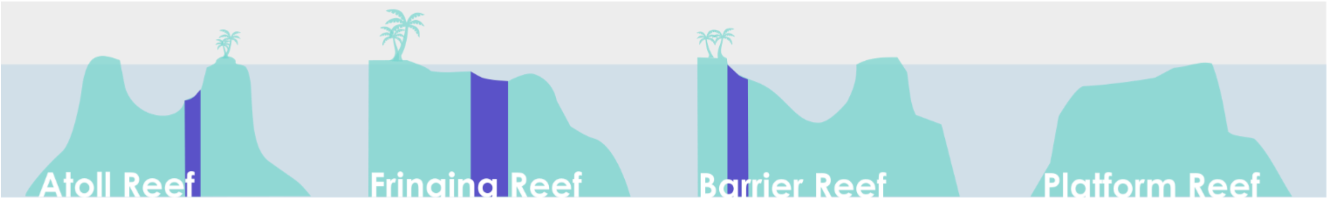


Shallow Lagoon • Laguna dangkal • Laguna somera / Laguna Pre-Arrecifal • Lagon peu profound • ضحلة بحيرة

*Shallow Water Body* • *Boat Channel* • *Pseudo-Lagoon* • *Lagoonlet* • *Miniature Lagoon* • *Back Reef Channel* • *Tidal Flat* • *Moat* • *Sand channel*

### Plateau

#### Plateau is any submerged (deeper than approx. 6 m), hard-bottomed, horizontal to gently sloping (angle less than 10°), seaward facing reef platform.

Including submerged offshore platforms, banks and shoals, the Plateau class refers to any **submerged** carbonate reef feature **beyond 6 m depth**, and is a feature rather than a standard geomorphic zone that occurs in all reefs. As well as being **submerged**, other characteristics are that Plateaus are largely **horizontal** (angle < 10 °), and are always found **adjacent to deeper water** (seaward facing) - in some cases it may be a standalone deep reef. Other features of this class are that the majority of the reef structure does not reach the surface, it cannot be subdivided into other geomorphic zones, and is made of **hard substrate**. Some of the world's largest reef structures – for example the 12,642 sq km Great Chagos Bank, and Cay Sal, the Bahamas third largest bank - are entirely submerged [8]. The "Plateau" map class refers exclusively to coral reefs - these reef features can be formed by other, non-scleractinian sources: *Halimeda* can also form submerged banks and biohermal structures [8].

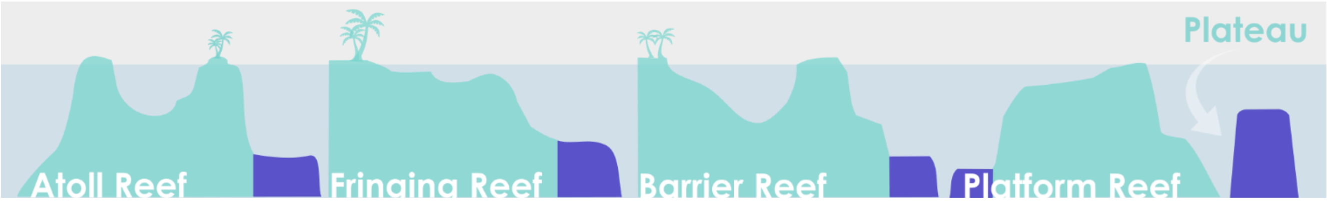


Plateau • Dataran tinggi • Meseta de arrecife • Plateau récifal (Terrasse) • المرجانية الشعاب هضبة

*Platform* • *Bank* • *Shelf* • *Shoal* • *Bank Shelf* • *Offshore Platform*

### Patch Reef

#### Patch Reef is any small, detached to semi-detached lagoonal coral outcrop arising from a sheltered, sandy-bottomed area.

A feature rather than an independent geomorphic zone, Patch Reefs are isolated reef outcroppings found in **sheltered** and **sandy areas** (e.g. **Lagoon**, **Back Reef Slope** and **Shallow Lagoon**) that may develop in close proximity to each other but are often physically separated by sandy lagoon floor. Patch Reefs are generally arranged independently of the broader geomorphic organisation of a reef (that is, along a structural axis relative to contours of shore or shelf edge) and to some degree **detached from other reef structures**: [11, 23] although their ability to aggregate and grow together into networks suggests "isolation" is not necessarily a characteristic that is easy to define. Patch Reefs will tend to have a **high ratio of vertical relief to planar reef** "with a vertical relief of one meter or more in relation to the surrounding seafloor" [11].

Patch Reefs are incredibly diverse in terms of size, distribution, shape and coalescence, meaning the class can be interpreted widely. A multitude of terms exist to describe different types of Patch Reefs – based on their **distribution** (e.g. "dense" or "diffuse"[23]); **morphology** ("single", "coalesced", "linear" and "reticulate"[46]); connectedness ("individual" or "aggregated"); **size** (defined by whether outcrops are large enough to be distinguished or too close together or small to be mapped independently [11]); **location** of the Patch Reef (splitting "shelf patch", "lagoon patch" and "intra seas patch" from "coastal/fringing patch" [47]), and relative **depth** (emergent "patches", submerged "knolls" and deep water "pinnacles" [18]. Large Patch Reefs (e.g., highly reticulated Patch Reefs, if broad enough, or Farus) will sometimes be zoned into geomorphic classes rather than as a Patch Reef.

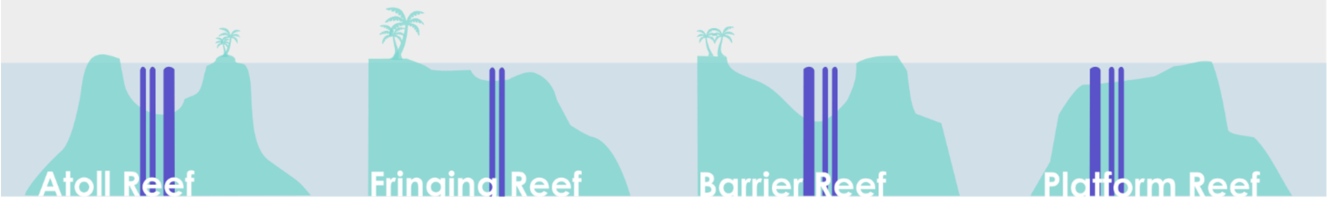


Patch Reef • Terumbu serpihan • Parche arrecifal • Massif corallien • رقعي شعب

*Lagoonal Reef • Mesh • Bommies • Coral Patches • Pinnacles • Knolls • Reticulate Reef • Coral Outcrops • Lagoon Reef*

### Small Reef

Small Reef refers to any detached (stand-alone) reef, surrounded by deep water and too small (generally ⪅ 1 sq km) to exhibit a central depression or other clear geomorphic zonation (e.g. **Reef Crest**, **Reef Flat**, **Back Reef**) besides a **Fore Reef**.

Size is the main defining feature of a Small Reef. Small Reefs are spatially independent reef outcrops, too small to lack the large defining features that would enable them to be confidently classified as either an **Atoll**, **Fringing**, **Barrier** or **Platform Reef**, and too shallow to be a submerged reef (**Plateau**). As well as being too "**small**" and "**featureless**" to be assigned a classical reef type (Hopley suggests small reefs can be "shelf reefs <1 km$^2$ growing off antecedent platforms"), Small Reefs are generally **rounded** or ovoid in shape [23], spatially **independent** (i.e. not a geomorphic sub-feature of a reef type, or connected to any other reef – therefore this class excludes lagoonal **Patch Reefs** and features like faros, that have developed inside of larger reef geomorphic structures), and largely **featureless** (e.g. lacking in strong geomorphic zonation - besides a **Reef Front** - that would allow the geomorphic zones to be clearly defined). This is because in order to develop a **Lagoon** and therefore strong zonation, **Platform Reefs** generally need to be at least 1 km across: since any reef flat >0.5 km in width is capable of causing rapid infilling and transition to a planar "platform" reef.

In the first ever attempted global coral reef map, Darwin's 1843 map of coral reef distribution, Darwin identified fringing reefs (in red), atolls (dark blue) and barrier reefs (pale blue), but commented that "there are many scattered reefs, of small size, represented in the chart by mere dots, which rise out of deep water: these cannot be arranged under either of the three classes". These Small Reefs may be often overlooked, but contribute a significant proportion to global coral reef habitat. Moreover, they often provide an important role in genetic connectivity of marine organisms, providing a stepping stone between reef features, and make popular fishing and diving spots.

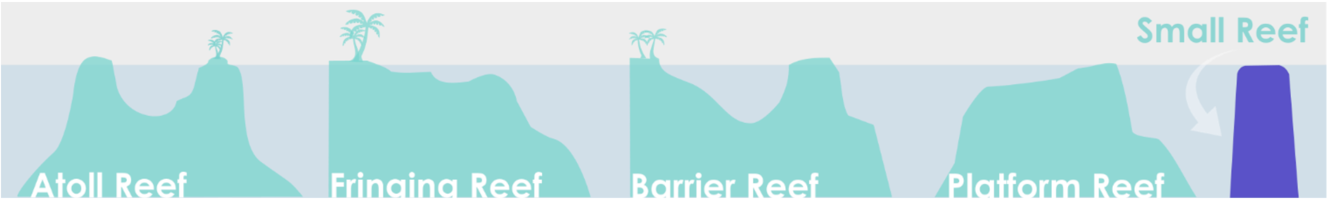


Small Reef • *Terumbu karang kecil* • *Arrecifes pequeños* • *Petit récif corallien* • شعب صغير

*Coral Knoll* • *Pinnacle Reef* • *Patch Reef* • *Marginal Structure*

### Reef Island

#### Reef Islands are supratidal structural coral reef features consisting of low-lying accumulations of reef-derived material.

Reef islands are an important geomorphological component of coral reef structures and as such should be distinguished from non-reefal shorelines in any reef mapping [3]. McLean and Woodroffe (1994) define Reef Islands as “low-lying (typically less than 5 m above sea level) accumulations of biogenically derived sediments” [48]. On **Platform** and **Barrier Reefs**, reef islands may consist of sand cays that form on top of shallow **Reef Flats**. **Atoll** islands are “commonly 50-100 m wide with a high ridge on the ocean side and a lower ridge on the lagoon side, a vegetated core and mobile beaches along both sides with cemented beach sand (beachrock) and cemented coral rubble (conglomerate)” [48]. Variability is usually in their composition – sand, shingle, coral rubble, beachrock - and whether the island is vegetated or unvegetated. Some classifications also sub-divide reef islands into zones – including intertidal area, mangroves and saltmarshes.

Reef Islands are commonly sub-divided based on size, shape, composition or whether they are vegetated [39, 48], or associated with atolls or not [49]. These differences are important in understanding island geomorphology and future change [50]. Reef Islands are critical for a range of socio-economic and ecological regions, including the provision of habitable land in low-lying countries (e.g., in the Maldives, reef islands provide a home for 516,000 people), nesting areas for turtles and seabirds, cultural and economic importance (for example islands of the Great Barrier Reef generate an estimated five billion in tourism alone) [50]. Concerns about stability and persistence of Reef Islands – particularly low lying coral atolls - in the face of anthropogenic climate change driven sea-level rise have made them an important area of research in recent decades.

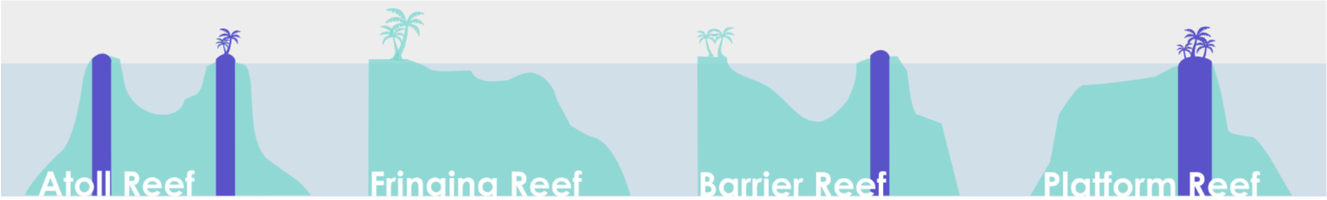


Reef Island • Pulau terumbu • Isla arrecifal • Complexe récifal d’îles • جزيرة مرجانية

Cay • Sand Cay • Motu

### Case study: the *Allen Coral Atlas*

#### Application of *Reef Cover* to a global reef habitat mapping exercise

The *Allen Coral Atlas* is a global-scale coral reef habitat mapping project that is using Planet Dove 3.7 m resolution daily satellite imagery (in combination with wave models and field data) to create consistent global coral reef habitat maps with the purpose of supporting science and conservation. The Global Geomorphic Zones mapped were based on Reef Cover but were adapted for the dataset.

The twelve reef geomorphic zones and features mapped by the *Allen Coral Atlas* are listed below. These zones known to be fairly consistent across different biogeographic regions, and zones are often associated with regionally distinct ecological assemblages of benthic animals and plants.

##### *Allen Coral Atlas* Geomorphic Classes

| | |
|---|---|
| Reef Slope* | **Reef Slope is a submerged, sloping area extending seaward from the Reef Crest (or Flat) towards the shelf break. Windward facing, or any direction if no dominant prevailing wind or current exists.** Reef Cover Class: *Reef Fron*t. |
| Sheltered Reef Slope* | **Sheltered Reef Slope is any submerged, sloping area extending into Deep Water but protected from strong directional prevailing wind or current, either by land or by opposing reef structures.** Reef Cover Class: *Sheltered Reef Front*. |
| Reef Crest | **Reef Crest is a zone marking the boundary between the Reef Flat and the Reef Slope, generally shallow and characterised by highest wave energy absorbance.** |
| Outer Reef Flat | **Adjacent to the seaward edge of the reef, Outer Reef Flat is a level (near horizontal), broad and shallow platform that displays strong wave-driven zonation.** |
| Inner Reef Flat | **Inner Reef Flat is a low energy, sediment-dominated, horizontal to gently sloping platform behind the Outer Reef Flat.** |

| | |
|---|---|
| Terrestrial Reef Flat* | **Terrestrial Reef Flat is a broad, flat, shallow to semi-exposed area fringing reef found directly attached to land at one side, and subject to freshwater run-off, nutrients and sediment.** No Reef Cover class. |
| Back Reef Slope | **Back Reef Slope is a complex, interior - often gently sloping - reef zone occurring behind the Reef Flat. Of variable depth (but deeper than Reef Flat and more sloped), it is sheltered, sediment-dominated and often punctuated by coral outcrops.** |
| Deep Lagoon* | **Deep Lagoon is any sheltered broad body of water, fully to semi-enclosed by reef, with a variable depth (but deeper than 5 m approx. and shallower than surrounding ocean) and a soft bottom dominated by reef-derived sediment.** Reef Cover Class: *Lagoon*. |
| Shallow Lagoon | **Shallow Lagoon is any fully to semi-enclosed, sheltered, flat-bottomed sediment-dominated lagoon area, shallower than 5 m approx.** |
| Plateau | **Plateau is any deeper submerged (> 5 m approx), hard-bottomed, horizontal to gently sloping (angle shallower than 10 ° approx), seaward facing reef platform.** |
| Patch Reef | **Patch Reef is any small, detached to semi-detached lagoonal coral outcrop arising from sandy bottomed area.** |
| Small Reef | **Small Reef refers to any detached (stand-alone) reef, surrounded by Deep Water and too small (generally less than approx. 1 sq km) to show a central depression and/or other clear geomorphic zonation (e.g. crest, flat, backreef) besides a Reef Slope.** |

**Table 1. Allen Coral Atlas global geomorphic map classes, as an example of how *Reef Cover* can be adapted to a specific mapping purpose. *Class modified from Reef Cover.**

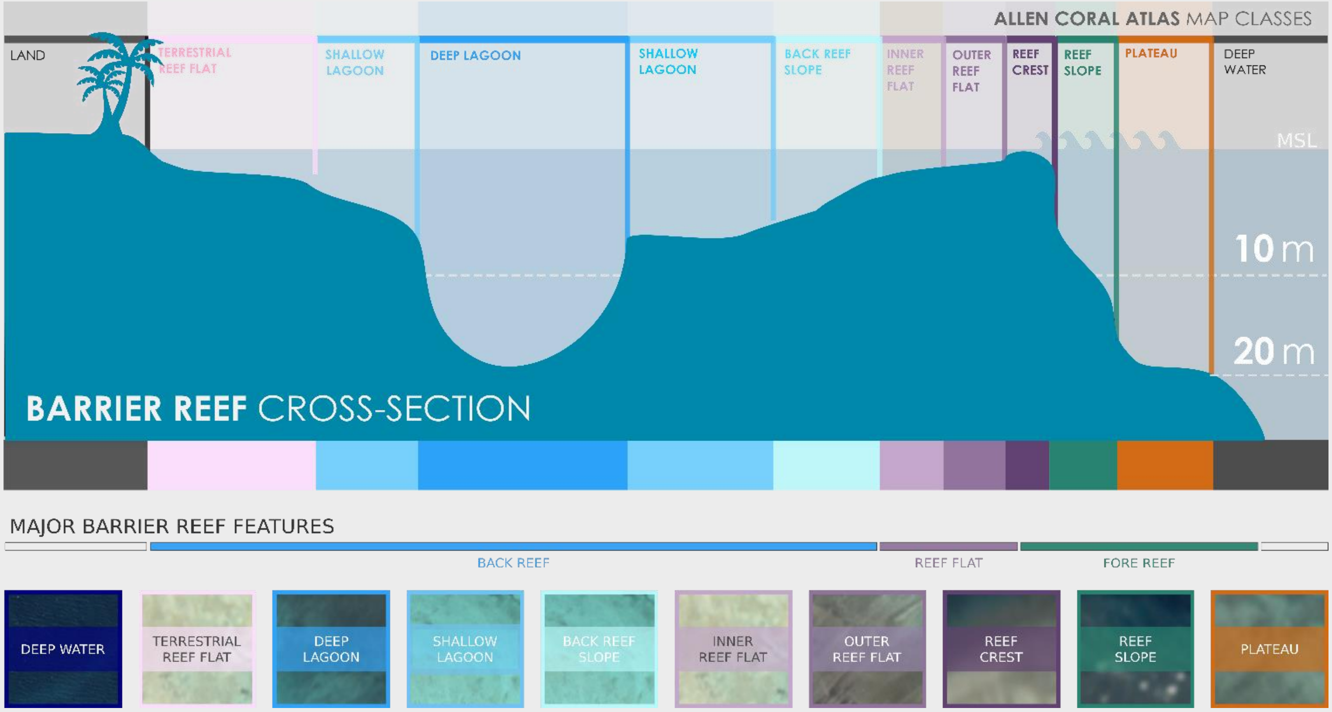


**Figure 1. Example of Allen Coral Atlas Global Geomorphic Zones as applied to a Barrier Reef.**

#### Benthic Map Classes

Benthic classification for a global scale coral reef mapping approach is still highly constrained by spatial resolution, quality and accessibility of biophysical earth observation data [2, 51, 52]. At the time of developing the *Reef Cover* classification, these data limitations (namely resolution of available remote sensing datasets and processing capabilities) prevented development of a scalable benthic classification framework that adequately addressed user needs and/or referenced the rich natural history knowledge of reef benthic diversity (usually assessed at the sub-meter scale). While many coral reef classification systems – whether for monitoring, mapping or management – include a benthic component (see Crosswalk table), the number of classes for global applications is often condensed to three-to-six and relevance to reef practitioners limited in comparison to the information collected from in-water benthic assessments.

Remote sensing is still fundamentally unable to distinguish some of the key measures that ecologists prefer to assess reef health – cover of living coral, cover of dead coral, cover of bleached corals and functional forms of algae [24], but broad classes like Coral Habitat and Rubble can still be very useful. For the *Allen Coral Atlas*, Planet Dove-derived spectral reflectance data provides some information about benthic composition, while bathymetry maps, slope angle and wave data (used to differentiate geomorphic zones) can be useful as surrogates for aspects of the physical environment (light availability, temperature, wave exposure) that determine most coral reef ecological partitioning. Underlying reef structure also helps determine gradients in light (depth and turbidity) and water movement (waves and currents) known to be the main drivers of benthic zonation on reefs [9]. This means global geomorphic maps can help influence benthic map classes.

The Allen Coral Atlas Global Benthic Mapping Classes described below were developed by Roelfsema *et al* [53] with input from other coral reef benthic classifications. This classification maximizes the breadth of information available from Planet Dove remote sensing data to create the best classes we can to support users. With rapidly developing technological and computing capabilities, we would hope to integrate a full benthic level to the *Reef Cover* classification at a later date, when data resolution improves to the degree where it can better relate to benthic field classifications (hundreds of coral taxa and morphologies at the sub-meter scale).

### *Allen Coral Atlas* Benthic Classes

#### Coral Habitat (Coral / Algae)

Coral Habitat is any hardbottom area supporting living coral and/or algae.

The Coral Habitat mapping class describes a habitat characterised by a **hard underlying framework** (usually coral-derived limestone framework, but may be non-carbonate) with a benthic **covering of coral** (including soft coral) and/or seaweeds (including macroalgae and turf algae). Coral Habitat is generally the most **topographically complex** class (sand, seagrass, rubble and rock are comparatively flat), supports the greatest amount of **animal diversity and biomass**, and most commonly associated with **Reef Front** and **Slope**, **Sheltered Reef Front** and **Slope**, **Patch Reefs** and **Outer Reef Flat** classes.

Coral abundance is a widely used proxy for coral reef health and an important metric for ecological monitoring of reefs. Most benthic classifications distinguish coral and algae habitat (e.g. [23]) and many move beyond this to classify coral morphology (e.g. branching, sheet, massive, encrusting [23]), identify dominant taxa (e.g., Palmata zone, Cervicornis zone, [26]), or estimate proportional cover (<10% cover, >50% cover [11, 36, 54]). The photosynthetic nature of both corals and seaweeds mean they are spectrally similar making them challenging to distinguish clearly though remote sensing [55]. The epilithic algal turf or film that quickly covers corals following death creates similar spectral signatures, meaning dead coral matrix cannot be reliably distinguished either, because of the speed at which it becomes covered.

Coral Habitat classes will generally have a cover of coral or algae of at least 1%, normally more than 5% and sometimes exceeding 40%, but does not necessarily have a dominance of any of these groups over non-living substrate. With average coral cover 10-20% globally, most reef habitats - even those supporting extensive coral growth - are unlikely to be quantitatively dominated by coral [23]). Other benthic classifications have pointed out that "*describing reefs with the highest coral cover as algal-dominated may be politically unacceptable and confuse interpretation*" [23] suggesting that systematic accuracy should be sacrificed to aid intuitive acceptance of a scheme.

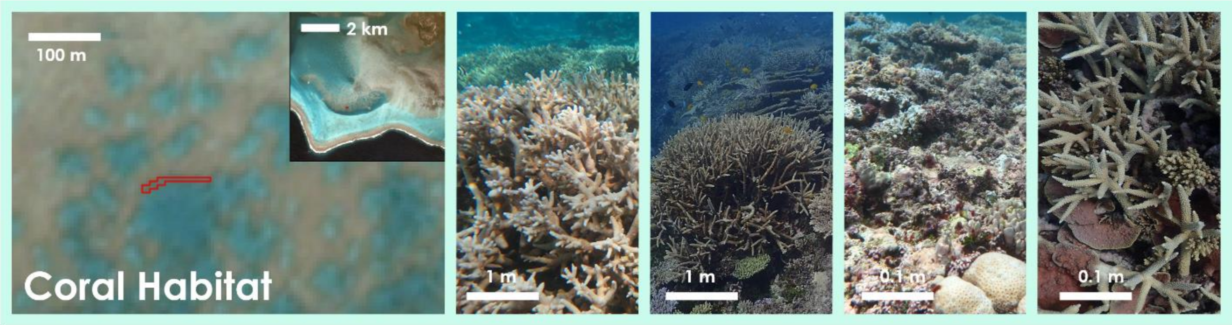


Coral Habitat • Habitat karang • Hábitats coralinos • Habitat corallien • مرجاني موطن

Coral / Algae • Coral dominated • Coral framework • Hardbottom • Hard Coral • Stony Coral • Live Coral • Coral Reef • Mixed coral • Coral • Coral field • Carbonate framework

#### Seagrass

Seagrass is any habitat where seagrass is the dominant biota.

The Seagrass class describes a **soft-bottomed** habitat dominated by any single species of **seagrass** from the order Alismatales (e.g. *Syringodium sp., Thalassia sp*., and *Halophila sp.*) or any combination of species [11]. Seagrasses can form extensive beds, called meadows, which shelter abundant diverse species (epiphytes, small invertebrates and juvenile fish), support herbivory (e.g. of turtles and dugongs) and play an important role in trapping sediment, as well as biogeochemical cycling. This class also includes sparser or more spatially restricted seagrasses, as long as it is the dominant biota, and/or has a total cover >10%. Seagrass habitats are most commonly associated with **Shallow Lagoon** and **Back Reef Slope**.

Other classifications further distinguish this class based on biomass, for example "high density" and "low density" or "sparse" seagrass [23, 28].

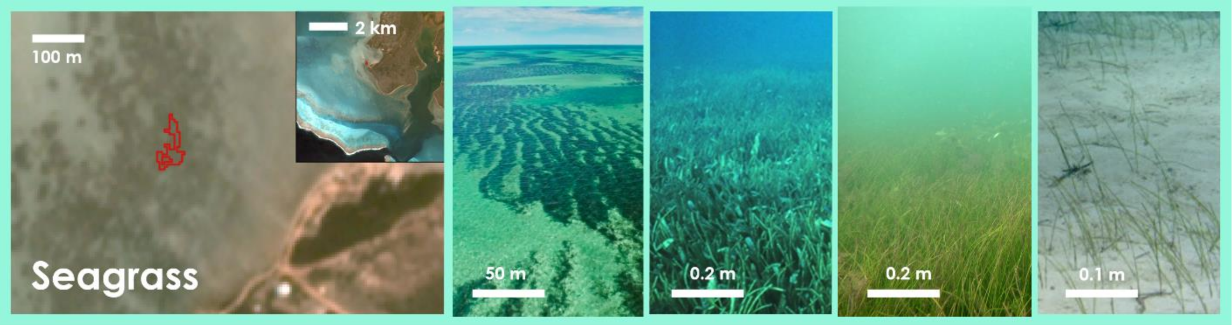


Seagrass • Padang/ hamparan lamun • Pastos marinos / ceibadales • Herbiers marins / gazons algaux • البحرية الأعشاب مرج

Seagrass Meadow • Seagrass Bed • Seagrass Dominated • Phanerogam Beds

### Microalgal mats

Microalgal Mats are visible accumulations of microscopic algae in sandy sediments.

The Microalgal Mat class describes microscopic communities - abundant and spatially extensive enough to be visible as mats - growing on or in the top few centimetres of **shallow, sandy sediments**. The benthic microalgae that comprise these mats, also known as microphytobenthos, are primarily diatoms, but include cyanobacteria, chlorophytes, and other microscopic organisms that grow on sand, silts and muds in both marine and freshwater habitats [56].

In shallow, sandy and sheltered reef areas, such as the leeside of islands and in **Lagoons**, benthic microalgae aggregate into mats which can be geographically extensive (up to several kilometers), and penetrate up to 15 cm into the sediment (although most biomass occurs in the upper centimetres). Benthic microalgal mats are productive habitats that play important roles in sediment stabilisation, trophodynamics and biogeochemical cycling [57]. They may promote benthic recovery by rapidly re-oxygenating the sediment surface. In ecology, patterns of herbivory can create grazing halos in the mats, the size and number of which can indicate ecosystem health [58]. These habitats are most often associated with sheltered **Back Reef** areas and **Shallow Lagoons [28]**.

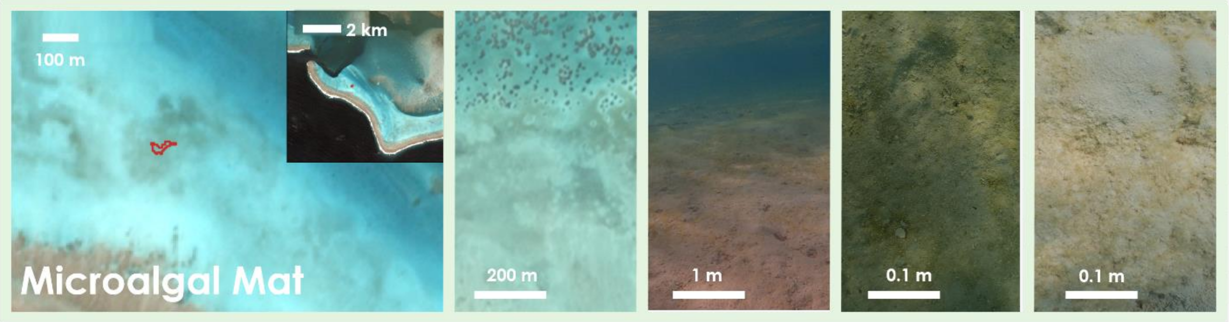

Microalgal Mat • Ganggang mikro bentik • Microalgas bénticas • Micro-algues benthiques • المجهرية القاعية الطحالب.

Benthic Microalgae • BMA • Cynobacterial Mats • Filamentous Algal Mat

#### Sand

Sand includes any soft-bottom area dominated by fine unconsolidated sediments.

The Sand class describes **soft-bottomed** reef areas where fine unconsolidated **granular material** (finer than coral rubble but coarser than muds) dominate, thickly obscuring any underlying bedrock. Sparse algae, scattered rocks or small, isolated coral heads may also occur in the Sand class, but these features do not exceed 10% of the area [11]. Most reef-associated sands are largely comprised of aragonite (50-80%) and magnesium calcite [27], although the source of the grain (including corals, coralline algae, molluscs, benthic foraminifers and Halimeda, among others) and the grain size will vary and often shows strong cross reef zonation, driven by biogeographic and hydrodynamic factors [27]. Sand class is associated with **Back Reef** zones such as **Inner Reef Flat**, **Back Reef Slope** and **Lagoon** classes in particular, where it can occupy 80% of the area [27].

Sand classes are often subdivided by type (rudstones, grainstones, packstones, mud [27]), biogenic source (e.g. Halimeda grainstones, Foraminiferal Sands [27]), grain size (e.g. fine, medium, coarse sand and gravel [59]) or on which often exhibits strong cross-reef zonation.

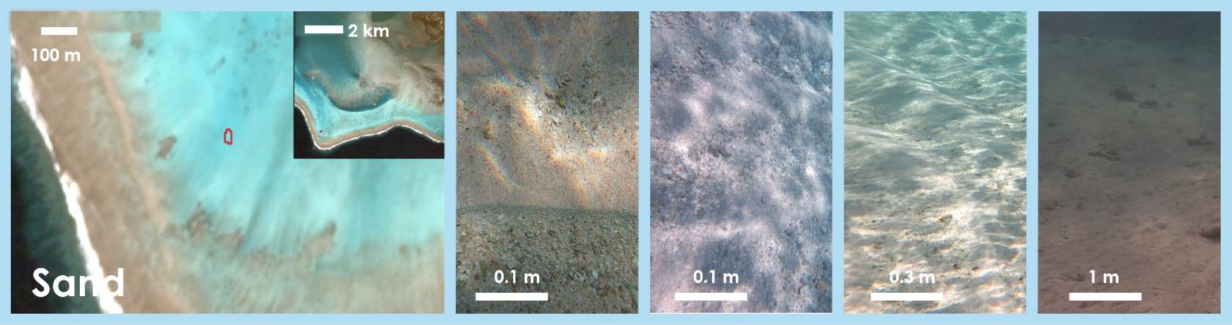


Sand • Pasir • Arena • Le Sable • رمال

Sand dominated • Bioclastic Sand • Sand and sparse algae • Coral Sand • Fine Sediment • Detrital Sand • Sand with Scattered Coral and Rock • Biogenic sand

#### Rubble

Rubble is any habitat featuring loose, rough fragments of broken reef material.

The Rubble class describes any area featuring **loose**, cylindrical to irregularly shaped **fragments** of bedrock or clasts of corals, bivalves and coralline algae [27]. Rubble pieces - while themselves are non-living - can be heavily encrusted by foraminifers, bryozoans or coralline algae, and contain boring organisms. This important habitat often occurs landward of well-developed reef formations in the **Reef Crest**, **Back Reef** or **Reef Flat** zones, and may be associated with some fringing reefs, and also rubble-dominated reef flats [34].

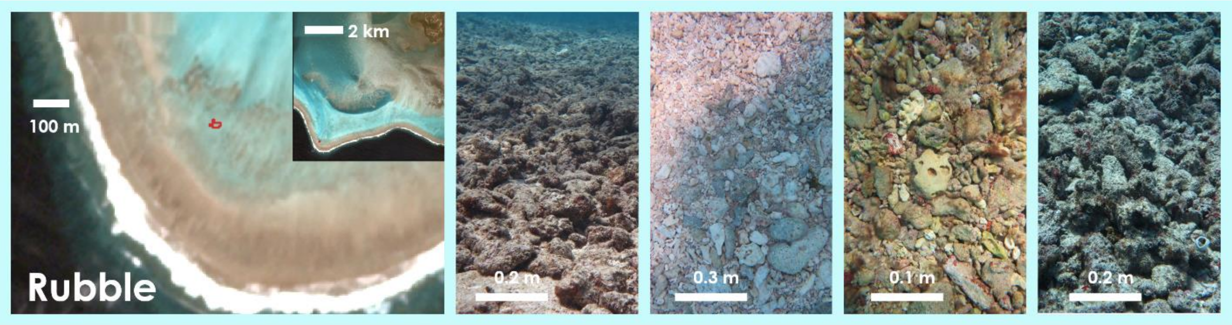


Rubble • Dominan puing/ pecahan karang • Substrato dominado por escombros de arrecifes • Débris dominants • مرجانية بحطام سائد نطاق

Rubble dominated • Skeletal Rubble • Coral Gravel • Coral Rudstone • Reef Rubble • Rhodoliths • Unconsolidated substrate

### Rock

Rock is any exposed hardbottom area with uncommon to scarce corals and fleshy macroalgae.

The Rock class describes any habitat dominated by "*exposed areas of hard bare substratum without visible corallite structure*" [23]. This habitat often has a near horizontal, pavement-like appearance and is usually associated with areas of high energy (e.g. **Reef Crest**) where the cover of living organisms is low (< 10%) - although it may have high coverage of crustose coralline algae. The class encompasses limestone reef matrix, but also underlying non-reefal bedrock and "beach rock" (calcarerite) - areas of coral sand cemented together which are difficult to distinguish by earth observations at present.

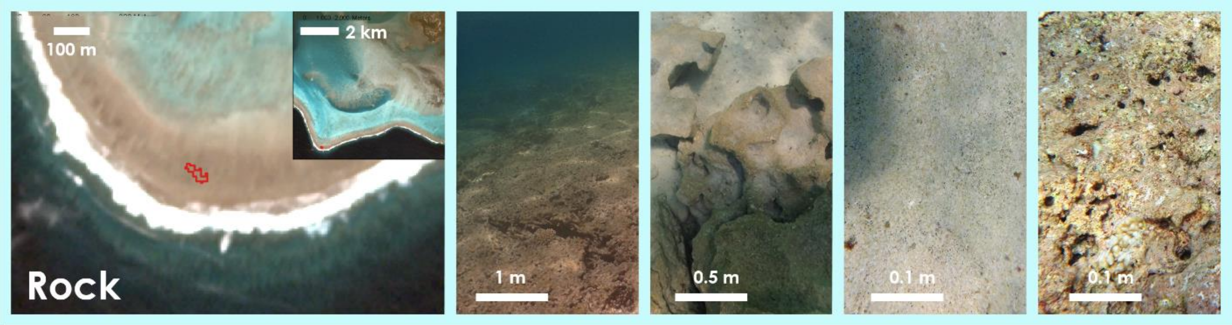


Rock • Batuan • Roca • Roches • صخر

Rock dominated • Bedrock • Pavement • Rock Outcrop • Boulder • Beach Rock • Uncolonised hardbottom

### Attribute table

Attributes of reef zones that help determine Reef Cover class.

| ATTRIBUTE / REEF COVER CLASS | | Depth | | | | | Slope | | | | Exposure | | | Substrate | | | Colour | | | Rugosity | | | Benthic Cover | | | |
|---|---|---|---|---|---|---|---|---|---|---|---|---|---|---|---|---|---|---|---|---|---|---|---|---|---|---|
| | | | | Subtidal | | | | | | | | | | | | | | | | | | | | | | |
| | | Supratidal | Intertidal | Shallow | Medium | Deep | Horizontal | Shallow | Steep | Vertical | Exposed | Average | Sheltered | Hardbottom | Mixed | Soft substrate | Bright | Medium | Darker | Low | Medium | Complex | Coral | Coralline | Sandy | Seagrass |
| Reef Slope | Fore Reef | | | | ✓ | ✓ | | ✓ | ✓ | | ✓ | | | ✓ | | | | ✓ | ✓ | | ✓ | ✓ | ✓ | | | |
| Sheltered Slope | Fore Reef | | | | ✓ | ✓ | | ✓ | ✓ | | | ✓ | | ✓ | | | | ✓ | ✓ | | | ✓ | ✓ | | | |
| Reef Front | Fore Reef | | | ✓ | ✓ | | | ✓ | ✓ | | ✓ | | | ✓ | | | | | ✓ | | | ✓ | ✓ | ✓ | | |
| Sheltered Front | Fore Reef | | | ✓ | ✓ | | | ✓ | ✓ | | | ✓ | | ✓ | | | | | ✓ | | | ✓ | ✓ | | | |
| Wall | Fore Reef | | | | ✓ | ✓ | | | | ✓ | ✓ | | | ✓ | | | | | | | ✓ | ✓ | ✓ | ✓ | | |
| Sheltered Wall | Fore Reef | | | | ✓ | ✓ | | | | ✓ | | ✓ | | ✓ | | | | | | | ✓ | | ✓ | | | |
| Terrace | Fore Reef | | | | ✓ | ✓ | ✓ | | | | ✓ | ✓ | | ✓ | | | | | ✓ | | ✓ | ✓ | ✓ | | ✓ | |
| Reef Crest | Fore Reef | | ✓ | ✓ | ✓ | | ✓ | ✓ | | | ✓ | | | ✓ | | | ✓ | ✓ | | | | | | ✓ | | |
| Outer Reef Flat | Flat | | ✓ | ✓ | | | ✓ | | | | ✓ | ✓ | | ✓ | | | ✓ | | | | ✓ | ✓ | ✓ | ✓ | | |
| Inner Reef Flat | Flat | | | ✓ | ✓ | | ✓ | ✓ | | | | ✓ | | | ✓ | | ✓ | | | | ✓ | | ✓ | | ✓ | |
| Lagoon | Back Reef | | | | | ✓ | ✓ | ✓ | | | | | ✓ | | | ✓ | ✓ | ✓ | | ✓ | | | | | ✓ | |
| Patch Reef | Back Reef | | | ✓ | ✓ | | ✓ | ✓ | | | | | ✓ | ✓ | | | | ✓ | | | | | ✓ | | | |
| Shallow Lagoon | Back Reef | | | ✓ | ✓ | | ✓ | ✓ | | | | | ✓ | | | ✓ | ✓ | | | ✓ | | | | | ✓ | |
| Back Reef Slope | Back Reef | | | | ✓ | ✓ | | ✓ | | | | | ✓ | | ✓ | ✓ | | ✓ | | | ✓ | | ✓ | | ✓ | |
| Plateau | Other | | | | ✓ | ✓ | ✓ | ✓ | | | | ✓ | | ✓ | | | | | ✓ | | ✓ | | ✓ | | | |
| Small Reef | Other | | | | ✓ | ✓ | | | | | ✓ | ✓ | | ✓ | ✓ | | | ✓ | ✓ | | ✓ | | ✓ | | | |
| Reef Island | Other | ✓ | ✓ | | | | | | | | | | | | ✓ | | | | ✓ | | | | | | | |

**Table 2.** Coral reef internal zonation can be linked to an array of gradients in depth, wave action, current, light and sediment across the reef structure. Physical (and biological) attributes can be useful to distinguish between geomorphic zones. Data from different sources e.g. **reflectance** (from satellites, drones and aircraft), **bathymetry** (e.g. LiDAR and Sonar) and **ecological information** (from in-water surveys) all generate attribute datasets that can be used to help determine the 17 Reef Cover classes.

### Crosswalk table

Comparing how Reef Cover classes align with internal coral reef geomorphic classes defined by a selection major global reef classification, mapping and monitoring efforts

| | Marine Ecosystem Classification System (MECS) [49] *classification* | Geomorphological Classification of Reefs (NESP) [60] *classification* | Reef Cover Classification System (GBRMPA) [36] *classification* | Millennium Coral Reef Mapping Project [47] *classification and regional maps* | World Atlas of Coral Reefs [10] *map classes* | Living Oceans Foundation [54] *map classes* | NOAA Biogeography Reef Mapping Program* [61] *map classes* | Allen Coral Atlas [8] *map classes* | Atlantic and Gulf Rapid Reef Assessment [46] *monitoring classes* |
|---|---|---|---|---|---|---|---|---|---|
| **Reef Slope** | ✓ REEF SLOPE (NON-TERRACE) (L5): **REEF EDGE SCARP** (L6). *Orientation: windward ++* | ✓ **SCARP** (L4), steeply sloping (L6) | ✓ **REEF SLOPE** (L1) **(lower reef slope), windward** (L3) | ✓ **SHELF SLOPE** (L4) + multiple L5 categories | ✓ **REEF SLOPE/REEF FRONT** | ✓ **BANK/SHELF ESCARPMENT** (L1) \| **DEEP FORE REEF SLOPE** (L5) | ✓ **BANK/SHELF ESCARPMENT** (L2) | x | ✓ **OUTER FORE REEF** Attribute: *windward* or *seasonally variable, moderate* or *steep* slope |
| **Sheltered Slope** | ✓ REEF SLOPE (NON-TERRACE) (L5): **REEF EDGE SCARP** (L6). *Orientation: leeward ++* | | ✓ **REEF SLOPE** (L1) **(lower reef slope), leeward** (L3) | | | x | | x | ✓ **OUTER FORE REEF** Attribute: *leeward, protected leeward* or *exposed leeward, moderate* or *steep slope* |
| **Reef Front** | ✓ REEF SLOPE (NON-TERRACE) (L5) **SLOPE, SPUR AND GROOVE** *Orientation: windward ++* | ✓ **LEDGE** (L4) **steeply sloping (L6)** | ✓ **REEF SLOPE** (L1) **(upper reef slope) windward** (L3) | ✓ **FOREREEF** (L4) | ✓ **REEF SLOPE/REEF FRONT** | ✓ **FORE REEF** (L1) \| **SHALLOW FORE REEF SLOPE** (L5) | ✓ **FORE REEF** (L2) | ✓ **REEF SLOPE** | ✓ **INNER FORE REEF** Attribute: *windward* or *seasonally variable, moderate* or *steep* slope |
| **Sheltered Front** | ✓ REEF SLOPE (NON-TERRACE) (L5) **SLOPE, SPUR AND GROOVE** *Orientation: leeward ++* | | ✓ **REEF SLOPE** (L1) **(upper reef slope) leeward** (L3) | | | | | ✓ **SHELTERED SLOPE** | ✓ **INNER FORE REEF** Attribute: *leeward, protected leeward* or *exposed leeward, moderate* or *steep slope* |
| **Terrace** | ✓ REEF SLOPE OUTER (L4) \|, **TERRACE (L5)** | ✓ **TERRACE** or **LEDGE** (L4) | ✓ **TERRACES** | ✓ **SHALLOW TERRACE** (L4) | | ✓ ✓ **FORE REEF** (L1) \| **SHALLOW FORE REEF SLOPE** | ✓ **BANK/SHELF** (L2) | x | ✓ **OUTER** or **INNER FORE REEF** Attribute: *flat* |

| | | | | | | | | | |
|---|---|---|---|---|---|---|---|---|---|
| | **PLATFORM – SUBMARINE** (L6) ++ 7 | | | | | or **SAND FLATS** (L5) | | | |
| **Wall** | ✓ REEF SLOPE OUTER (L4) \| REEF SLOPE NON-TERRACE (L5) \| **SUBMARINE CLIFF, SUBMARINE WALL** (L6) *Orientation: windward++* | ✓ **SCARP** (L4), vertical (L6) | x | | | x | ✓ **VERTICAL WALL** (L2) | x | ✓ **OUTER** or **INNER FORE REEF** Attribute: *wall* |
| **Sheltered Wall** | ✓ REEF SLOPE OUTER (L4) \| REEF SLOPE NON-TERRACE (L5) \| **SUBMARINE CLIFF, SUBMARINE WALL** (L6) *Orientation: leeward++* | | x | | | x | | x | ✓ **OUTER** or **INNER FORE REEF** Attribute: *wall, leeward, protected leeward* or *exposed leeward* |
| **Reef Crest** | ✓ REEF TOP (L4) \| **REEF TOP SURFACE FEATURES** (L5) ++ | ✓ **RIDGE** (L4) | ✓ **REEF RIM** (L1) | ✓ **RIM** (L3) / **CREST** (L4) | ✓ **REEF CREST** | ✓ **REEF CREST** (L1) | ✓ **REEF CREST** (L2) | ✓ **REEF CREST** | ✓ **BREAKER ZONE** |
| **Outer Reef Flat** | ✓ REEF TOP (L4) \| REEF FLAT SURFACE FEATURES (L5) \| **REEF PAVEMENT, CORAL BED, COBBLE FIELD** ++ | ✓ **PLATFORM** (L4) rock (L8) | ✓ **OUTER REEF FLAT** (L1) | ✓ **REEF FLAT** (L4) | ✓ **REEF FLAT** | ✓ **BACK REEF** (L1) \| **CORAL REEF AND HARDBOTTOM** (L2) ++ | ✓ **REEF FLAT** (L2) | ✓ **OUTER REEF FLAT** | ✓ **FLAT ZONE** |
| **Inner Reef Flat** | ✓ REEF TOP (L4) \| REEF FLAT SURFACE FEATURES (L5) \| **SAND, RUBBLE, ROCK FLATS (L6)** ++ | ✓ **PLATFORM** (L4) unconsolidated hard (L8) | ✓ **INNER REEF FLAT** (L1) | ✓ **SUBTIDAL REEF FLAT** (L4) | | ✓ **BACK REEF** (L1) \| **UNCONSOLIDATED SEDIMENT** (L2) ++ | | ✓ **INNER REEF FLAT** | ✓ **BACK ZONE** |
| **Back Reef Slope** | ✓ **REEF SLOPE - LAGOON** (L4) + 7 subclasses | ✓ **LEDGE** (L4) **gently sloping** (L6) **unconsolidated soft** (L8) | ✓ **BACK REEF ZONE** (L1) | ✓ **BACK REEF** (L4) | ✓ **BACK REEF** | ✓ **SEDIMENT APRON** (2) | ✓ **BACK REEF** (L2) | ✓ **BACK REEF SLOPE** | ✓ **SUBTIDAL CREST** |
| **Lagoon** | ✓ **LAGOON FLOOR** (L4) ++ 4 subclasses | ✓ **DEPRESSION** (L4) **unconsolidated soft** (L8) | ✓ **LAGOON** (L1) | ✓ **LAGOON** (L3) | ✓ **LAGOON** | ✓ **LAGOON** (L1) \| **UNCONSOLIDATED SEDIMENT** (++4) | ✓ **LAGOON** | ✓ **DEEP LAGOON** | ✓ **LAGOON** |
| **Shallow Lagoon** | ✓ REEF TOP (L4) \| REEF TOP SUBTIDAL FEATURES (L5) \| **MOAT AND DEPRESSION** ++ | ✓ **CHANNEL** (L4) | ✓ **SHALLOW LAGOON** (L1) | ✓ **SHALLOW LAGOON WITH CONSTRUCTIONS / SHALLOW LAGOONAL TERRACE** (L3) | | ✓ **LAGOON** (L1) \| **LAGOON FLOOR** (2) | x but see **CHANNEL** | ✓ **SHALLOW LAGOON** | ✓ **CHANNEL** |

| | | | | | | | | | |
|---|---|---|---|---|---|---|---|---|---|
| **Plateau** | ✓ **SHOAL** + Submerged Atoll, Table Reef, Bank (>20m) ++ | ✓ **BANK** (L4) | ✓ **REEFAL SHOAL** (L1) | ✓ **DEEP TERRACE** (L4) | | x | ✓ **BANK/SHELF** (L2) | ✓ **PLATEAU** | ✓ **PLATFORM** |
| **Patch Reef** | ✓ **LAGOON REEF** (L4) \| ++ 3 subclasses (Patch Reef, Patch Reef + Islet, Reticulate Reef) ++ | ✓ **MOUND** (or **RIDGE**) (L4) | ✓ **PATCH REEF / CORAL HEAD / MICROATOLL** (L2) | ✓ **INTRA-LAGOON PATCH** (L3) | | ✓ LAGOON (L1) \| **PINNACLE REEFS** (++3), **AGGREGATE REEFS** (++ 2), **PATCH REEFS** (++2) (L5) | ✓ **INDIVIDUAL PATCH, AGGREGATED PATCH REEF** (L3) | ✓ **PATCH REEF** | ✓ **SINGLE PATCH, COALESCED PATCH, LINEAR PATCH, RETICULATE PATCH** |
| **Small Reef** | x | ✓ **KNOB** (or **MOUND**) (L4) | x | ✓ **PATCH** (L2) | | x | x but see Aggregate Reef | ✓ **SMALL REEF** | ✓ **SHOAL** |
| **Reef Island** | ✓ **REEF ISLETS** ++ 20 additional classes | x | ✓ **CAY** (L1) | ✓ **LAND ON REEF** (L4) | | ✓ **LAND** and **SHORELINE INTERTIDAL** (L1) ++10 classes | x included in LAND and SHORELINE INTERTIDAL | **x** included in **LAND** | |

✓= same category and class terminology, ✓ = same meaning but different term (synonym), ✓= similar term but slightly different meaning/interpretation (homonym), x = class not considered / not mapped / absent, ++ further subcategories available in classification *same as/similar to **Coastal and Marine Ecological Classification Standard (CMECS)**

**Table 3.** Comparing how some of the major regional to global reef mapping and monitoring efforts (detailed below) classify internal reef structures and how categories relate to Reef Cover classes.

**Marine Ecosystem Benthic Classification (*MECS*)**. MECS, a detailed classification system describing A) pelagic and B) benthic marine features, was developed in 1995 to accompany a broader South Pacific Ecosystem Classification System (SPECS). This classification is applicable outside of the South Pacific and has been successfully modified to other areas (e.g. Caribbean [23]). Within the "Benthic" typology there are **six levels** (L1-L6), with "ecological units" being the lowest level. This classification is highly detailed, Reef Cover classes relate most closely to MECS L4 classes. A major difference from Reef Cover is classes are first split between **Oceanic** Reefs and **Continental Shelf** Reefs, and further by **Reef Type** (e.g. fringing, atoll) before being assigned a geomorphic class.

**Reef Cover Classification System.** Kuchler's 1986 classification system was commissioned by Australia's *Great Barrier Reef Marine Park Authority* (GBRMPA) to standardize geomorphic classification of the Great Barrier Reef for the purposes of mapping using remote sensing data. The classification reviewed decades of commonly used geomorphological nomenclature before designing a typology with **five levels**: **L1 Zones** (36 classes), L2 Features (41), L3 Composition (39), L4 Condition/Morphology (4 levels) and L5 Presence (depths and percent covers, 4 levels). The classification is not strictly hierarchical so a mapping unit could fall into multiple categories. Reef Cover aligns best with L1.

**NESP Geomorphological Classification of Reefs.** The Australian Government's *National Environmental Science Programme* (NESP) geomorphology classification scheme developed for managing continental shelf reefs (biogenic and non-biogenic) using bathymetric (LiDAR) data. The classification draws heavily on the US **Coastal and Marine Ecological Classification Standard** (CMECS) published by the Federal Geographic Data Committee [62]. **Seven categories** include L1 Reef Origin (4 classes), L2 Climatic Region (5), L3 Shelf Zone (4), L4 Geofeature (12), L5 Relief (4), L6 Slope (6), L7 Rugosity (6) and L8 Substrate (4). Reef Cover most closely aligns with L4 to L6.

**Millennium Coral reef Mapping Project (*MCRMP*)**. MCRMP defined a globally standardised coral reef geomorphological typology based on satellite (Landsat 7) imagery. A **five-level** classification system, L1 (Main Division, 2 classes), L2 Nodes (12), L3 Blocks (68), L4 Geomorphological (126) to L5, the most detailed description scheme, a list of 800 different codes and habitats, which are defined through unique combinations of L1 to L4 features (similar to Living Oceans Foundation Approach). Reef Cover aligns most closely with L3/L4 classes, intermediate geomorphological description level that reflects the main structures of a reef complex. Like the MECS and CMECS, MCRMP contains a high level of detail and is clearly split first between **Oceanic** Reefs and **Continental Shelf** Reefs, and further by **Reef Type** (e.g. fringing, atoll, barrier), but unlike the other schemes is strictly morphological and does not include benthic cover classes.

**World Atlas of Coral Reefs**. The *United Nations Environment Program World Conservation Monitoring Centre* (UNEP-WCMC) began its global coral reef mapping work in 1994, eventually publishing the *World Atlas of Coral Reefs*: the most comprehensive global extent map of coral reefs currently in existence in 2000 [10]. The map was derived from multiple sources, primarily maps supplied by the MCRMP at L3 (80% of the map), but also including existing US Defence Force navigational charts, and digitised regional topographic and specialised scientific reef maps (some as old as 150 years), collated as part of the *Corals of the World* volumes. Because of the different data sources a map a uniform classification was not possible, but the *World Atlas of Coral Reefs* suggests key reef classes in the accompanying book and dataset.

**Living Oceans Foundation *Global Reef Expedition***. A hierarchical geomorphic classification scheme was developed for the Global Reef Expedition for the purpose of supporting mapping of 65,000 sq km of the world's remotest reefs from 11 countries [54], using combined satellite, bathymetric and ecological data. Hierarchical classification that accompanies the maps have **five levels** that include benthic as well as geomorphic components: L1 Zones (8 classes, based purely on bathymetry), L2 Major Geomorphological Structure (3), L3 Detailed Geomorphological Structure Map (11) together form the hierarchical classification, then L4 Biological Cover (from field surveys) and L5 Aggregate Map Class (36 classes). The 36 classes include 6 shoreline intertidal class, 4 land classes. Reef Cover classes align best with L1-L5.

**NOAA Biogeography Reef Mapping Program**. The US *National Oceanic and Atmospheric Administration* (NOAA)'s National Ocean Service developed a series of hierarchical geomorphic habitat classification schemes to support a reef habitat mapping effort, aimed at mapping 43,000 sq km of coral reef across the US territories. Classifications had some regional variations with five classifications developed for the seven mapped regions [61], but largely followed the same format, with **five levels**: L1 Geographic Zones (~14 classes), L2 Geomorphological Structures (~12), L3 Biological Cover types (~7), L4 Coral Cover categories (4 classes: 10-50%, 50-90%, 90-100% or unknown) and L5 Percent Hardcover categories (4) [11]. Reef Cover most closely aligns with L1, but misses key classes SALT POND, SHORELINE INTERTIDAL, DREDGED, CHANNEL and REEF HOLE. This classification also feeds into the US **Coastal and Marine Ecological Classification Standard (*CMECS*)** published by the Federal Geographic Data Committee [62], which describes 10 similar reef classes to level L2 (Back Reef, Bank/Shelf, Bank/Shelf Escarpment, Fore Reef, Lagoon, Reef Crest, Reef Flat, Vertical Wall, Shoreline/Intertidal and an extra class, Ridges and Swales).

**Allen Coral Atlas**. The Allen Coral Atlas developed by Vulcan has 18 map classes largely derived from a classification developed for reef mapping based on hierarchical rulesets based on environmental attributes for object-based image classification, which has four levels L1 Reef Type, L2 Reef Type, L3 Geomorphic (12 classes), and L4 Benthic (6) [53]. Reef Cover is similar to L3 Global Geomorphic Classes but shallow map depth means Reef Slope classes are missing and additional class TERRESTRIAL REEF FLAT also exists.

**Atlantic and Gulf Rapid Reef Assessment (*AGRRA*).** AGRRA is a Caribbean-specific classification developed to support field monitoring purposes and guide surveywork. The "Reef Type" classification is comprised of four-levels: L1 Location (6 classes), L2 Reef Type (15), L3 G**eomorphic Zone (9)**, L4 Habitats (9) which can be combined with 14 **Attribute Classes** related to exposure, incline and relief) to describe the reef type being monitored. Reef Cover aligns well with the L3 classes, but is missing "FLANK".

### Reef Cover Typology

#### Example decision tree for classification of Reef Cover

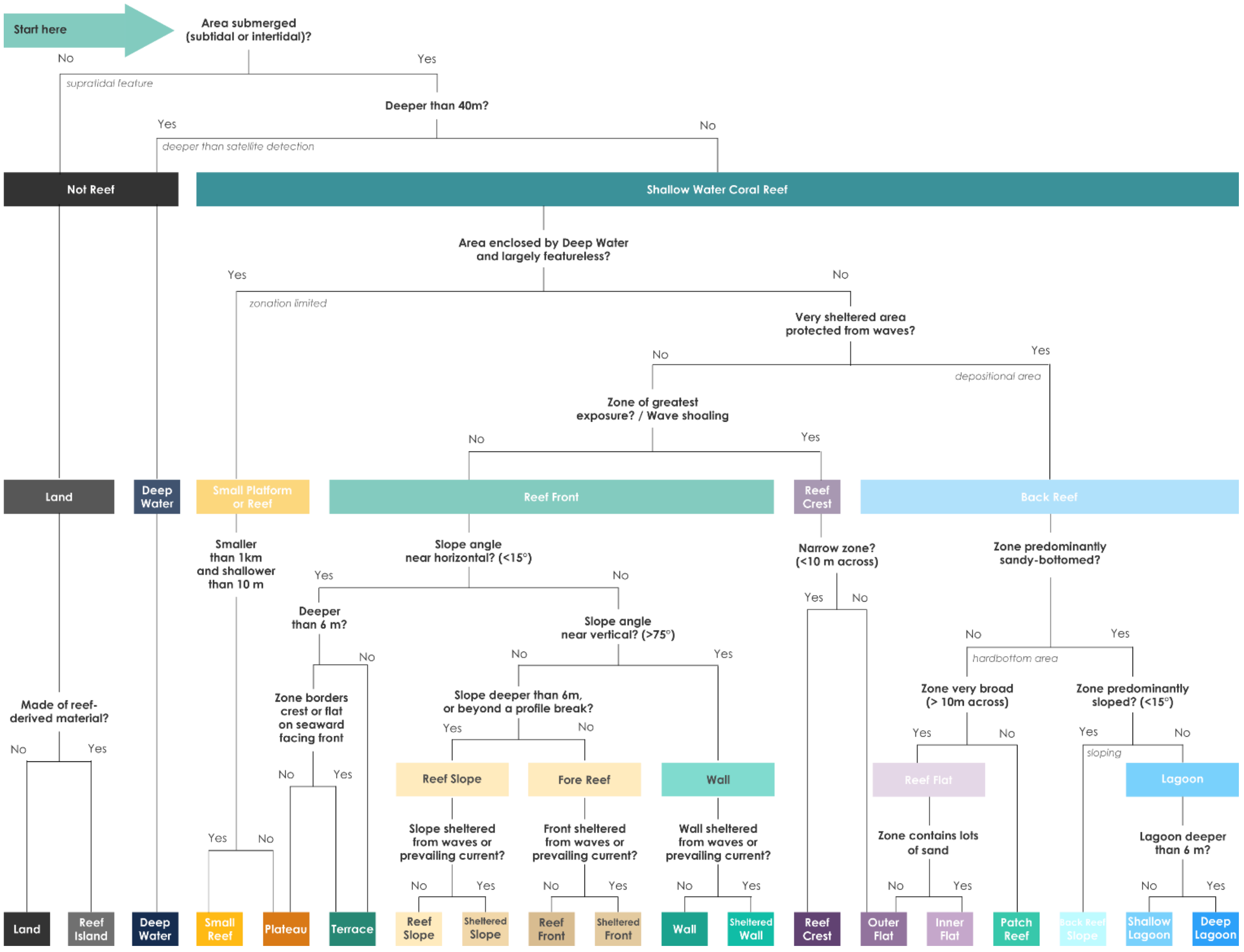


**Figure 2**. Example decision tree for Reef Cover classification. The decision tree is based on attribute information (Table 2) that would typically be available at the global scale from either remotely sensed information on in-water ecological information, and related to the physical attributes (depth, slope angle and exposure), colour and texture, and spatial relationships.

### Glossary

| | |
|---|---|
| **Fore Reef** | **Fore Reef** is the classical description for one of the three major geomorphic elements of a coral reef, along with **Reef Flat** and **Back Reef** [12, 23, 27, 30, 42, 54]. Characteristics include being **subtidal**, **seaward-facing** (opening up into open sea or ocean, facing away from land or lagoons) [12, 13] and **sloping** (to various degrees: atolls have straighter, steeper profiles, barriers tend to be subdivided into bumps or "terraces", and fringing reef profiles are gentler [9, 14-16]) part of the reef.<br><br>**Fore Reefs** are often further divided into geomorphic sub-zones, primarily based on a) **depth** and natural breaks in slope (e.g. **Fore Reef** / **Reef Front** / Shelf Break) b) slope **angle** (e.g., **Wall** / **Terrace** / Spur and Groove) and c) the level of **exposure** (e.g. Leeward or Sheltered / Windward or Exposed) - three attributes directly linked to their three main defining characteristics (in green). This is not surprising, given how reef slope communities experience large gradients in light (related to depth) and exposure [15]. |
| **Reef Flat** | Boasting the **broadest extent** of any reef zone, Reef Flat is a **level** (horizontal) and **shallow** (generally <3 m deep) carbonate platform extending inwards from the **Reef Front** with a gentle downward gradient in slope. Often from 500 – 1000 m across but in the case of some Pacific atolls sometimes several kilometres across. The Reef Flat represents an area where upward reef growth has been constrained by the sea surface, and are sometimes but not always intertidal. Characterised by **strong zonation** driven by water movement, which can result in strongly banded **Outer** and **Inner Reef Flat** sub-zones. |
| **Back Reef** | Back Reef is a widely-used term and broadly agreed to be the third of the three classical geomorphic zones (which include **Fore Reef** and **Reef Flat**). In the literature it has varied definitions, probably due to biogeographic variability, and adoption by ecologists of what was originally a geological term to describe the detritus-dominated slowly accumulating zone (as opposed to the rapidly accumulating coral dominated reef front) [63]. Sometimes, it refers purely to the transitional landward sloping area that links the **Reef Flat** and the **Lagoon** (often also called a **Back Reef Slope**) [10]. Other times it is defined positionally in relation to the **Reef Crest**, as any reef feature found landward of the crest. In this definition, it can encompass a) **Reef Flat**, b) **Lagoon** *and* c) **Back Reef Slope**. Alternatively, its defined more around its functional role, around the fact that it is a largely depositional environment, receiving debris swept landwards from the **Reef Crest** and **Reef Front** – from rubble strewn across **Reef Flats**, to sand sheets and mud deposits into **Lagoons** [12]. In these cases, it tends to include **Lagoon** and **Back Reef Slope** but not the more productive **Reef Flat** with up to 80% sediment produced by the windward reef flats are transported into this back reef areas [27]. From this geological perspective, the Back Reef is also seen as transitional stage in a reefs life, the end point will be infilling [27]. Features are that it **is interior-reef zone** (either landward of the crest or flat) and that it is characteristically a **sheltered** and **depositional environment** – so will be sand-dominated. |

| | |
|---|---|
| **Deep Water** | Deep Water is any off-reef **pelagic body of water** beyond remote sensing depth detection limits (generally around 20-30 m) or outside of the scope of mapping exercise, and could include neritic (water on continental shelf, seas) or oceanic waters and epipelagic to hadalpelagic zones. |
| **Land** | Land is any **supratidal** area beyond the littoral zone with a non-biogenic origin. Land is a commonly adopted class in almost all existing reef cover classifications, given the importance of reef positioning relative to land to determine reef type (e.g. **Atoll**, **Barrier** and **Fringing Reef**), and the large influence of land on reefs. Several classifications further subdivide the land category e.g. Living Oceans Foundation subdivide into "man-made" (e.g. roads, infrastructure), "vegetated" and "non-vegetated" [54]. |
| **Atoll** | An Atoll is a **solitary ring-to-irregularly-shaped** coral reef rising up out of **Deep Water** and encircling or partially **enclosing a wide central Lagoon**.<br><br>The main distinguishing characteristic of an Atoll is an **enclosed (or partially enclosed) central lagoon**; the second universal feature is a "**reef rim**" (**Reef Crest**/**Reef Flat**) generally less than 10 m deep [41]. The mapped ratio of lagoon to rim is high [39], the rim is generally elongated "ribbon-like", narrow in respect to the lagoon and generally steep-sided on the seaward side [39]. Reef flats of atolls in the Pacific tend to be 100-1000 m wide (average 500 m) [41]. Most definitions describe the rim as "**ring-shaped**" [64] (although they can be in various shapes from "commas" to "figure eights" [65]. Other defining features often described are a) the relative **isolation** of the reef (variously described as "stand apart" [66], "detached" or "solitary" – which is why Maldivian farus are sub-atoll features); b) **variable continuousness of the rim** (from "almost enclosed" [67] to "more or less continuous" [66]) affecting the degree to which the **Lagoon** is connected to the surrounding ocean [41]; c) **low-lying** "slightly submerged" [41], d) **associated with low-lying islands and islets** ("slightly emerged reefs or small sand cays" (Shepard, 1948)) although recently proposed to have islands should be <5% of the whole rim area [65] – North Male has more than 50 islands, and d) **arising from deep water** "typically found in oceanic locations" [67], and "arising from deep water" [41] "far from land in the open ocean" [68]. |
| **Barrier Reef** | A Barrier Reef is an extensive **linear reef complex** running **parallel to the coastline**, with one side neighbouring **Deep Water**, and **separated from the shore** by some distance by deep and broad body of water (a **Lagoon**).<br><br>The critical defining feature of a Barrier Reef is the presence of a **Lagoon** separating the Reef Front from the shoreline. In terms of depth, **Lagoons** can be up to 50-80 meters [69], most agree a depth of >10 m to prevent barrier being a **Fringing Reef**, while in the Caribbean AGRRA use a threshold of 300 m width to distinguish a Barrier [46]. Additional features are that, unlike circular **Atolls**, they are strongly **asymmetric in plan and section**, being steeper on the ocean side (often dropping abruptly to 1000+ meters) compared to the interior, which grades off gently to the interior with a sediment wedge dotted with small reef patches, pinnacles and coral heads [67]. |

| | |
|---|---|
| **Fringing Reef** | Fringing Reefs are reefs connected to or closely associated with the mainland or island shoreline, consisting of a **Fore Reef**, **Reef Crest** and **Back Reef**, and where the **Back Reef** water depth is too shallow (generally <10 m) to be a barrier reef.<br><br>Fringing reefs [30] are the most prevalent class of coral reefs globally [42], and often described as the "simplest" form of reef and a potential geologic "precursor" to other reefs [39]. A primary characteristic is its **close association with the mainland or island shoreline** [14, 70], with reefs frequently being shore-attached [30], or "within easy swimming distance" [46]. The second main feature to distinguish from a Barrier Reef is a **lack of a significant break (Lagoon) between the shoreline and the reef.** While Fringing Reefs can have a narrow **Shallow Lagoon**, generally both the depth and width are limited. "A water depth over the back reef of less than 10 m is usually the principal criterion used to define a reef as a fringing reef" but "no quantitative standards are provided" for the proposed threshold on lagoon width [70]. Other features of Fringing Reefs are their relatively thin veneer (compared to **Atolls** and **Barriers** that can form platforms several hundred meters thick) since their upward growth is limited as they develop from the shallow shelving coastline. They are also narrow: **Reef Flats** are typically 50 m to 1500 m across [38], narrow belt (1-2km max), with their dimensions a function of the underlying slope [30]. Unlike **Atolls** and **Barriers**, Fringin Reef development is not so dependent on a reefal foundation and different types of shelving land determine different origins. For example, they often form a semi-circular fan around headlands [38], but are continuous along limestone coasts. Lack of dependence on a hard reefal foundation means Fringing Reefs are less likely to drown, and their development is driven more by local ecological factors (like rivers and sediment) and less by eustatic and tectonic factors [39]. Fringing reefs are subject to sedimentation, freshwater dilution and among the poorest developed reefs, corals often growth inhibited [39]. |
| **Platform Reef** | Platform Reef is an **irregularly shaped**, **broad flat** tabulate reef arising from relatively shallow water on a **continental shelf**.<br><br>Platform Reefs are a variable and complex group of reefs found on continental shelves, defined as any "irregular islandless reefs which rise from the shallower parts of a continental shelf" [39]. The primary feature is these reefs is their association with the relatively shallow water (<50 m) of the shelf or continental slope [39]. Other features include being **irregularly shaped**, unlike their **Atoll** (generally ovoid), **Barrier** (elongate) and **Fringing** (narrow shelf hugging) counterparts. This is due to a combination of reef morphology being determined by antecedent platform, but also because theoretically platform reefs can extend outwards in any direction, whereby growth of **Atolls** (upward growth) and **Barriers** / **Fringing Reefs** (outward growth) tends to be mainly unidirectional. Irregular size and extent also means platform reefs have varied internal morphology: larger platforms often have deep lagoons or networks of lagoons while others are flat and table-like. Platform Reefs on the Great Barrier Reef are often flat and broad, but in other ocean basins can be deeper or less flat. Finally, most definitions of platform classify heavily according to size. While platform reefs are described as variable "ranging from a few hundred metres to many kilometres across" some definitions have suggested a threshold of >1 km across, with smaller platforms <1 km across (variously described as patch reefs [69]. |

### References


1. Sokal, R.R., *Classification: Purposes, Principles, Progress, Prospects.* Science, 1974. **185**(4157): p. 1115-1123.

2. Purkis, S.J., *Remote sensing tropical coral reefs: The view from above.* Annual Review of Marine Science, 2018. **10**(1): p. 149-168.

3. Stoddart, D.R., R. McLean, and D. Hopley, *Geomorphology of reef islands, Northern Great Barrier Reef.* 1978.

4. Zann, M., E. Kenna, and M. Ronan, *Queensland Intertidal and Subtidal Ecosystem Classification Scheme: Introduction and implementation of Intertidal and Subtidal Ecosystem Classification*, Q.G. Department of Environment and Heritage Protection, Editor. 2017: Brisbane, Australia. p. 77.

5. Di Gregorio, A., *Land Cover Classification System: Software version (3)*. 2016, Food and Agriculture Organisation of the United Nations: Rome.

6. UNEP-WCMC, et al., *Global distribution of warm-water coral reefs, compiled from multiple sources (listed in "Coral_Source.mdb"), and including IMaRS-USF and IRD (2005), IMaRS-USF (2005) and Spalding et al. (2001).* . 2010: Cambridge (UK): UNEP World Conservation Monitoring Centre. URL: data.unep-wcmc.org/datasets/13.

7. Hamylton, S., S. Andréfouët, and T. Spencer, *Comparing the information content of coral reef geomorphological and biological habitat maps, Amirantes Archipelago (Seychelles), Western Indian Ocean.* Estuarine, Coastal and Shelf Science, 2012. **111**: p. 151-156.

8. Lyons, M.B., et al., *Mapping the world's coral reefs using a global multiscale earth observation framework.* Remote Sensing in Ecology and Conservation, 2020. **n/a**(n/a).

9. Kuchler, D., *Reef cover and zonation classification system for use with remotely sensed Great Barrier Reef data: uder guide and handbook*, ed. G.B.R.M.P.A.-t.m.n. GBRMPA-TM-8. 1987, Townsville, Australia.

10. Spalding, M., C. Ravilious, and E. Green, *World Atlas of Coral Reefs*, ed. P.a.t.U.W.C.M. Centre. 2001, Berkeley (California, USA): University of California Press. 436 pp.

11. Zitello, A.G., et al., *Shallow-Water Benthic Habitats of St. John, U.S. Virgin Islands*, N.T.M.N.N. 96, Editor. 2009: Silver Spring, MD. 53 pp. .

12. Collins, L.B., *Reef Structure*, in *Encyclopedia of Modern Coral Reefs: Structure, Form and Process*, D. Hopley, Editor. 2011, Springer Netherlands: Dordrecht. p. 896-902.

13. Cabioch, G., *Forereef/Reef Front*, in *Encyclopedia of Modern Coral Reefs: Structure, Form and Process*, D. Hopley, Editor. 2011, Springer Netherlands: Dordrecht. p. 422-423.

14. Montaggioni, L.F. and C.J. Braithwaite, *Quaternary Coral Reef Systems: history, development processes and controlling factors*. Developments in Marine Geology. 2009.

15. Done, T.J., *Coral zonation: Its nature and significance,*, in *Perspectives on Coral Reefs*, D.J. Barnes, Editor. 1983, Australian Institute of Marine Science: Townsville. p. 107-147.

16. Maxwell, W.G.H., *Atlas of the Great Barrier Reef*. 1968.

17. Bongaerts, P., et al., *Deep reefs are not universal refuges: Reseeding potential varies among coral species.* Sci Adv, 2017. **3**(2): p. e1602373.

18. Blanchon, P., *Geomorphic Zonation*, in *Encyclopedia of Modern Coral Reefs: Structure, Form and Process*, D. Hopley, Editor. 2011, Springer Netherlands: Dordrecht. p. 469-486.

19. Sheppard, C., *Coral population on reef slopes and their major controls.* Marine Ecology Progress Series, 1982. **7**: p. 83-115.

20. Bridge, T., et al., *Predicting the location and spatial extent of submerged coral reef habitat in the Great Barrier Reef world heritage area, Australia.* PloS one, 2012. **7**(10): p. e48203-e48203.

21. Bongaerts, P., et al., *Assessing the 'deep reef refugia' hypothesis: focus on Caribbean reefs.* Coral Reefs, 2010. **29**(2): p. 309-327.

22. Eakin, C.M., H.P.A. Sweatman, and R.E. Brainard, *The 2014–2017 global-scale coral bleaching event: insights and impacts.* Coral Reefs, 2019. **38**(4): p. 539-545.

23. Mumby, P.J. and A.R. Harborne, *Development of a systematic classification scheme of marine habitats to facilitate regional management and mapping of Caribbean coral reefs.* Biological Conservation, 1999. **88**(2): p. 155-163.

24. Geister, J. *The influence of wave exposure on the ecological zonation of Caribbean coral reefs,*. in *Proceedings of Third International Coral Reef Symposium*. 1977. Miami, Florida.: Rosenstiel School of Marine and Atmospheric Science.

25. Done, T.J., *Patterns in the distribution of coral communities across the central Great Barrier Reef.* Coral Reefs, 1982. **1**(2): p. 95-107.

26. Stoddart, D.R., *Ecology and morphology of recent coral reefs.* Biological Reviews, 1969. **44**(4): p. 433-498.

27. Montaggioni, L.F., *History of Indo-Pacific coral reef systems since the last glaciation: Development patterns and controlling factors.* Earth-Science Reviews, 2005. **71**(1): p. 1-75.

28. Bruckner A, R.G., Riegl B, Purkis SJ, Williams A, Renaud P, *Khaled bin Sultan Living Oceans Foundation atlas of Saudi Arabian Red Sea marine habitats*. ISBN-978-0-9835-611- 1-8. 273 pp. 2011.

29. Edinger, E.N. and M.J. Risk, *Reef classification by coral morphology predicts coral reef conservation value.* Biological Conservation, 2000. **92**(1): p. 1-13.

30. Kennedy, D.M. and C.D. Woodroffe, *Fringing reef growth and morphology: a review.* Earth-Science Reviews, 2002. **57**(3): p. 255-277.

31. Ferrario, F., et al., *The effectiveness of coral reefs for coastal hazard risk reduction and adaptation.* Nature Communications, 2014. **5**: p. 3794.

32. Lowe, R.J., et al., *Spectral wave dissipation over a barrier reef.* Journal of Geophysical Research: Oceans, 2005. **110**(C4).

33. Harris, D.L., et al., *Coral reef structural complexity provides important coastal protection from waves under rising sea levels.* Science Advances, 2018. **4**(2): p. eaao4350.

34. Thornborough, K.J. and P.J. Davies, *Reef Flats*, in *Encyclopedia of Modern Coral Reefs: Structure, Form and Process*, D. Hopley, Editor. 2011, Springer Netherlands: Dordrecht. p. 869-876.

35. Hopley, D., *Morphological classifications of shelf reefs: a critique with special reference to the Great Barrier Reef*, in *Perspectives on Coral Reefs*, D.J. Barnes, Editor. 1983, Australian Institute of Marine Science.

36. Kuchler, D., *Reef cover and zonation classification system for use with remotely sensed Great Barrier Reef data*, in *Technical memorandum TM-7*. 1986: Townsville.

37. Bellwood, D.R., et al., *The role of the reef flat in coral reef trophodynamics: Past, present, and future.* Ecology and Evolution, 2018. **8**(8): p. 4108-4119.

38. Fairbridge, R.W., *Fringing Reef*, in *Geomorphology. Encylopedia of Earth Science*. 1968, Springer Berlin Heidelberg: Berlin, Heidelberg. p. 366-369.

39. Fairbridge, R.W., *Recent and Pleistocene coral reefs of Australia.* The Journal of Geology, 1950. **58**(4): p. 330-401.

40. Dickenson, W.R., *Pacific Atoll living: how long already and until when?* GSA Today, 2009.

41. Woodroffe, C.D. and N. Biribo, *Atolls*, in *Encylopedia of Modern Coral Reefs*. 2011, Springer: The Netherlands.

42. Hopley, D., S.G. Smithers, and P.K. E, *The Geomorphology of the Great Barrier Reef: development, diversity and change*. 2007, Cambridge: Cambridge University Press.

43. Aswani, S. and I. Vaccaro, *Lagoon Ecology and Social Strategies: Habitat Diversity and Ethnobiology.* Human Ecology, 2008. **36**(3): p. 325-341.

44. Fairbridge, R.W., *Lagoon (coral-reef type)*, in *Geomorphology. Encyclopedia of Earth Science.* 1968, Springer Berlin Heidelberg: Berlin, Heidelberg. p. 594-598.

45. Guilcher, A. *A heretofore neglected type of coral reef: the Ridge Reef. Morphology and Origin*. in *Proceedings of the 6th International Coral Reef Symposium*. 1988. Australia.

46. Lang, J. and K. Marks. *AGRRA (Atlantic and Gulf Rapid Reef Assessment): Reef Terms*. 2018.

47. Andréfouët, S., et al. *Global assessment of modern coral reef extent and diversity for regional science and management applications: a view from space*. in *Proceedings of 10th International Coral Reef Symposium*. 2006. Okinawa, Japan. June 28-July 2, 2004. 1732-1745 pp.

48. McLean, R. and P. Kench, *Destruction or persistence of coral atoll islands in the face of 20th and 21st century sea-level rise?* Wiley Interdisciplinary Reviews: Climate Change, 2015. **6**(5): p. 445-463.

49. Holthus, P.F. and J.E. Maragos, *Chapter 13: Marine ecosystem classification for the Tropical Island Pacific*, in *Marine and Coastal Biodiversity in the Tropical Island Pacific Region*. 1995: Honolulu, Hawaii. p. 239-278.

50. Hamylton, S., *Will coral islands maintain their growth over the next century? A deterministic model of sediment availability at lady elliot island, great barrier reef.* PLOS ONE, 2014. **9**(4): p. e94067.

51. Hamylton, S.M., *Mapping coral reef environments:A review of historical methods, recent advances and future opportunities.* Progress in Physical Geography: Earth and Environment, 2017. **41**(6): p. 803-833.

52. Goodman, J.A., S. Purkis, and S. Phinn, *Coral Reef Remote Sensing: a Guide for Mapping, Monitoring and Management*. 2013: Springer Netherlands.

53. Roelfsema, C., et al., *Mapping coral reefs at reef to reef-system scales, 10s–1000s km2, using object-based image analysis.* International Journal of Remote Sensing, 2013. **34**(18): p. 6367-6388.

54. Purkis, S.J., et al., *High-resolution habitat and bathymetry maps for 65,000 sq. km of Earth's remotest coral reefs.* Coral Reefs, 2019.

55. Hedley, J.D., et al., *Spectral unmixing of coral reef benthos under ideal conditions.* Coral Reefs, 2004. **23**(1): p. 60-73.

56. Roelfsema, C., S. Phinn, and W. Dennison, *Spatial distribution of benthic microalgae on coral reefs determined by remote sensing.* Coral Reefs, 2002. **21**(3): p. 264-274.

57. Pinckney, J.L., *A mini-review of the contribution of benthic microalgae to the ecology of the continental shelf in the south atlantic bight.* Estuaries and Coasts, 2018. **41**(7): p. 2070-2078.

58. Madin, E.M.P., et al., *Marine reserves shape seascapes on scales visible from space.* Proceedings of the Royal Society B: Biological Sciences, 2019. **286**(1901): p. 20190053.

59. Davies, P.J., *Reef Growth*, in *Perspectives on coral reefs*, D.J. Barnes, Editor. 1983, Australian Institute of Marine Science: Townsville.

60. Nichol, S.L., et al., *Geomorphological classification of reefs*, in *Report to the National Environmental Science Program, Marine Biodiversity Hub.* 2016: Geoscience Australia. p. 27.

61. Monaco, M.E., et al., *National Summary of NOAA's Shallow-water Benthic Habitat Mapping of U.S. Coral Reef Ecosystems*, in *NOAA Technical Memorandum NOS NCCOS 122.* 2012, Prepared by the NCCOS Center for Coastal Monitoring and Assessment Biogeography Branch: Silver Spring, MD.

62. Committee, F.G.D., *Coastal and Marine Ecological Classification Standard.* 2012.

63. Montaggioni, L.F., *Anatomy of shallow-water carbonate bodies : lessons from the Recent coral reef record.* 2001.

64. Allaby, *Dictionary of Geology and Earth Sciences (4th edition)*. 2013, Oxford University Press.

65. Goldberg, W.M., *Atolls of the world: revisiting the original checklist.* Atoll Research Bulletin, 2016. **610**.

66. Weins, H.J., *Atoll environment and ecology.* 1962.

67. Darwin, C., *The Structure and Distribution of Coral Reefs. Being the first part of the geology of the voyage of the Beagle, under the command of Capt. Fitzroy, R.N. during the years 1832 to 1836*. 1843, London: Smith Elder & Co.

68. Moberg, F. and C. Folke, *Ecological goods and services of coral reef ecosystems.* Ecological Economics, 1999. **29**(2): p. 215-233.

69. Fairbridge, R.W., *Coral reefs—morphology and theories*, in *Geomorphology*. 1968, Springer Berlin Heidelberg: Berlin, Heidelberg. p. 186-197.

70. Smithers, S.G., *Fringing Reefs*, in *Encyclopedia of Modern Coral Reefs*. 2011, Springer.

### Acknowledgments

This work was initiated and funded primarily through **Paul Allen Philanthropies** and **Vulcan Inc.** as part of the **Allen Coral Atlas**. We acknowledge the late Paul Allen and Ruth Gates for their fundamental vision and drive to enable us to work together on this critical reef mapping problem. Project partners providing financial, service and personnel include: **Planet Inc.**, **National Geographic**, **University of Queensland**, **Arizona State University**, and **University of Hawai'i**. Significant support has also been provided by **Google Inc.**, **Great Barrier Reef Foundation**, and **Trimble (Ecognition)**. Contributors to establishing and running the project include: **Vulcan Inc.** [James Deutsch, Lauren Kickam, Paulina Gerstner, Charlie Whiton, Kirk Larsen, Sarah Frias Torres, Kyle Rice, Janet Greenlee]; **Planet Inc.** [Andrew Zolli, Trevor McDonald, Joe Mascaro, Joe Kington]; **University of Queensland** [Chris Roelfsema, Stuart Phinn, Emma Kennedy, Mitch Lyons, Nicholas Murray, Doddy Yuwono, Dan Harris, Eva Kovacs, Rodney Borrego, Meredith Roe, Jeremy Wolff, Katherine Markey, Alexandra Ordonez, Chantal Say, Paul Tudman]; **Arizona State University** [Greg Asner, Dave Knapp, Jiwei Li, Yaping Xu, Nick Fabina, Heather D'Angelo]; and **National Geographic** [Helen Fox, Brianna Bambic, Brian Free, Zoe Lieb] and **Great Barrier Reef Foundation** [Petra Lundgren, Kirsty Bevan].

This work was also partly supported by the **Great Barrier Reef Marine Park Authority**, and by an **Australian Research Council** Discovery Early Career Researcher Award (DE190100101) to **Nicholas Murray**.

We would also like to thank Coral Reef Classification workshop participants **Emma Kennedy**, **Chris Roelfsema**, **Eva Kovacs**, **Daniel Harris**, **Mitchell Lyons**, **Greg Webb**, **Doddy Yowono** and **Atefeh Sansoleimani** (University of Queensland), **Sarah Hamylton** (University of Wollongong), **Javier Leon** (Southern Cross University) and **Stephanie Duce** (James Cook University).

We are grateful to **Serge Andréfouet**, **Hiroya Yamano**, **Sam Purkis**, **Mark Spalding** and **Sarah Hamylton** for providing expert guidance and helpful feedback on an early manuscript draft, to **Mike Ronan** and **Maria Zann** from the Queensland Government for guidance with classification, and to **Doddy Yuwono**, **Rodney Borrego-Acevedo**, **Jeremy Wolff**, **Farid Al Abdali** and **Abdullah Al Kindi** for help translating technical terms.

"As long as we work together with both urgency and determination, there are no limits to what we can achieve."

**Paul G Allen, 1953-2018**

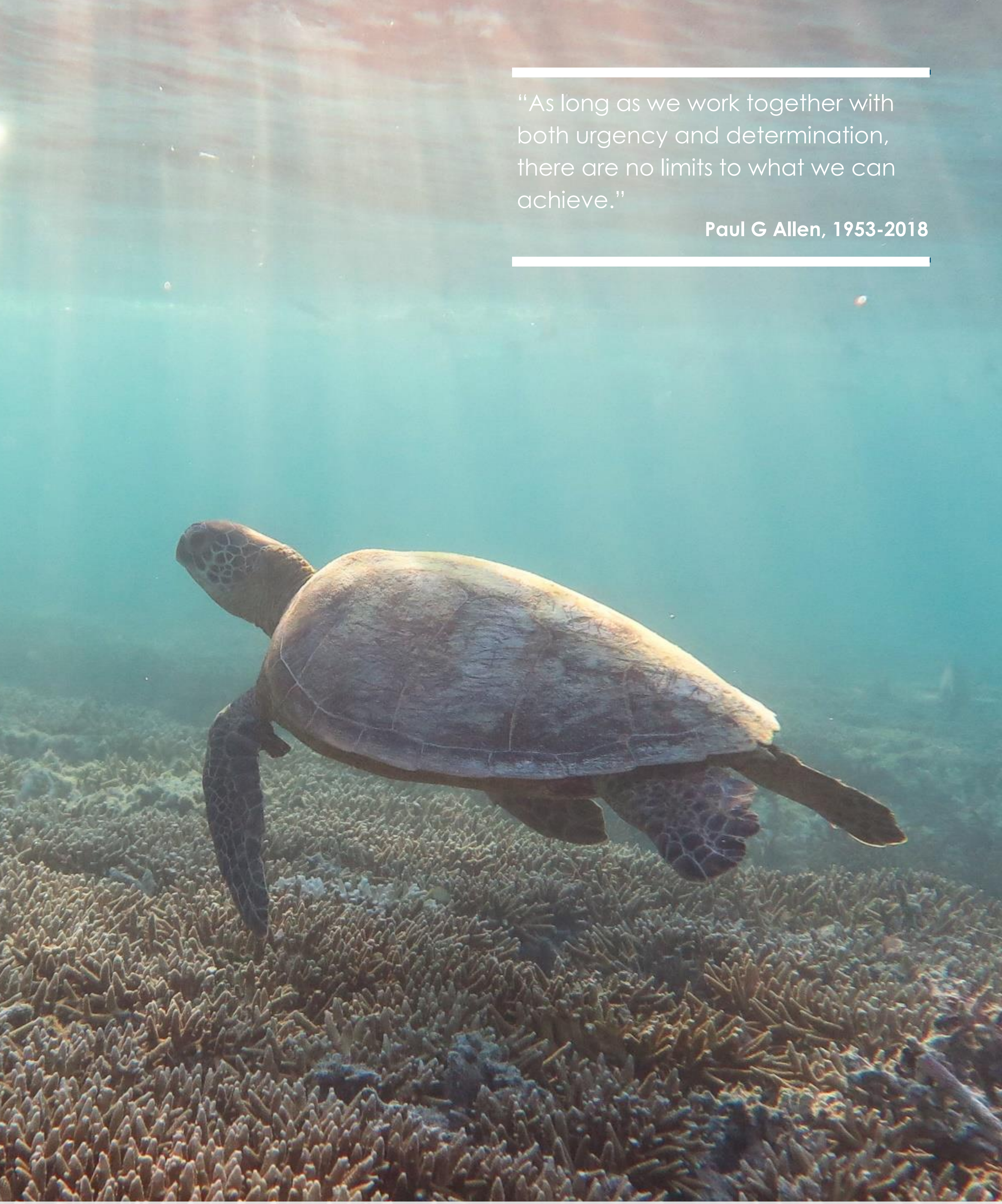

Remote Sensing Research Centre | The University of Queensland Australia | Australian Government Great Barrier Reef Marine Park Authority | planet. | ASU Center for Global Discovery and Conservation Science Arizona State University | National Geographic | Vulcan
